# Sequential Transitions of Male Sexual Behaviors Driven by Dual Acetylcholine-Dopamine Dynamics

**DOI:** 10.1101/2023.12.21.572798

**Authors:** Ai Miyasaka, Takeshi Kanda, Naoki Nonaka, Yuka Terakoshi, Yoan Cherasse, Yukiko Ishikawa, Yulong Li, Hotaka Takizawa, Jun Seita, Masashi Yanagisawa, Takeshi Sakurai, Katsuyasu Sakurai, Qinghua Liu

## Abstract

The neural mechanisms regulating sequential transitions of male sexual behaviors, such as mounting, intromission, and ejaculation, in the brain remain unclear. Here, we report that dopamine (DA) and acetylcholine (ACh) dynamics in the ventral shell of the nucleus accumbens (vsNAc) closely aligns with serial transitions of sexual behaviors in male mice. During intromission, the vsNAc exhibits dual ACh-DA rhythms generated by reciprocal regulation between ACh and DA signaling via nicotinic acetylcholine (nAChR) and dopamine D2 (D2R) receptors. Knockdown of choline acetyl transferase (ChAT) or D2R in the vsNAc diminished the likelihood of intromission and ejaculation. Optogenetic manipulations reveal that DA signaling sustains male sexual behaviors by suppressing activities of D2R^vsNAc^ neurons. Moreover, ACh signaling promotes the initiation of mounting and intromission, but also induces the intromission-to-ejaculation transition by triggering a slowdown of DA rhythm. Therefore, dual ACh-DA dynamics harmonize in the vsNAc to drive sequential transitions of male mating behaviors.

## Introduction

In mammals, males engage in a stereotypical sequence of sexual behaviors when they encounter females in estrous. This sequence of male sexual behaviors consists of courtship (appetitive) behaviors, such as sniffing and chasing, followed by copulatory (consummatory) behaviors, such as mounting, intromission, and ejaculation in rodents (Everitt, 1990; Ågmo, 1997; Pfaus et al., 2001; Paredes and Ågmo, 2004; Hull and Dominguez, 2007; Balthazart and Ball, 2007). Appropriate expression of these sequential sexual behaviors is not only considered a measure of male attractivity, but also is important for the successful completion of sexual behaviors (Beach, 1976). It has been suggested that each sexual behavioral sequence is regulated by a distinctive neuronal mechanism. For example, the medial preoptic area (MPOA) has been implicated in copulatory behavior, while the nucleus accumbens (NAc) is thought to be involved in appetitive behavior (Balthazart and Ball, 2007; Everitt, 1990). However, recent studies have suggested that the MPOA controls mounting (Bayless et al., 2023; Karigo et al., 2021; Sano et al., 2013; Wei et al., 2018), and that the NAc may be involved in the facilitation of overall sexual behavior or the learning and reinforcement of male sexual behavior (Domínguez-Salazar et al., 2014; Pitchers et al., 2013; Beny-Shefer et al., 2017). Thus, sexual behavior is a complex sequence of actions involving multiple brain regions working in concert. However, the neural mechanisms that govern the sequential transitions of these sexual behaviors, specifically transitions from mounting to intromission to ejaculation, remain largely unclear.

Accumulating studies have suggested that DA signaling functions as a primary driver for sexual motivation and behaviors (Hull et al., 1992, 1993; Bitran and Hull, 1987; Sun et al., 2018, 2020; Dai et al., 2022; Fiorino and Phillips, 1999; Holstege et al., 2003). Moreover, administration of cholinergic agonists in the preoptic area (POA) or substantia nigra altered the number or frequency of intromissions preceding ejaculation in rodents, implicating a regulatory role of ACh signaling during male copulatory behaviors (Bitran and Hull, 1987; Hull et al., 1988; Winn, 1991). However, it remains unclear where and how these neurotransmitters interplay and regulate male sexual behaviors (Bitran and Hull, 1987; Sun et al., 2018, 2020; Dai et al., 2022; Qian et al., 2023; Hull et al., 2004; Hillegaart and Ahlenius, 1998). Early microdialysis studies had detected an overall increase in extracellular DA in the NAc during male sexual behaviors in rats (Damsma et al., 1992; Pfaus et al., 1990; Pleim et al., 1990). Recent fiber photometry studies recorded transient increases of DA levels during intromission and ejaculation in male mice using the high temporal resolution G protein-coupled receptor activation-based (GRAB)_DA_ sensor (Dai et al., 2022; Sun et al., 2018, 2020). Because previous studies have not suggested a critical involvement of the NAc in the regulation of sexual behavior (Coolen et al., 2004; Veening and Coolen, 2014; Hashikawa et al., 2016), the regulatory mechanism and functional significance of DA dynamics in the NAc during male sexual behaviors have not been carefully investigated.

In this study, we hypothesize that ACh signaling modulates DA dynamics in the NAc to regulate male sexual behaviors, based on recent reports that ACh signaling regulates DA release from the dopaminergic axon terminals in the striatum (Krok et al., 2023; Liu et al., 2022; Threlfell et al., 2012; Wang et al., 2014a, 2014b). By monitoring DA dynamics in various subregions of the NAc using the GRAB_DA2m_ sensor (Sun et al., 2018, 2020), we found that real-time DA dynamics in the ventral shell of the NAc (vsNAc) closely aligned with the sequential transitions of sexual behaviors in male mice. Consistent with our hypothesis, we observed a unique signature of dual ACh-DA rhythms in the vsNAc during intromission. Ex vivo brain slice imaging and computational modeling revealed that the dual ACh-DA rhythms were generated by reciprocal regulation between ACh and DA signaling via nAChR and D2R receptors, respectively. By optogenetic manipulations of ACh and DA release into the vsNAc, we demonstrated that dual ACh and DA dynamics in the vsNAc cooperatively drive the sequential transitions of male copulatory behaviors.

## Results

### DA dynamics in the vsNAc closely align with serial transitions of male sexual behaviors

After sniffing and chasing receptive female mice, male mice engage in repeated cycles of mounting and intromission, which culminate in ejaculation followed by a brief falling and immobile phase (Figure 1A). To study DA dynamics in the NAc during male sexual behaviors, we performed fiber photometry recording at the NAc of C57BL/6J male mice following injection with an adeno-associated virus (AAV) expressing GRAB_DA2m_ sensor in neurons under the human synapsin promotor (hSyn) (Figure 1B). Since the NAc is anatomically heterogeneous (Al-Hasani et al., 2015; Beier et al., 2015; Chen et al., 2021; Poulin et al., 2018), we examined DA dynamics in different subregions of the NAc by inserting optic fibers into the core (c), medial shell (ms), or ventral shell (vs) of the NAc, which was classified based on the expression pattern of substance P (Figures 1C and 1D) (Voorn et al., 1989). Distinct patterns of extracellular DA dynamics were detected in the cNAc, msNAc, and vsNAc during male sexual behaviors (Figures 1E–1J). Both the vsNAc and cNAc, but not msNAc, exhibited an increase in GRAB_DA2m_ signals that corresponded to all the sequential transitions of male sexual behaviors, from the initial encounter of female mice to the final intromission-to-ejaculation transition (Figures 1E–1G; Table S1). Among these, only the vsNAc exhibited a 1–2 Hz rhythmic fluctuation of GRAB_DA2m_ signals during intromission, which appeared to be concurrent with the intromission thrust rhythm (Figures 1H–1J; Table S2). These observations identify the vsNAc as a candidate brain region where DA dynamics may regulate the sequential transitions from intromission to ejaculation.

**Figure 1.**
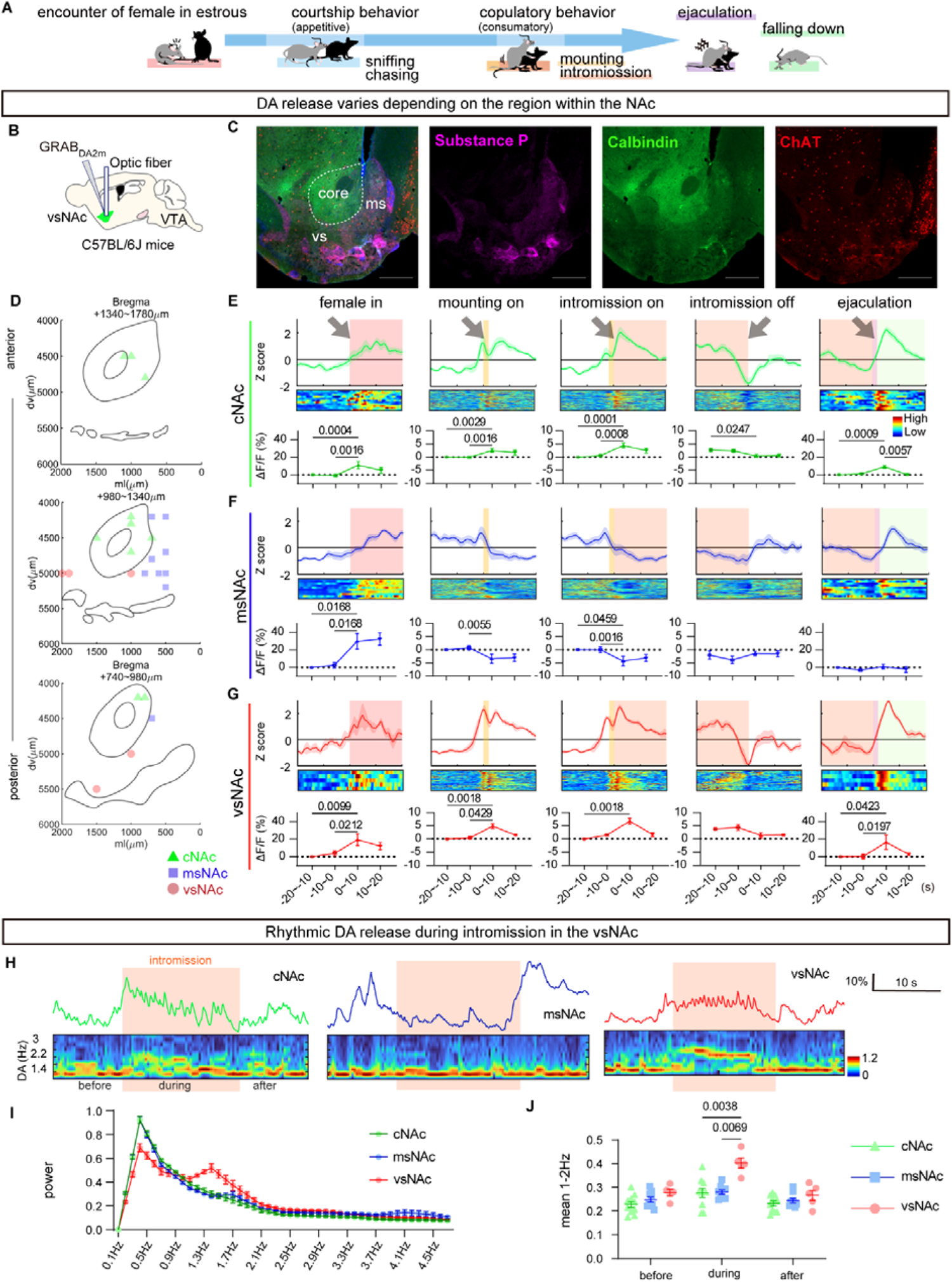
DA dynamics in the vsNAc closely align with serial transitions of male sexual behaviors. (A) Schematic of sexual behaviors in male mice. (B) Schematic of fiber photometry recording of GRAB_DA2m_ signals in the vsNAc of C57BL/6J mice. (C) Representative image showing immunostaining of ChAT, Calbindin, and ChAT expression in the NAc and surrounding regions (scale bar = 500 μm). (D) Schematics of recording sites of GRAB_DA2m_ imaging in various NAc subregions. (E–G) Top: Peri-event time plot (PETP) of z-scored ΔF/F of GRAB_DA2m_ fluorescence around each transition of male sexual behavior in the cNAc (E, *n* = 11 mice / group), msNAc (F, *n* = 10 mice / group), and vsNAc (G, *n* = 5 mice / group). Middle: Heatmap of PETP, with individual events/row; all data from males. Bottom: Mean ΔF/F of GRAB_DA2m_ of the three recording sites every 10 s in each behavioral event. See Table S1. (H) Representative traces of GRAB_DA2m_ dynamics during intromission and spectrum band calculated by FFT analysis in the cNAc (left), msNAc (center), and vsNAc (right). (I) Comparative power spectrum analysis of DA dynamics in the cNAc, msNAc, and vsNAc during intromission. (J) Mean power spectrum of 1–2 Hz frequency of DA dynamics before, during, and after intromission in the cNAc, msNAc, and vsNAc. See Table S2. Mean ± SEM. See also Figure S1.

### Local regulation of DA rhythm in the vsNAc during intromission

The ventral tegmental area (VTA) comprises a heterogeneous population of dopaminergic neurons (DA^VTA^ neurons) that, depending on their genetic and anatomical characteristics, send axonal projections to different target regions, including the NAc (Beier et al., 2015; Poulin et al., 2018). By two-color cholera toxin B subunit (CTB)-mediated retrograde labeling, we found that the vsNAc and msNAc received projections from the anterolateral (al) VTA and the posteromedial (pm) VTA, respectively (Figures S1A–S1C).

To compare the activities of the dopaminergic neurons in the alVTA and pmVTA during male sexual behaviors, we unilaterally injected AAV-hSyn-DIO-GCaMP6s and inserted optical fibers into the alVTA or pmVTA of *DAT-ires-Cre* (*DAT^Cre^*) mice, respectively (Figures S1D and S1E). Consistent with the different projection patterns, fiber photometry recording revealed that the alVTA, but not pmVTA, neurons exhibited GCaMP (Ca^2+^) dynamics in accordance with GRAB_DA2m_ dynamics in the vsNAc during male sexual behaviors (Figures S1F–S1H; Table S3). However, we did not observe rhythmic fluctuation of intracellular Ca^2+^ concentration in the alVTA or pmVTA during intromission (Figure S1F).

We also performed fiber photometry to monitor the neuronal activity in DA^VTA^ axons projecting to the vsNAc (DA^VTA→ vsNAc^) by injecting AAV-hSyn-DIO-GCaMP6s into the VTA and inserting optical fibers into the vsNAc of *DAT^Cre^* mice (Figure S1I). Unlike the neuronal activity in the somata of alVTA, the Ca^2+^ dynamics in DA^VTA→vsNAc^ axons were similar to GRAB_DA2m_ dynamics in the vsNAc, both in terms of high signal intensity and rhythmic fluctuations during intromission (Figures S1J and S1K; Table S4). By recording the 405 nm isosbestic GCaMP control signals, we confirmed that the rhythmic GCaMP signals in the vsNAc reflected real neuronal activity rather than motion artifacts during intromission (Figure S1K). These observations suggest that DA dynamics in the vsNAc are regulated locally within the vsNAc and, to some extent, independent of the somata localized in the VTA.

### Dual ACh-DA rhythms in the vsNAc during intromission

Previous studies have shown that ACh released from cholinergic interneurons promotes DA release from DA^VTA^ axons in the striatum (Krok et al., 2023; Liu et al., 2022; Threlfell et al., 2012; Wang et al., 2014a, 2014b). Using immunohistochemistry of choline acetyltransferase (ChAT) in *DAT^Cre^*mice, in which DA neurons were marked by GFP expression, we confirmed the existence of cholinergic interneurons (ChAT^vsNAc^) in proximity of the DA^VTA^ axons in the vsNAc (Figures S2A**-**S2C). To test our hypothesis that ACh signaling regulates DA dynamics in the vsNAc during male sexual behaviors, we unilaterally injected AAV-hSyn-DIO-GCaMP6s and inserted optic fibers into the vsNAc of *ChAT-ires-Cre* (*ChAT^Cre^*) male mice (Figures S2D–S2F; Table S5). Fiber photometry recording revealed that ChAT^vsNAc^ neurons exhibited rhythmic activity during intromission (Figure S2F). Next, we examined the ACh dynamics in the vsNAc by injecting AAV-hSyn-GRAB_ACh3.0_ into the vsNAc of C57BL/6J mice (Figures S2G–S2I; Table S5). Similar to the rhythmic fluctuation of GRAB_DA2m_ signals, we also observed the rhythmic fluctuation of GRAB_ACh3.0_ signals during intromission (Figure S2I). These results are consistent with the idea that the rhythmic activity of cholinergic interneurons may regulate the rhythmic release of ACh and possibly DA in the vsNAc during intromission.

Next, we performed dual fiber photometry imaging to simultaneously monitor ACh and DA dynamics after AAV-mediated expression of green-fluorescent GRAB_ACh3.0_ and red fluorescent GRAB_rDA2m_ in the vsNAc of C57BL/6J male mice (Figure 2A). Importantly, we observed a specific signature of dual ACh and DA rhythms during intromission (Figure 2B). Through the wavelet coherence analysis, we found that the coherence between ACh and DA rhythms in the 1–4 Hz band was significantly increased during intromission (Figures 2C, 2D, and S2J; Table S6 and S7). Both ACh and DA rhythms were characterized by a frequency of 1.5–2.2 Hz by as determined by fast Fourier transform analysis (Figure 2E). The ACh rhythm, but not DA rhythm, appeared just before the onset of intromission: the 1.5–2.2 Hz power of ACh rhythm was significantly higher than that of DA rhythm from 6-s before to 2-s after the intromission onset (Figure 2F; Table S6).

**Figure 2.**
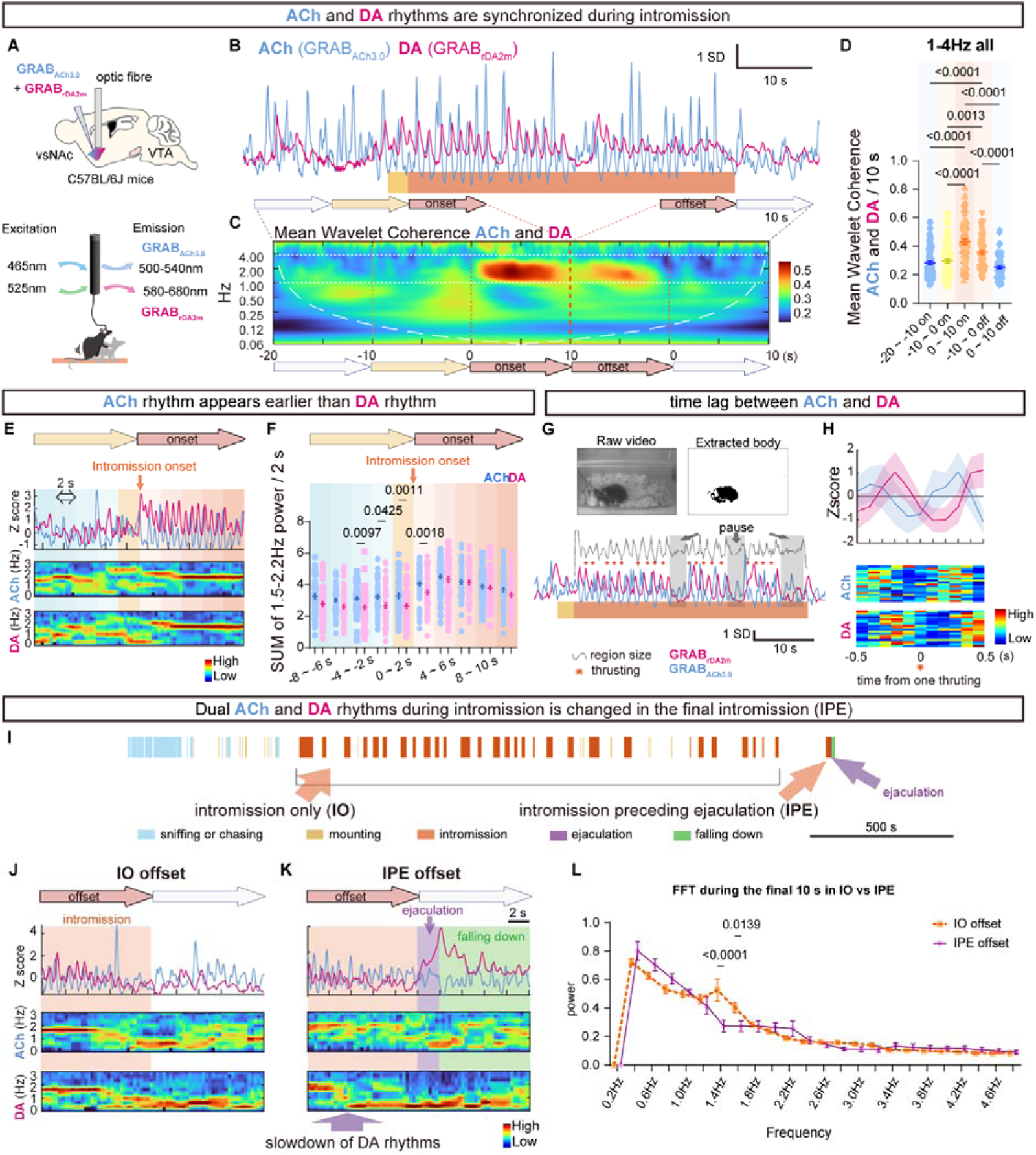
Dual ACh-DA rhythms are observed in the vsNAc during intromission. (A) Strategy for simultaneous fiber photometry recordings of ACh and DA dynamics in the vsNAc. (B) Representative traces of GRAB_ACh3.0_ and GRAB_rDA2m_ before, during, and after intromission. (C) Heatmap plot of mean wavelet coherence between GRAB_ACh3.0_ and GRAB_rDA2m_ dynamics around the extracted 50 s intromission. (D) Comparison of 1–4 Hz wavelet coherence between GRAB_Ach3.0_ and GRAB_rDA2m_ among every 10 s around intromission. (E) Representative Z-scored traces of GRAB_ACh3.0_ and GRAB_rDA2m_ in the vsNAc (top); heatmap showing the power spectrum processed by the FFT analysis of GRAB_ACh3.0_ (center) and GRAB_rDA2m_ (bottom) around intromission onset. (F) Comparison of the total amounts of 1.5–2.2 Hz power collected every 2 s around intromission onset. (G) Thrusting was determined by the region size change of the extracted male and female mice. (H) GRAB_ACh3.0_ and GRAB_rDA2m_ around one thrusting during one intromission. (Top) The average traces (bottom) heatmap plot. (I) Representative laser illustration of sexual behavior. (J and K) Same as (E), FFT analysis around IO offset (J), and IPE offset (K). (L) Power spectrum of GRAB_DA2m_ in the vsNAc during 10 s offset of IO and IPE (*n* = 5 mice / group). Two-way ANOVA Šídák’s multiple comparisons test over IO offset and IPE offset every 0.2 Hz. Mean ± SEM. See Table S6 (D, F, and L).

To assess the time-lag between ACh and DA rhythms during intromission, we compared the ACh/DA rhythm with the pelvic thrust rhythm by analyzing the dynamic changes in the body size of male and female mice in the video (Figure 2G). By plotting the ACh and DA rhythms around each thrusting event, we found the trough of the ACh rhythm was approximately 0.1 s before, whereas that of DA rhythm was ∼0.1–0.2 s after the thrusting (Figure 2H). Moreover, the ACh rhythm, but not the DA rhythm, persisted during the short pauses of pelvic thrusting during long intromissions (Figure 2G). Taken together, these observations suggest that the ACh rhythm may function as the primary driver in generating the dual ACh-DA rhythm during intromission (Figures 2E–2H).

### Signatures of the intromission preceding ejaculation (IPE)

While the majority of intromissions do not lead to ejaculation (intromission only (IO)), the last intromission typically transitions into ejaculation (intromission preceding ejaculation (IPE)) during a successful cycle of sexual behaviors (Figure 2I). Thus, a comparison of dual ACh-DA dynamics between IO and IPE may provide insights into the neural basis for controlling the intromission-to-ejaculation transition. Interestingly, the coherence of the dual ACh-DA rhythms was significantly enhanced at the intromission onset of IPE relative to IO (Figures S2K and S2L; Table S7). While the dual 1.5–2.2 Hz ACh-DA rhythms continued until the termination of IO (Figure 2J), there was a specific slowdown of the 1.5–2.2 Hz DA rhythm just before ejaculation that coincided with the big surge of DA release in the IPE (Figure 2K). Importantly, this brief suppression of DA rhythm was only observed in the vsNAc (Figure 2L; Table S6), not in the msNAc or cNAc, during the offset of IPE (Figure S2M; Table S8), suggesting that it may be a prerequisite for the intromission-to-ejaculation transition.

### ACh signaling regulates DA release in the vsNAc

By AAV-mediated Cre-dependent GFP expression in the vsNAc of *ChAT^Cre^*mice, we confirmed that cholinergic neurons are locally distributed in the vsNAc (Figures S3A and S3B). To examine whether ACh signaling could regulate DA release by acting on DA^VTA^ axons in the vsNAc, we injected AAV-hSyn-FLEX-ChrimsonR and AAV-hSyn-GRAB_DA2m_ into the vsNAc of *ChAT^Cre^* mice, followed by ex vivo NAc brain slice imaging coupled with optogenetics and pharmacological treatments (Figures 3A–3C). Optogenetic activation of ChAT^vsNAc^ neurons evoked an immediate and robust increase in GRAB_DA2m_ signals in the NAc slices (Figures 3B and 3C). This cholinergic stimulation of DA release was suppressed by a nAChR antagonist DhβE at all frequencies (Figures 3B and 3C). Similarly, the cholinergic activation of DA release was suppressed by a mAChR antagonist atropine during 20 or 40 Hz stimulation of ChAT^vsNAc^ neurons (Figures 3B and 3C). However, in response to 1 or 10 Hz stimulation of ChAT^vsNAc^ neurons, the blockade of mAChR induced a sharp increase of GRAB_DA2m_ signals (Figures 3B and 3C), possibly due to the disinhibition of mAChR-mediated suppression of ACh release (Cachope et al., 2012). Accordingly, in situ hybridization detected the expression of all five mAChR1–5 (*Chrm1*–*5*) in the majority of ChAT^vsNAc^ neurons (Figures S3C and S3E; Table S9). In addition, DA^VTA^ neurons expressed both nAChR (Jones et al., 2001) and mAChR1-5 (Figures S3D and S3F; Table S9). These results suggest that ACh promotes DA release via nAChR signaling but also inhibits DA release in some contexts via mAChR signaling in the vsNAc.

**Figure 3.**
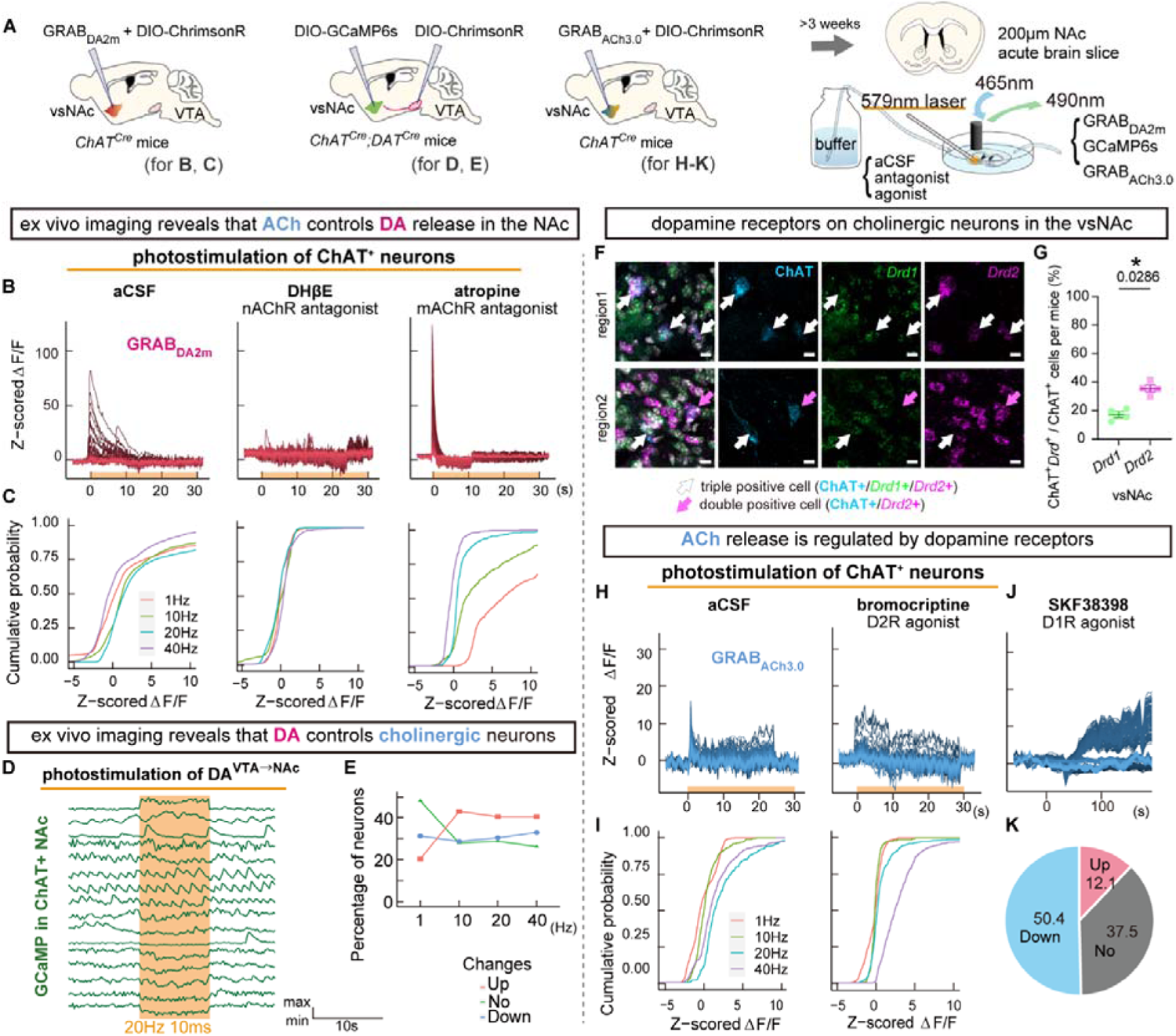
Ex vivo NAc brain slice imaging reveals mutual regulation of ACh and DA signaling. (A) Diagram of ex vivo experiments. Letters in parentheses indicate the corresponding figures. (B and C) Optogenetic stimulation of ChAT^NAc^ neurons and imaging of GRAB_DA2m_. Representative traces of z-scored ΔF/F of GRAB_DA2m_ signals in response to 10 ms-photostimulation at 20 Hz for 30 s with neither DHβE or atropine (left), with only DHβE (center), and with only atropine (right). Bottom orange bars indicate the duration of photostimulation (B). Cumulative probability plots of mean z-scored ΔF/F of GRAB_DA2m_ signals for 30 s after the onset of photostimulation. Plot colors indicate frequencies of the photostimulation. The experimental conditions are the same as those of the figures directly above. *n* = 2332 (left), 776 (center), and 570 (right) ROIs at any photostimulation frequency (C). (D and E) Optogenetic stimulation of DA^VTA→NAc^ axon terminals and Ca^2+^ imaging with GCaMP. Representative traces of GCaMP signals in response to 10 ms-photostimulation at 20 Hz for 10 s 20 Hz (D). Classification of ChAT^NAc^ neurons by the response to photostimulation. *n* = 181 cells (E). (F and G) ChAT^+^ neurons in the vsNAc are colocalized with *Drd1* and *Drd2* (scale bars = 10μm) (F). The ratio of ChAT^+^*Drd1*^+^ neurons / ChAT^+^ neurons and that of ChAT^+^*Drd2*^+^ neurons / ChAT^+^ neurons (G). See Table S10. (H and I) Optogenetic stimulation of ChAT^NAc^ neurons and imaging of GRAB_ACh3.0_. Same as (B), but for GRAB_ACh3.0_ signals in aCSF with (left) or without bromocriptine (right) (H). Same as (C), but for GRAB_ACh3.0_ signals. *n* = 657 (1 Hz), 490 (10 Hz), 325 (20 Hz), and 654 (40 Hz) ROIs (left). *n* = 657 (1 Hz), 165 (10 Hz), 654 (20 Hz), and 489 (40 Hz) ROIs (right) (I). (J and K) Bath application of a D1R agonist and imaging of GRAB_ACh3.0_. All traces of z-scored ΔF/F of GRAB_ACh3.0_ signals in response to bath application of SKF38398 (J). Pie chart of the response pattern of the ROI. *n* = 685 ROIs (K).

### DA signaling reciprocally regulates ACh release in the vsNAc

To examine whether DA signaling reciprocally regulated the activity of ChAT^vsNAc^ neurons, we performed ex vivo Ca^2+^ imaging with NAc slices following injection of AAV-Syn-FLEX-ChrimsonR into the VTA and AAV-hSyn-DIO-GCaMP6s into the vsNAc of *ChAT^Cre^*;*DAT^Cre^* mice (Figures 3A, 3D, and 3E). Optogenetic (1 Hz) stimulation of DA^VTA➔NAc^ axons evoked different responses in the ChAT^NAc^ neurons: ∼20% increased, ∼30% decreased, and ∼50% exhibited no change in intracellular Ca^2+^ levels (Figure 3E). By contrast, higher frequency (≥10 Hz) stimulation of DA^VTA➔NAc^ axons increased the activity of ∼40% ChAT^NAc^ neurons and decreased the activity of ∼30% ChAT^NAc^ neurons (Figure 3E). Notably, a small subset of ChAT^NAc^ neurons displayed rhythmic fluctuation of intracellular Ca^2+^ levels in the NAc slices (Figure 3D), which was consistent with the rhythmic fluctuation of Ca^2+^ levels in ChAT^vsNAc^ neurons and rhythmic release of ACh in the vsNAc observed by in vivo fiber photometry during intromission (Figures S2D–S2I).

The existence of both *Drd1*^+^ChAT^+^ and *Drd2*^+^ChAT^+^ neurons in the vsNAc (Figures 3F and 3G; Table S10) raised two potential mechanisms for the regulation of ChAT^vsNAc^ neurons by DA signaling: 1) inhibition of ChAT^vsNAc^ neurons via D2R-mediated Gi/o signaling; 2) activation of ChAT^vsNAc^ neurons via D1R-mediated Gs signaling (Neve et al., 2004). To probe these possibilities, we injected AAV-Syn-FLEX-ChrimsonR and AAV-hSyn-GRAB_ACh3.0_ into the vsNAc of *ChAT^Cre^* mice (Figures 3A and 3H–3K). Optogenetic activation of ChAT^vsNAc^ neurons increased GRAB_ACh3.0_ signals in the NAc slices, which was suppressed by D2R agonist bromocriptine under all conditions except for 40 Hz stimulation (Figures 3H and 3I). Thus, the D2R-mediated inhibition of ChAT^NAc^ neurons could suppress the release of ACh from ChAT^NAc^ neurons following ≤20 Hz optogenetic stimulation (Figure 3I). Next, we investigated whether ACh release was also modulated by D1R signaling in the NAc slices by application of D1R agonist SKF38398. Activation of D1R signaling caused distinct changes in ACh release in the NAc slices, with 12.1% (up), 50.4% (down), and 37.5% (no change) in GRAB_Ach3.0_ signals (Figures 3J and 3K). Taken together, these results suggest the reciprocal regulation of ACh release by DA signaling in the vsNAc, which is potentially mediated by both D1R and D2R expressed on ChAT^vsNAc^ neurons.

### Modeling of dual ACh-DA rhythms by reciprocal regulation between ACh and DA signaling

To further study how the dual ACh-DA rhythms were generated by reciprocal regulation between ACh and DA in the vsNAc during intromission, we performed computational modeling using a simplified neuronal assemblage consisting of the dopaminergic 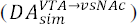 axon terminals, cholinergic neurons 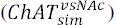, and the extracellular environment (Figures 4A and S4A–S4D). In this modeling, we defined the release probability measurement (*RPM*) of the neuronal assemblage as a measure of the intensity of neuronal activity (Figure 4B). We set *Activation* as a constant value to mimic the external stimuli, such as the spontaneous firing of each neuronal assemblage or stimulation by other neurons (Figure S4A). Additionally, the *Receptor* function represents the effect of each receptor upon binding of ACh or DA (Figure 4B). Both *Activation* and *Receptor* constantly change the *RPM* that determines the volume of neurotransmitter (NT) released to the extracellular environment (Figures 4B and S4A).

**Figure 4.**
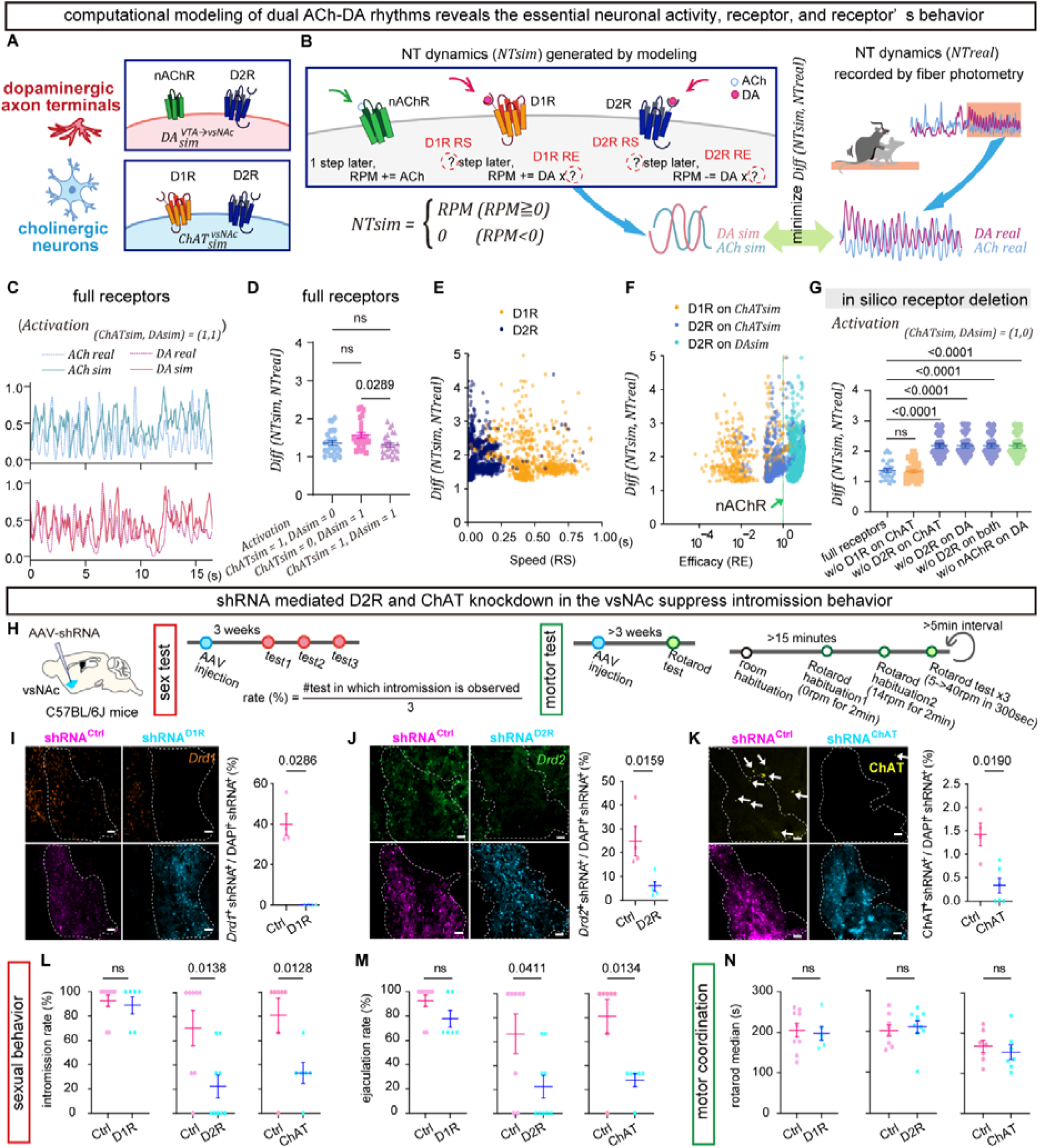
Computational modeling reveals that nAChR and D2R signaling are necessary for the generation of dual ACh-DA rhythms. (A–G) Computational modeling revealed the essential receptor and behavior of each candidate receptor during dual ACh-DA rhythms. (A) Setting of receptors distribution in the model. (B) Schematic representation of the model. (C) Representative traces of *NT_sim_* and *NT_real_* of ACh and DA under the full receptor distribution. (D) Comparison of *Diff(NT_sim_, NT_real_)* with four full receptors under three activation conditions; only 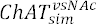, only 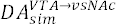 vs. both 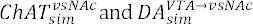. See Table S11. (E) Comparison of *Diff(NT_sim_, NT_real_)* with four full receptors vs. *RS* (Receptor speed) of D1R and D2R during generation of dual ACh-DA rhythms. (F) Comparison of *Diff(NT_sim_, NT_real_)* with four full receptors vs. *RE* (Receptor efficacy) of D1R and D2R in 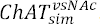 and D2R in 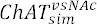 during generation of dual ACh-DA rhythms. The *RE* of nAChR is shown as a green dashed line. (G) Comparison of *Diff(NT_sim_, NT_real_*) under activation only with 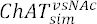 with each receptor deleted condition. See Table S12. (D–G) *n* = 25 *NT_real_* signals from 3 mice. (H–N) Knockdown of D2R or ChAT, but not D1R, expression in the vsNAc diminished male sexual behavior. (I) Schematic for AAV-shRNA mediated gene knockdown (right) and experimental schedule of sexual behavior tests (center) and rotarod tests following AAV-shRNA injection (left). (I–K) Representative images showing AAV-shRNA^D1R^ (I), AAV-shRNA^D2R^ (J) or AAV-shRNA^ChAT^ (K) injected vsNAc region (left). Scale bars = 50μm. The effect of shRNA-mediated gene knockdown of D1R (I, *n* = 4 mice / group), D2R (J, *n* = 4 mice / group), and ChAT (K, *n* = 4 mice / group) on target gene expression (right). Knockdown score = (# D1R^+^ (I), D2R^+^ (J), ChAT^+^ (K) ) cells / (# DAPI). See Table S13. (L–N) The effect of shRNA-mediated gene knockdown of D1R (*n* = 8 shRNA^Ctrl^ vs. 6 shRNA^D1R^ mice), and D2R (*n* = 7 shRNA^Ctrl^ vs. 9 shRNA^D2R^ mice), and ChAT (*n* = 7 shRNA^Ctrl^ vs. 6 or 7 shRNA^ChAT^ mice) on the rate of intromission (L) and ejaculation (M) as well as the motor coordination scored by the median duration of rotarod test (N). See Table S14. Intromission rate (%) = (# test in which the intromission is observed) / 3; ejaculation rate (%) = (# test in which the ejaculation is observed) / 3. Mean ± SEM. See also Figure S5.

The binding of DA to different dopamine receptors activates distinct intracellular G protein signaling to regulate neuronal activity: D1R triggers Gs signaling to activate neuronal activity, whereas D2R activates G protein-gates inwardly rectifying potassium (GIRK) channels-mediated signaling to inhibit neuronal activity (Cruz et al., 2004; Greif et al., 1995; Neve et al., 2004). On the other hand, the binding of ACh to nAChR causes a rapid influx of cations, such as Na^+^ and Ca^2+^, to activate neuronal activity (Galzi et al., 1992; Fucile, 2004). Herein, we focused on nAChR rather than mAChR1-5 to simplify our modeling to explore the characteristics of D1R and D2R relative to nAChR. We hypothesized that the characteristics of each receptor consist of two axes: the effect intensity (effect: *RE*) with which the receptor changes neuronal activity; the processing speed (speed: *RS*) of the receptor in affecting the neuronal activity upon binding of ACh or DA (Figure 4B).

### Cholinergic activation is required for modeling dual ACh-DA rhythms

The goal of the modeling is to estimate the *RE* and *RS* values of each receptor to minimize the *Diff*(*NTsim*, *NT_real_*) , i.e., the difference between simulated signals (*NT_sim_*) and experimentally recorded in vivo GRAB signals (*NT_real_*). To simplify the model, both the *RS* and *RE* values of the nAChR were defined as 1, and the *RS* and *RE* values of D1R and D2R relative to the nAChR were explored (Figure 4B). Based on the ex vivo imaging (Figure 3) and in situ hybridization results (Figure S3) and previous report of D2R expression in dopaminergic neurons (De Mei et al., 2009), we placed D1R and D2R on 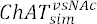 neurons and nAChR and D2R on 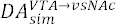 axon terminals in this model (Figure 4A). Importantly, we found that the simulated dual rhythms of ACh and DA closely resembled the experimentally recorded ACh and DA rhythms (Figure 4C).

To investigate the relative contribution of 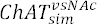 neurons and 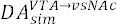 axon terminals, we compared the similarity between *NT_real_* and *NT_sim_ (Diff(NT_sim_ , *NT_real_*))* in three activation patterns: 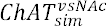 and 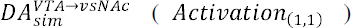; only 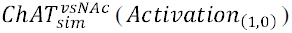; only 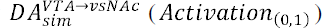 (Figure S4A). The similarity between *NT_real_* and *NT_sim_* rhythms suggested that the *Activation*_(1,1)_ and *Activation*_(1,0)_ conditions were much better than the *Activation*_(0,1)_ condition to simulate dual ACh-DA rhythms (Figure 4D; Table S11). These observations suggest that the activation of cholinergic neurons is necessary to generate dual ACh-DA rhythms.

### D2R, but not D1R, is required for modeling dual ACh-DA rhythms

We visualized the explored values of the *RS* and *RE* for each assumed receptor during the simulation. Notably, the *RS* of D2R was in the range of 5–50 ms, whereas the *RS* of D1R was approximately 350–400 ms (Figure 4E), indicating that the processing speed of D2R is approximately 7–80 fold faster than that of D1R in our model. This finding is consistent with the fact that the GIRK-mediated D2R signaling is much faster than Gs-mediated D1R signaling. On the other hand, the *RE* of D1R or D2R on 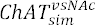 neurons are approximately 1/100 or 1/50 of the *RE* of nAChR on 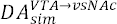 axon terminals, respectively. By contrast, the *RE* of D2R on 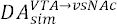 axon terminals is about 2 to 4 times of the *RE* of nAChR (Figure 4F), such that the effect intensity of D2R on 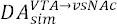 axon terminals is hundreds fold larger than that of D2R on 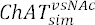 neurons in our model.

To determine the receptors essential for generating the dual ACh-DA rhythms, we performed simulations to evaluate the similarity between *NT_sim_* and *NT_real_ following* ablation of each *receptor* (Figures 4G, S4E, and S4F; Table S12). Consistent with estimation that D2R has larger *RE* than D1R, the ablation of D2R, but not D1R, on 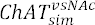 neurons worsens the similarity between *NT_sim_* and *NT_real_* as compared with the full receptor condition (Figure 4G). Similarly, the ablation of D2R or nAChR on 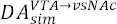 axon terminals significantly worsened the similarity between *NT_sim_* and *NT_real_* . These observations underscore the importance of D2R-mediated DA signaling and nAChR-mediated ACh signaling in the generation of dual ACh-DA rhythms.

### Knockdown of D2R and ChAT in the vsNAc diminishes intromission and ejaculation

To validate our simulation results in vivo, we performed RNAi-mediated knockdown of D1R, D2R, or ChAT expression by injecting AAV expressing short hairpin (sh)RNA targeting *Drd1*, *Drd2,* or *Chat* mRNA into the vsNAc of male mice. A (Figures 4H–4N, and S4G–S4N). In situ hybridization verified efficacy of knockdown of target gene expressions in the vsNAc (Figures 4I–4K, and S4N; Table S13). Three weeks post-AAV-shRNA injection, sexual behavior tests were conducted over a period of 3 weeks (Figures 4H, 4L, 4M, and S4G–S4M). Consistent with the simulation results of in silico receptor ablation (Figure 4G), knockdown of D2R or ChAT, but not D1R, expression in the vsNAc diminished the frequency of sexual behaviors, such as intromission and ejaculation, relative to the scrambled shRNA-expressing mice (Figures 4L and 4M; Table S14). Moreover, knockdown of D2R or ChAT prolonged the average latency of mounting, intromission, and ejaculation (Figures S4G, S4J, and S4M; Table S15), while knockdown of D2R reduced the average number of bouts and total duration of mounting and intromission (Figures S4H, S4I, S4K, and S4L; Table S15). Importantly, neither D2R nor ChAT knockdown affected the motor coordination of the mice as measured by the rotarod test (Figure 4N; Table S14), excluding the possibility that male sexual behaviors were indirectly affected by impairing motor coordination. These results indicate that both ACh and DA signaling play important roles in male copulatory behaviors. Based on the results of ex vivo brain slice imaging, computational modeling, and RNAi knockdown experiments, we concluded that the dual ACh-DA rhythms are likely generated in the vsNAc by reciprocal ACh-DA regulations mediated by nAChR and D2R signaling in DA^VTA→vsNAc^ axon terminals and D2R signaling in ChAT^vsNAc^ neurons, respectively.

### High level of DA signaling in the vsNAc is important for the maintenance of intromission

A high level of extracellular DA was observed in the vsNAc, and only the vsNAc exhibited the 1.5–2.2 Hz DA rhythm during intromission (Figure 1). To examine whether DA signaling plays a crucial role during intromission, we optogenetically manipulated the DA^VTA➔vsNAc^ axon terminals to investigate whether changing DA dynamics in the vsNAc could affect male copulatory behaviors (Figures 5 and S5). We bilaterally injected AAV-Ef1a-DIO-ChR2 or AAV-Ef1a-DIO-eNpHR3.0 into the VTA and inserted optic fibers into the vsNAc of *DAT^Cre^* mice (Figure 5D). We performed optogenetic stimulation of DA^VTA➔vsNAc^ axons at 20 Hz (10 ms width) because optogenetic stimulation at 1 or 10 Hz did not increase DA release (Figures 5A–5C, S5A, and S5B; Table S16). Notably, optogenetic activation of the DA^VTA➔vsNAc^ axons either before intromission (during sniffing, chasing, or mounting) or during intromission (after two pelvic thrust movements) increased the total duration of intromission (Figures 5G and 5J; Table S17). Additionally, optogenetic activation before intromission also increased the number of intromission bouts, although it did not immediately trigger intromission (Figures 5H and 5I; Table S17). Conversely, optogenetic suppression of the DA^VTA→vsNAc^ axons before or during intromission did not affect male sexual behaviors (Figures 5G–5K, and S5C–S5G; Table S17 and S18), possibly owing to incomplete suppression of DA signaling (see discussion). These results suggest that a high level of DA signaling in the vsNAc promotes the initiation and maintenance of intromission (Figure 5L).

**Figure 5.**
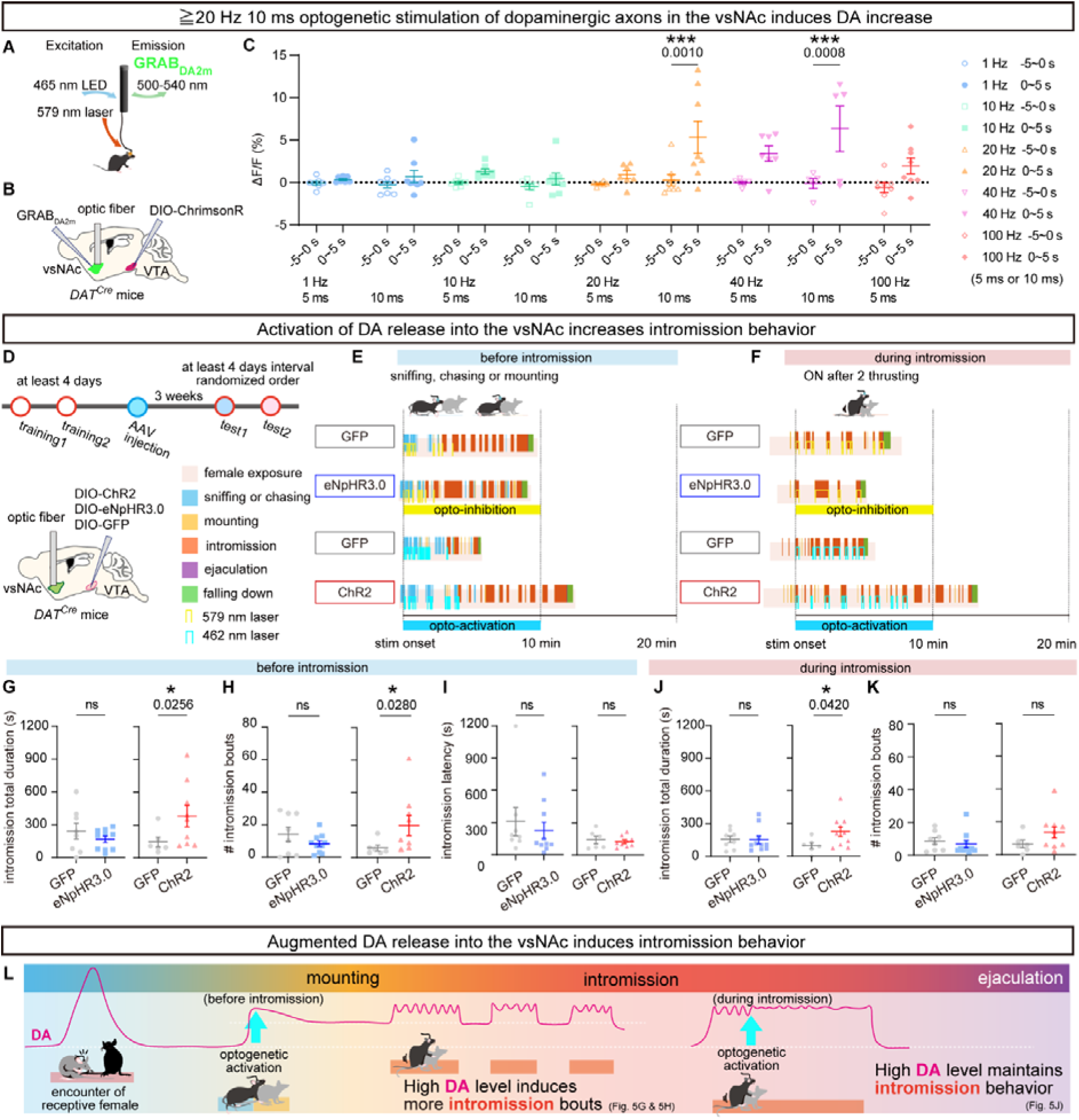
Optogenetic activation of DA release into the vsNAc increases intromission total duration. (A–C) The effects of optogenetic activation of DA^VTA→vsNAc^ axons on DA release level in the vsNAc. (A) Strategy for simultaneous optogenetic manipulation of DA^VTA→vsNAc^ axons and recording of GRAB_DA2m_ in the vsNAc. (B) Surgery design. (C) Quantitation of DA release level following optogenetic stimulation of dopaminergic axons in the vsNAc with various frequencies (*n* = 7 [1 Hz 5 ms], 8 [1 Hz 10 ms], 7 [10 Hz 5 ms], 8 [10 Hz 10 ms], 7 [20 Hz 5 ms], 8 [20 Hz 10 ms], 7 [40 Hz 5 ms], 5 [40 Hz 10 ms], and *n* = 8 [100 Hz 5 ms] signals. Average of ΔF/F of GRAB_DA2m_ fluorescence during -5 ∼ 0 s vs. 0 ∼ 5 s around the stimulation onset). See Table S16. (D–K) The effects of optogenetic activation of DA^VTA→^ ^vsNAc^ axons on male sexual behavior. (D) Strategy for optogenetic manipulation. Top: experimental schedule. Bottom: surgery design. (E and F) Laser illustration of sexual behavior and optogenetic manipulation before (E) and during intromission (F). (G–I) The effects of optogenetic manipulation of DA axons in the vsNAc before intromission (*n* = 8 GFP^DA^ vs. 10 eNpHR3.0^DA^ mice; *n* =6 GFP^DA^ vs. 9 ChR2^DA^ mice) on the intromission total duration (G), the number of intromission bouts (H), and intromission latency (I). (I) Intromission latency (s) = the 1^st^ intromission onset – the 1^st^ optogenetic stimulation onset. (J and K) The effects of optogenetic manipulation of DA axons in the vsNAc during intromission (*n* = 8 GFP^DA^ vs. 10 eNpHR3.0^DA^ mice; *n* = 6 GFP^DA^ vs. 10 ChR2^DA^ mice) on the intromission total duration (J) and the number of intromission bouts (K). (L) Enhancing DA release level in the vsNAc before intromission increases intromission bouts, while those during intromission prolongs intromission duration. Mean ± SEM. See also Figure S6 and Table S17 (G–K). \**p* < 0.05, \*\*\**p* < 0.001.

### DA-mediated inhibition of D2R^vsNAc^ neurons sustains male sexual behaviors, whereas suppression of D1R^vsNAc^ neurons ensures forward progression of sexual behaviors

To elucidate how DA dynamics regulate activity of D1R- and D2R-expressing neurons in the vsNAc during male sexual behaviors, we first confirmed the distribution of D1R and D2R in the NAc (Figures 6A–6C). The number of D1R^+^ neurons was twice that of D2R^+^ neurons in all subregions of the NAc (Figure 6B): 67.13% of dopamine receptor neurons in the vsNAc express *Drd1*, 30.26% express *Drd2*, and 2.60% express both (Figure 6C). Next, we unilaterally injected AAV-hSyn-DIO-GCaMP6s and inserted optic fibers in the vsNAc of *Drd1-ires-Cre* (*Drd1^Cre^*) and *Drd2-ire-Cre* (*Drd2^Cre^*) mice, respectively (Figures 6D–6F, S6A, and S6B; Table S19 and S20). Fiber photometry recording revealed that the activity of both D1R^vsNAc^ and D2R^vsNAc^ neurons were suppressed during intromission (Figures 6E, 6F, S6A, and S6B), despite the opposite effects of D1R (excitatory) and D2R (inhibitory) signaling on neuronal activity (Cruz et al., 2004; Greif et al., 1995; Neve et al., 2004). The simplest explanation for this paradoxical observation is that DA signaling preferentially activates the D2R-mediated inhibition of neuronal activities during intromission owing to the different thresholds for D1R and D2R activation (Richfield et al., 1989; Bazzett et al., 1991; Hull et al., 1992) (see discussion).

**Figure 6.**
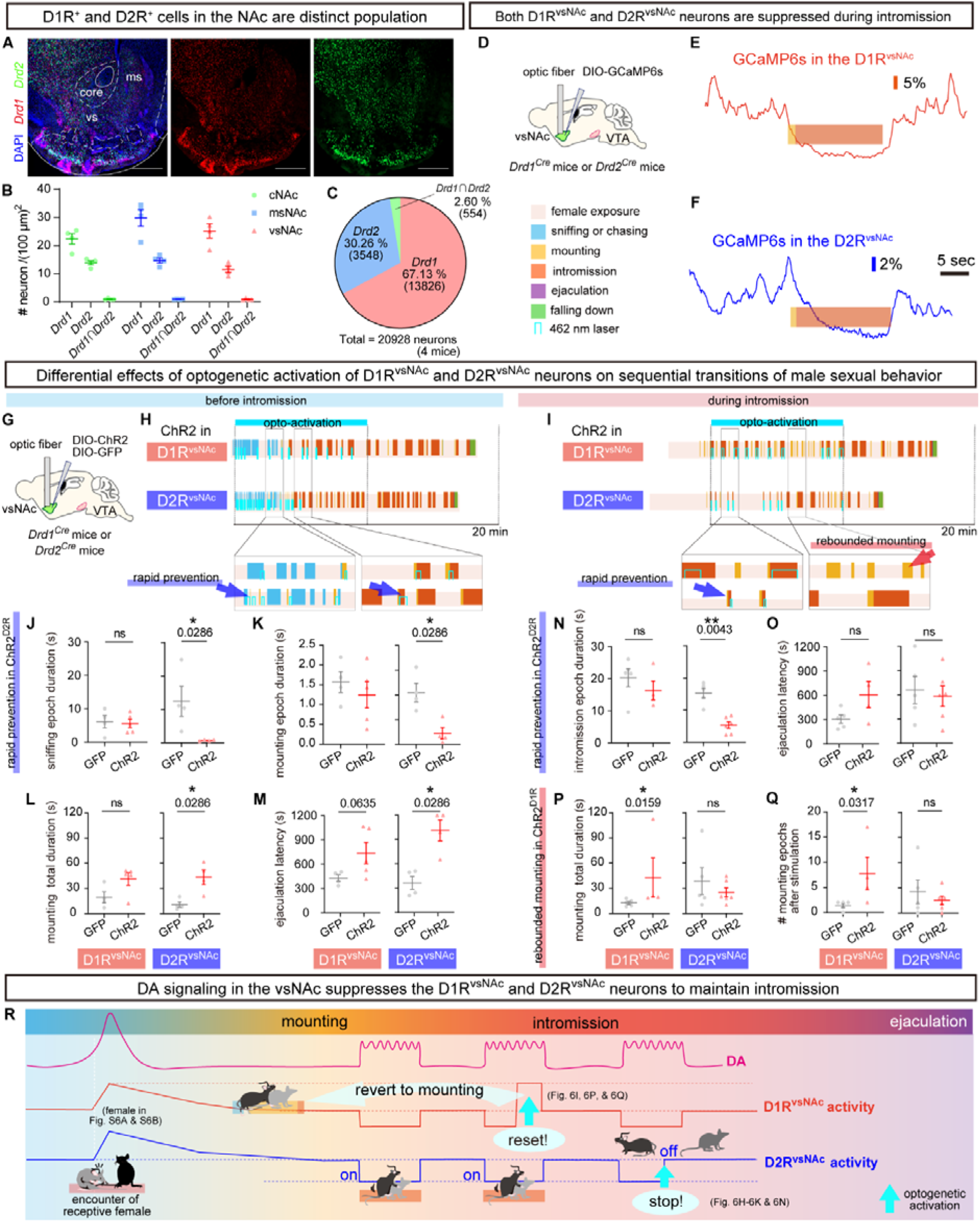
Suppression of D1R^vsNAc^ and D2R^vsNAc^ neurons during intromission are both required to maintain intromission. (A) Representative image showing in situ staining of *Drd1* and *Drd2* in the NAc (scale bar = 500 μm). (B) # Neurons expressing *Drd1*, *Drd2*, or both *Drd1* and *Drd2* in each subregion of the NAc /(100 μm)^2^ (*n* = 4 mice / group). (C) Percentage of neurons expressing only *Drd1*, only *Drd2*, or both *Drd1* and *Drd2* in the vsNAc. (D) Strategy for imaging of GCaMP6s expressed in the D1R or D2R in the vsNAc of *Drd1^Cre^* or *Drd2^Cre^* mice. (E and F) The neuronal activity of D1R^vsNAc^ (E) and D2R^vsNAc^ (F) are suppressed during intromission. (G) Strategy for optogenetic manipulation of D1R^vsNAc^ and D2R^vsNAc^. (H and I) Laser illustration of sexual behavior and optogenetic manipulation of D1R^vsNAc^ and D2R^vsNAc^ before (H) and during intromission (I). Blue arrows indicate the representative ethograms suggesting the rapid prevention effect in ChR2^D2R^ and a red arrow indicates the ethogram suggesting the effect of rebounded mounting in ChR2^D1R^. (J–M) The effect of the optogenetic activation of D1R^vsNAc^ (left) and D2R^vsNAc^ neurons (right) before intromission (*n* = 5 GFP^D1R^ vs. 4 ChR2^D1R^ mice; *n* = 4 GFP^D2R^ vs. 4 ChR2^D2R^ mice) on the epoch duration of sniffing (J), epoch duration of mounting (K), mounting total duration (L), and ejaculation latency (M). (N–Q) The effect of the optogenetic activation of D1R^vsNAc^ (left) and D2R^vsNAc^ neurons (right) during intromission (*n* = 4 GFP^D1R^ vs. 5 ChR2^D1R^ mice; *n* = 6 GFP^D2R^ vs. 5 ChR2^D2R^ mice) on the epoch duration of intromission (N), ejaculation latency (O), mounting total duration (P), mounting bouts (Q). (J, K, and N) Epoch duration (s) = the target behavior offset – the optogenetic stimulation onset. (M and O) Ejaculation latency (s) = the ejaculation onset – the 1^st^ optogenetic stimulation onset. (Q) # mounting is the number of mounting behaviors observed during 10 min after the onset of the optogenetic stimulation, excluding the mounting preceding intromission. (R) DA release level around female encounter activates D1R^vsNAc^ and D2R^vsNAc^ neurons, while those during intromission suppresses D1R^vsNAc^ and D2R^vsNAc^ neurons. Activation of D1R^vsNAc^ neurons during intromission induces mounting, previous behavior, by mimicking the neuronal activity around female encounter. Activation of D2R^vsNAc^ neurons rapidly stops any sexual behavior. Mean ± SEM. See also Figure S7; Table S21 (J–Q) and S22 (J–Q). \**p* < 0.05, \*\**p* < 0.01.

To study the functional importance of suppression of D1R^vsNAc^ and D2R^vsNAc^ neurons during intromission, we specifically expressed ChR2 in the D1R^vsNAc^ or D2R^vsNAc^ neurons by injecting AAV-Ef1a-DIO-ChR2 into the vsNAc of *Drd1^Cre^* or *Drd2^Cre^* mice, respectively (Figures 6G–6Q, S6C–S6F; Table S21 and S22). While optogenetic activation of D1R^vsNAc^ neurons before intromission did not affect male sexual behaviors, optogenetic stimulation of these neurons during intromission increased the total mounting duration by inducing repetitive mounting behaviors (Figures 6P and 6Q). Thus, the suppression of D1R^vsNAc^ neurons during intromission is critical for ensuring the forward progression of male sexual behaviors.

On the other hand, optogenetic activation of D2R^vsNAc^ neurons during sniffing, chasing, mounting, and intromission immediately stopped any of these sexual behaviors (Figures 6J, 6K, and 6N). Consequently, optogenetic stimulation of D2R^vsNAc^ neurons before intromission increased total mounting duration and prolonged ejaculation latency (Figures 6L and 6M), whereas optogenetic stimulation of D2R^vsNAc^ neurons during intromission decreased total intromission duration by shortening the epoch duration of intromission (Figure 6N and S6F). These observations indicate that DA-mediated suppression of D2R^vsNAc^ neurons is necessary for sustaining male sexual behaviors including intromission (Figure 6R).

### ACh signaling in the vsNAc promotes sequential transitions of intromission and ejaculation

To study how ACh signaling in the vsNAc regulates male sexual behaviors, we bilaterally injected AAV-Ef1a-DIO-ChR2 or AAV-Ef1a-DIO-eNpHR3.0 and inserted optic fibers into the vsNAc of *ChAT^Cre^* male mice (Figure 7A). Next, we optogenetically activated or inhibited the ChR2-or eNpHR3.0-expressing ChAT^vsNAc^ neurons, respectively, before or during intromission and examined the effects on male copulatory behaviors (Figures 7B–7G, and S7). While optogenetic activation of ChAT^vsNAc^ neurons before intromission did not affect mating behaviors, optogenetic inhibition of ChAT^vsNAc^ neurons before intromission significantly prolonged the latency of mounting, intromission, and ejaculation (Figures 7B, 7D, 7E, and S7A; Table S23). These results suggest that ACh signaling in the vsNAc is required for the initiation of mounting, intromission and ejaculation.

**Figure 7.**
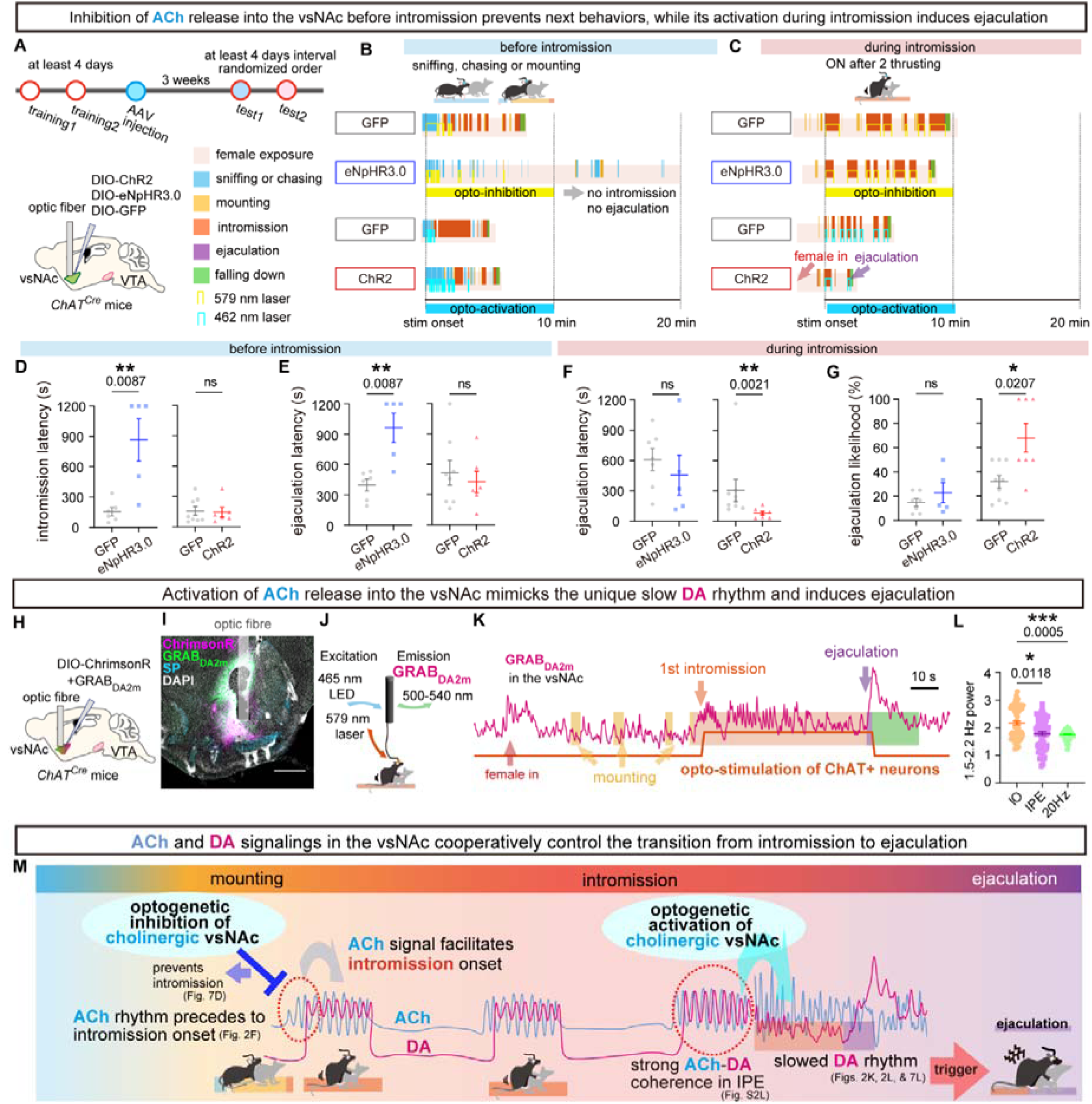
Optogenetic activation of ACh release into the vsNAc during intromission induces a specific slowdown of DA rhythm followed by immediate ejaculation. (A) Strategy for optogenetic manipulation. Experimental schedule (top) and surgery design (bottom). (B and C) Laser illustration of sexual behavior and optogenetic manipulation before intromission (B) and during intromission (C). (D and E) The effects of the optogenetic inhibition (left) and activation (right) of ChAT^vsNAc^ neurons before intromission (*n* = 6 GFP^ACh^ vs. 5 eNpHR3.0^ACh^ mice; *n* = 8 GFP^ACh^ vs. 6 ChR2^ACh^ mice) on the intromission latency (D) and ejaculation latency (E). See Table S23. (F and G) The effects of the optogenetic inhibition (left) and activation (right) of ChAT^vsNAc^ neurons during intromission (*n* = 7 GFP^ACh^ vs. 5 eNpHR3.0^ACh^ mice; *n* = 9 GFP^ACh^ vs. 7 ChR2^ACh^ mice) on the ejaculation latency (F) and ejaculation likelihood (G). See Table S24. Latency (s) = the 1^st^ target behavior onset – the 1^st^ optogenetic stimulation onset. Ejaculation likelihood (%) = ejaculation occurred or not in the test (1 or 0) / # optogenetic stimulation. (H–J) Schematic of surgery design (H), representative brain slice image (I), and recording system design (J) for simultaneous fiber photometry recording and optogenetic manipulation. (K) Representative GRAB_DA2m_ trace during sequential transitions of male sexual behaviors from female exposure to ejaculation with opto-stimulation marked by the increase of orange line. (L) Comparison of 1.5–2.2Hz power in GRAB_DA2m_ signals from 20 s before to the intromission offset among IO, IPE, and IO with optogenetic stimulation of ChAT^vsNAc^ neurons. n = 462 (IO w/o stimulation), 198 (IPE w/o stimulation), 1320 (IO with 20Hz10ms stimulation) bins from 5 mice. See Table S25. (M) ACh rhythm appears before intromission, while the optogenetic inhibition of ChAT^vsNAc^ neurons before intromission may prevent these ACh rhythms and the intromission onset. The optogenetic activation of ChAT^vsNAc^ neurons during intromission may mimic the increased coherence of ACh and DA and that ACh precedes DA more in IPE than in IO, which may trigger a slowdown of DA rhythm and ejaculation. Mean ± SEM. See also Figure S8. \**p* < 0.05, \*\**p* < 0.01, \*\*\**p* < 0.001.

Notably, optogenetic activation of ChAT^vsNAc^ neurons during intromission induced rapid ejaculation by shortening the ejaculation latency and increasing the likelihood of ejaculation (Figures 7C, 7F, and 7G; Table S24). Moreover, the number of mounting or intromission bouts and the total duration of mounting were also decreased (Figures S7G–S7I). In contrast, optogenetic inhibition of ChAT^vsNAc^ neurons during intromission did not affect mating behaviors (Figures 7F, 7G, S7G–S7J; Table S24). These results indicate that activation of ACh signaling in the vsNAc triggers the intromission-to-ejaculation transition.

To study how activation of ACh signaling changed DA dynamics during intromission, we injected AAV-hSyn-GRAB_DA2m_ and AAV-Syn-FLEX-ChrimsonR into the vsNAc of *ChAT^Cre^* mice and monitored DA dynamics by fiber photometry during optogenetic stimulation of ChAT^vsNAc^ neurons (Figures 7H–7J). Interestingly, optogenetic activation of ChAT^vsNAc^ neurons during intromission resulted in a decrease in the 1.5–2.2 Hz power of the DA rhythm followed by a brief suppression of DA release, which resembled the characteristic slowdown of DA rhythm just before ejaculation during IPE (Figures 7K and 7L; Table S25). We observed stronger coherence between ACh-DA around IPE onset than those around IO onset (Figures S2L and 7M). Taken together, these results suggest that the unique strong ACh-DA coherence in the vsNAc around the onset of IPE leads to an enhanced ACh release, which induces a slowdown of DA rhythm followed by immediate ejaculation (Figure 7M). This suggests a different ACh-DA modulatory mechanism such as via mAChRs in IPE compared with IO (see discussion).

## Discussion

### Dual ACh-DA dynamics in the vsNAc drive sequential transitions of male copulatory behaviors

The neural mechanisms that regulate the sequential transitions of male copulatory behaviors, such as mounting, intromission, and ejaculation, remain unclear. Previous studies have identified specific brain regions that control mounting (Bayless et al., 2023; Karigo et al., 2021; Sano et al., 2013; Wei et al., 2018), but the brain regions that regulate intromission and ejaculation are yet to be elucidated. Here, we identified for the first time the vsNAc as a critical brain region where dual ACh-DA dynamics align with and coordinate the sequential transitions from intromission to ejaculation. In particular, the vsNAc exhibits a unique signature of 1.5–2.2 Hz dual ACh-DA rhythms that are concurrent with the pelvic thrust rhythm during intromission. The ACh rhythm, but not DA rhythm, appears before the onset of intromission and persists during the short pauses of pelvic thrusting during intromission. These observations suggest that ACh rhythm functions as the primary driver for generating the dual ACh-DA rhythms during intromission. We proposed that dual ACh-DA rhythms play a crucial role in the maintenance of intromission possibly by reinforcing the pelvic thrust rhythm. By optogenetic manipulations of ACh and DA release in the vsNAc, we showed that ACh signaling promoted the initiation of intromission, whereas DA signaling was important for sustaining intromission through the suppression of D2R^vsNAc^ neurons. Furthermore, optogenetic activation of ChAT^vsNAc^ neurons during intromission elicits immediate ejaculation by triggering a slowdown of DA rhythm–a specific activity signature that typically precedes ejaculation during IPE. Taken together, these results demonstrate that dual ACh-DA dynamics harmonize in the vsNAc to drive the sequential transitions of male copulatory behaviors from intromission to ejaculation.

### Dual ACh-DA rhythms are generated by reciprocal regulations between ACh and DA release

Both ChAT^vsNAc^ neurons and DA^VTA➔vsNAc^ axons, but not DA^VTA^ somata, exhibited rhythmic neuronal activity during intromission, suggesting that the dual ACh-DA rhythms are regulated locally by mutual interaction between ChAT^vsNAc^ neurons and DA^VTA➔vsNAc^ axons. Consistent with this idea, we demonstrated reciprocal regulation between ACh and DA release mediated by nAChR and D1R or D2R signaling in the NAc brain slices using ex vivo (GRAB or GCaMP) imaging combined with optogenetic and pharmacological manipulations. Computational modeling suggests that the dual ACh-DA rhythms in the vsNAc are generated by reciprocal ACh-DA regulations via D2R signaling in cholinergic neurons and D2R and nAChR signaling in dopaminergic axon terminals. Accordingly, the shRNA-mediated knockdown of ChAT and D2R expression, but not D1R, in the vsNAc diminished the probability of sexual behaviors such as intromission and ejaculation. Although we primarily focused on nAChR, D1R, and D2R to simplify our modeling of the dual ACh-DA rhythms, we could not exclude a potential role for mAChRs in the generation of dual ACh-DA rhythms during intromission. A recent study also reported dual ACh-DA (∼2 Hz) rhythms in the dorsal striatum during reward delivery, but these rhythms were not generated by direct interaction between ACh and DA release within the striatum (Krok et al., 2023). The inhibition of ACh via D2R in the dorsal striatum have been implicated in locomotive and reward behaviors (Gritton et al., 2019; Chantranupong et al., 2023; Krok et al., 2023), suggesting that the dual ACh-DA rhythms can be generated by different mechanisms to regulate diverse biological processes.

### Distinct functions of D1R and D2R signaling during male sexual behaviors

Our finding that D1R^vsNAc^ and D2R^vsNAc^ neurons, which are mostly distinct neuronal populations (Figures 6A–6C), were both suppressed during intromission is consistent with the proposed “threshold” hypothesis for D1R and D2R-mediated DA signaling during sexual behaviors (Bazzett et al., 1991; Hull et al., 1992). Since D2R has a higher affinity than D1R (Richfield et al., 1989), DA signaling may preferentially cause the D2R-mediated suppression of D2R^vsNAc^ neurons, while D1R^vsNAc^ neurons are not activated during intromission (Figure 6R). Alternatively, it is plausible that suppression of D2R^vsNAc^ neurons may indirectly lead to active suppression of D1R^vsNAc^ neurons through unknown mechanisms. Importantly, optogenetic activation of D2R^vsNAc^ neurons either before or during intromission rapidly stopped the sexual behaviors. It is plausible that activation of D2R^vsNAc^ neurons is associated with aversive emotions (Lin et al., 2021; Soares-Cunha et al., 2016), which may function as an “off” switch to pause male sexual behaviors. In our computational modeling, we also analyzed the behavior of each candidate receptor by estimating its *RS* and *RE* (Figures 4E and 4F). Notably, the *RS* (signaling speed) of D2R was faster than that of D1R in cholinergic neurons, suggesting that the hypothesis that D2R^vsNAc^ neurons function as a behavioral on/off switch is reasonable. Thus, the DA-mediated suppression of D2R^vsNAc^ neurons is necessary for sustaining male sexual behaviors such as intromission.

The highest level of DA release was observed at the onset of female encounter or during ejaculation, which might cause the activation of D1R^vsNAc^ neurons to regulate the initiation and termination of male sexual behaviors. Optogenetic activation of D1R^vsNAc^ neurons during intromission increased the total mounting duration by inducing repetitive mounting behavior. It is plausible that optogenetic stimulation of D1R^vsNAc^ neurons mimics the activation of D1R^vsNAc^ neurons at female encounter and (Figure 6R), thus, resetting the sexual behaviors. Therefore, the suppression of D1R^vsNAc^ neurons during intromission ensures the forward progression of male sexual behaviors by preventing animals from returning to the previous behavioral sequences, such as mounting.

### ACh signaling regulates ejaculation timing by triggering slowdown of DA rhythm

We observed a specific slowdown of DA rhythm just before ejaculation that coincides with a big surge of DA release during IPE. Accordingly, optogenetic activation of ChAT^vsNAc^ neurons resulted in a similar slowdown of DA rhythm followed by immediate ejaculation. By contrast, optogenetic inhibition of DA^VTA➔vsNAc^ axons during intromission did not elicit ejaculation. In the striatum, cholinergic interneurons make monosynaptic or polysynaptic axo-axonic contact on DA^VTA^ axons (Cachope et al., 2012; Threlfell et al., 2012). ACh released from cholinergic interneurons can evoke action potentials at the DA^VTA^ axons and promote DA release from proximal and distant DA axons via the action potential propagation (Liu et al., 2022; Wang et al., 2014a). Therefore, optogenetic activation of ChAT^vsNAc^ neurons may have a larger effect on DA dynamics in the vsNAc than direct optogenetic manipulation of DA^VTA➔vsNAc^ axons. Moreover, optogenetic stimulation of ChAT^vsNAc^ neurons results in a high level of ACh signaling, which then leads to a brief suppression of DA signaling, both of which may be required for inducing ejaculation.

It should be noted that pharmacological inhibition of mAChRs enhanced the release of DA following 1 or 10 Hz opto-stimulation of cholinergic neurons in the NAc brain slices, suggesting that mAChRs could suppress DA release in specific contexts of ACh signaling. Optogenetic stimulation of ChAT^vsNAc^ neurons may cause a surge of ACh signaling, leading to the suppression of DA release by activating the inhibitory mAChRs on DA^VTA^ axons. Similarly, we observed that the coherence of ACh-DA rhythms between 1–4 Hz frequency was significantly higher at the onset of IPE than IO. This increased coherence of dual ACh-DA rhythms probably results in a higher level of ACh signaling, which in turn may cause the suppression of DA rhythm by activating the inhibitory mAChRs. Future studies are warranted to elucidate the detailed mechanism for why dual ACh-DA rhythms specifically results in the slowdown of DA rhythm and triggers the intromission-to-ejaculation transition during IPE, but not during IO.

To the best of our knowledge, this study is the first report of ejaculation induction by manipulating a tiny population of genetically defined neurons in freely behaving animals. Our results reveal a crucial role of ACh signaling in regulating the timing of ejaculation in mammals. Premature ejaculation (PE) is a prevalent sexual disorder affecting approximately 20–30% of sexually active men (Raveendran and Agarwal, 2021). Moreover, PE has been observed in relatively high percentages in Parkinson’s disease patients, which is thought to be the result of the medications and the disease itself, with DA signaling being the common contributing factor (Bronner et al., 2004). Selective serotonin reuptake inhibitors are also used to treat patients with PE (Althof et al., 2014). Although the exact mechanism is not yet fully understood, serotonin can affect the activity of dopamine neurons and DA release (Dremencov et al., 2009). Therefore, these drugs may affect DA dynamics and, in turn, regulate ejaculation latency. We anticipate that our findings will be a starting point for more sophisticated studies into the molecular and neural mechanisms that govern ejaculation timing and the potential development of new therapeutics for sexual dysfunctions in humans.

## Methods

### Mice

All animal experiments were performed at the International Institute for Integrative Sleep Medicine, University of Tsukuba, in accordance with the guidelines for animal experimentation. The experiments were approved by the animal experiment committee of each institute and were conducted in accordance with NIH guidelines. Mice were given food and water *ad libitum* and maintained at a temperature of 23 °C and relative humidity of 50%, with a 12-h light/12-h dark cycle. C57BL/6J mice (ID# 000664 from Charles River), male *ChAT^Cre^*mice (B6;129S6-Chattm2(cre)Lowl/J; ID# 006410, Jaxson Laboratory)(Rossi et al., 2011), male *DAT^Cre^* mice (B6.SJLSlc6a3tm1.1(cre)Bkmn/J; ID# 006660, Jaxson Laboratory)(Bäckman et al., 2006), male *Drd1^Cre^*mice (B6.FVB(Cg)-Tg(Drd1-cre)EY262Gsat/Mmucd; ID# 030989-UCD from MMRRC originally from GENSAT BAC Tg project), and male *Drd2^Cre^* mice (B6.FVB(Cg)-Tg(Drd2-cre)ER44Gsat/Mmucd; ID# 032108-UCD from MMRRC originally from GENSAT BAC Tg project) were used in this study. The ages of the mice used for the experiments were 8 to 32 weeks.

### Viruses

The plasmids of AAV-hSyn-DIO-GCaMP6s (serotype 9), AAV-Syn-FLEX-rc[ChrimsonR-tdTomato] (serotype 1), AAV-hSyn-DIO-EGFP (serotype 9), AAV-EF1a-double floxed-hChR2(H134R)-EYFP-WPRE-HGHpA (serotype 9), AAV-Ef1a-DIO-EYFP (serotype 9), and AAV-Ef1a-DIO eNpHR3.0-EYFP (serotype 9) were purchased from Addgene. The plasmids of AAV[shRNA]-CMV>mCherry-U6>Scramble_shRNA (serotype 9), AAV[shRNA]-mCherry-U6>mChat[shRNA#1] (serotype 9), and AAV[shRNA]-mCherry-U6>*Drd1*[shRNA#1] (serotype 9) were purchased from vector-builder. AAV-H1-shScramble-4-CMV-mCHERRY (serotype 10) and AAV-H1-shD2R-4-CMV-mCHERRY (serotype 10) were shared by the University of Tsukuba. The plasmids of AAV-hSyn-GRAB_DA2m (GRAB_DA2m_) (serotype 9), AAV-hSyn-GRAB_ACh3.0 (GRAB_ACh3.0_) (serotype 9), and AAV-hSyn-rDA2m (GRAB_rDA2m_) (serotype 9) were provided by Yulong Li. All AAVs were prepared at the University of Tsukuba. AAV titres were 9.79 × 10^11^ –6.24 × 10^14^ genomic copies/mL.

### Stereotaxic surgery

AAVs were produced using the triple-transfection helper-free method and purified as previously described(Lazarus et al., 2011). The titers of AAVs were determined by qPCR and purified viruses were stored in aliquots at -80°C. Details of AAVs are listed in Table S26. Adult male mice (aged > 8 weeks) were stereotactically injected with AAVs and implanted with optical fibers after at least one week post AAV injection. The animals were anesthetized with isoflurane (2–4%) and placed in a stereotaxic frame (David Kopf Instruments, Tujunga, CA). Viruses were stereotactically delivered to the brains of mice at 2–5 months of age as described previously(Inoue et al., 2019). In brief, virus was delivered unilaterally (0.2 μL of AAV-hSyn-GRAB_DA2m_, AAV-hSyn-DIO-GCaMP6s, AAV-hSyn-GRAB_ACh3_, and AAV-hSyn-GRAB_rDA2m_) or bilaterally (0.3 μL of AAV-Syn-FLEX-rc[ChrimsonR-tdTomato], AAV-hSyn-DIO-EGFP, AAV-EF1a-double floxed-hChR2(H134R)-EYFP-WPRE-HGHpA, AAV-Ef1a-DIO EYFP, AAV-Ef1a-DIO eNpHR3.0-EYFP. Immediately after surgery, mice were placed on a heated pad individually and then returned to their home cage following recovery from anesthesia. Animals were allowed at least 2 weeks of recovery in a reversed light/dark room following surgery prior to being tested in behavioral assays.

For retrograde tracing, we then injected 0.3 μL of the retrograde tracer cholera toxin subunit B (CTB) conjugated with AlexaFluor-594 and AlexaFluor-488 (Thermo Fisher Scientific) into the msNAc and vsNAc. Mice were perfused 5 days after CTB injection. Whole brain sections were prepared.

### Immunohistochemistry

For histological analysis, animals were perfused with 4% paraformaldehyde, and the brains were dissected and post-fixed overnight in 4% paraformaldehyde. Brains were sectioned at 60 μm thickness using a vibrating microtome (Leica) and immunostaining was performed as described previously(Liu et al., 2021a). In brief, sections were collected in PBS and were treated with permeabilization solution (1% Triton in PBS) at room temperature for 3 h, followed by incubation with the blocking buffer (10% Blocking One [Nacalai Tesque] with 1% Triton X-100 in PBS), and incubated overnight at 4 °C using primary antibodies in the blocking buffer. After washing with PBS three times, brain slices were incubated with secondary (second) antibody in blocking buffer at 4 °C overnight. The slices were stained with NeuroTrace fluorescent Nissl stain (Invitrogen, N-21279) or 4’, 6-diamidino-2-phenylinodole (DAPI) (Dojindo, D523), washed, mounted, and coverslipped.

Primary antisera used were chicken anti-GFP (Novus Biologicals; 1:4,000), rat anti-GFP (Nacalai Tesque; 1:2,000), rat anti-Substance P (Merck & Co., Inc.; 1:500), rabbit anti-Substance P (ImmunoStar Inc.; 1:1500), goat anti-mCherry (Sicgen; 1:2,000), rat anti-mCherry (Thermo Fisher Scientific; 1:500), goat anti-ChAT (Merck & Co., Inc.; 1:500), mouse anti-TH (Santa Cruz Biotechnology; 1:250), sheep anti-Digoxigenin-POD (Roche Applied Science; 1:500), and sheep anti-Fluorescein-POD (Roche Applied Science; 1:2,000).

Secondary antisera used were as follows: Alexa Fluor 488 donkey anti-rat (Jackson ImmunoResearch; 1:500), Alexa Fluor 488 donkey anti-rabbit (Jackson ImmunoResearch; 1:500), Alexa Fluor 488 donkey anti-chicken (Jackson ImmunoResearch; 1:500), Alexa Fluor 647 donkey anti-rat (Jackson ImmunoResearch; 1:500), Alexa Fluor 647 donkey anti-rabbit (Jackson ImmunoResearch; 1:500), Alexa Fluor 647 donkey anti-goat (Jackson ImmunoResearch; 1:500), Alexa Fluor 647 donkey anti-goat (Thermo Fisher Scientific; 1:1,000), Cy3 donkey anti-rat (Jackson ImmunoResearch; 1:500), Cy3 donkey anti-rabbit (Jackson ImmunoResearch; 1:500), Cy3 donkey anti-goat (Jackson ImmunoResearch; 1:500), and Cy3 donkey anti-guinea pig (Jackson ImmunoResearch; 1:500).

### In situ hybridization

The cDNA fragments of mouse *Drd1*, *Drd2*, *Chrm1*, *Chrm2*, *Chrm3*, *Chrm4*, and *Chrm5* were amplified by PCR using antisense primers containing the T7 promoter sequence. In vitro transcription was performed with PCR-amplified templates using T7 RNA polymerase (Hoffman La Roche AG, Basel, Switzerland) to synthesize antisense RNA probes (Table S27). Two-color in situ hybridization was performed as previously described(Ishii et al., 2017). Briefly, the mice were deeply anesthetized with isoflurane and transcardially perfused with ice-cold 10% sucrose in Milli-Q water, followed by ice-cold 4% PFA. The collected brain samples were post-fixed with 4% PFA at 4 °C overnight, followed by displacement with 30% sucrose in PBS containing 0.1% diethylpyrocarbonate at 4 °C overnight. Coronal brain sections (40 μm) were prepared using a cryostat (Leica Biosystems, Nussloch, Germany). Brain sections were treated with proteinase K (Roche), acetylated, and then incubated with hybridization buffer containing antisense RNA probes at 60 °C for 16 h. After stringent washing, the brain sections were incubated with peroxidase (POD)-conjugated anti-Fluorescein antibody (Roche Applied Science, Germany; 1:2,000) or POD-conjugated anti-Digoxigenin antibody (Roche Applied Science, Germany; 1:500) at 4 °C overnight. To sequentially use a POD-conjugated antibody, a 2% sodium azide solution was used to inactivate HRP. TSA systems (TSA-FITC and TSA-Biotin) and DyLight™ 550 conjugated NeutrAvidin Protein (Thermo Fisher Scientific; 1:500) were used to visualize mRNA signals.

HCR^TM^ RNA-FISH was performed using reagents from Molecular Instruments. In brief, the sections were incubated in 70% Ethanol/PBS at 4 °C for 1 h. After washing in DEPC-PBS, the sections were treated with proteinase K (Roche Applied Science, Germany) at 37°C for 30 min. The sections were then incubated in probe hybridization buffer for 30 min at 37 °C for 30 min, followed by incubation with a combination of D1R and D2R probes (Molecular Instruments) overnight at 37 °C. After washing in HCR probe wash buffer and 5xSSCT, the sections were then incubated overnight at 25 °C with appropriate hairpins conjugated with Alexa Fluor 488 and 647 to visualize hybridization signals. After washing in 2× SSC, the sections were stained with anti-ChAT antibody using typical immunostaining method.

### Analysis of the brain slice images

Imaging of brain sections was conducted using a LSM800, NanoZoomer XR (Hamamatsu Photonics, Hamamatsu City, Japan) and Axio Zoom V16 on-axis zoom microscope (Carl Zeiss AG, Oberkochen, Germany). To confirm the AAV-infected sites in the fiber photometry (Figures 1, 2, 6, 7, S1, S2, S6, and S7), RNAi silencing (Figures 4I–4K, and S4J), and optogenetics experiments (Figures 5–7, and S5–S7), coronal brain sections were immunostained for substance P (SP) (Voorn et al., 1989). The recording site was identified based on fiber tracking of the brain histological section sample. Each subregion of the NAc was defined as follows. While the cNAc shows a low level of SP expression, the msNAc and vsNAc are characterized by high levels of SP expression. Moreover, the msNAc is located on the medial side of the cNAc, and the vsNAc is sandwiched between the cNAc and the dense SP axons (ventral pallidum) (Figure 1C).

### Quantification of mAChR1-5, Drd1, and Drd2 on ChAT^vsNAc^ and DA^VTA^ neurons in the brain slice images

To confirm the expression patterns of ChAT and dopamine receptors (*Drd1* and *Drd2*) in the NAc (Figure 3F), ChAT and *mAChR1*–*5* (*mAChR1*, *mAChR2*, *mAChR3*, *mAChR4*, *mAChR5*) in the vsNAc (Figure S3C), TH and *mAChR1*–*5* in the VTA (Figure S3D), and dopamine receptors in the NAc (Figure 6A), we manually annotated the target marker^+^ somata using a multi-point function and extracted the NAc subregions by polygon selection in ImageJ/Fiji. The number of signals in each subregion was counted using schemes implemented in Python 3.7.2. Expression ratios were calculated based on the number of regions of each target marker. The *Drd1* or *Drd2* ratio in the ChAT^vsNAc^ neurons was calculated as the target marker (*Drd1* or *Drd2*)^+^ChAT^+^ / ChAT^+^ (%) (Figure 3G). To calculate the *mAChR1-5* ratio in the ChAT^vsNAc^ neurons, the target marker (*mAChR1-5*)^+^ChAT^+^ / ChAT^+^ (%) was calculated (Figure S3E). To calculate the *mAChR1-5* ratio in DA^VTA^ neurons, the target marker (*mAChR1-5*)^+^TH^+^ / TH^+^ (%) was calculated (Figure S3F). The number of neurons / (100 μm)^2^ in each subregion of the NAc were calculated as the total number of the target marker (*Drd1* or *Drd2*) / the area of the region (100 μm)^2^ (Figure 6B). The mean values for each mouse are shown (Figures 3G, S3E, S3F, and 6B).

### The shRNA-mediated knockdown in the vsNAc

AAV[shRNA]-CMV>mCherry-U6>Scramble_shRNA, AAV[shRNA]-mCherry-U6>mChat[shRNA#1], AAV[shRNA]-mCherry-U6>*Drd1*[shRNA#1], AAV-H1-shScramble-4-CMV-mCHERRY, AAV-H1-shD2R-4-CMV-mCHERRY) at 100 nL/min with a Hamilton syringe using a micropump. To calculate the knockdown ratio of target marker by shRNA, we prepared three types of images in which shRNA (mCherry), the target marker (GFP: *Drd1, Drd2*, or ChAT), and DAPI were observed around the vsNAc. The target^+^ (mCheery^+^, GFP^+^, or DAPI^+^) somata were extracted from each image using OpenCV 4.5.5. First, cv2.medialBlur with a kernel size of 9 was applied to the image to reduce noise. Next, the histogram of the image was calculated using cv2.calcHist, and the peak of the histogram was obtained using scipy.signal.find_peaks with a distance of 5. V_pk_ was set to the pixel value corresponding to the maximum peak. The image was binarized with a threshold of V_pk_ minus 10, and the contours of the target^+^ somata were extracted by applying cv2.findContours to the binarized image. The mode and method were set to becv2.RETR_EXTERNAL and cv2.CHAIN_APPROX_SIMPLE, respectively. We selected the contours of DAPI and the target markers with areas between 10 and 200 in the images. The knockdown ratios were calculated from the region numbers of the target markers, shRNA, and DAPI. The shRNA knockdown ratio was calculated as the target marker (*Drd1, Drd2* or ChAT)^+^shRNA^+^ / DAPI^+^shRNA^+^ (%). The mean values for each mouse are shown in Figures 4I– 4K.

### Sexual behavior test

#### Hormone priming

Stimulus females in estrus were induced as described previously(Sano et al., 2013). C57BL/6J female mice were ovariectomized at > 8 weeks of age. One week after the surgery, estradiol benzoate dissolved in sesame oil (10 μg/0.1 mL; EB) was administered to all female mice as a stimulus. More than 1 d after the first EB injection, all female mice were hormonally primed with subcutaneous injections of EB 48 and 24 h before testing, and they received progesterone dissolved in sesame oil (500 μg/0.1 mL) 4–6 h before testing to ensure high sexual receptivity. Male mice were exposed to all female mice in their home cages. Female mice were examined for ejaculation using a copulatory plug. Female stimuli that experienced ejaculation one time were used in the sexual behavior test. Male mice were exposed to a different receptive female mouse from that used in the previous test.

#### Behavioral analysis

Sexual behaviors, including mounting, intromission, ejaculation, and falling, were analyzed based on videos obtained using cameras. The video frames were manually annotated with these sexual behaviors using BORIS (Friard and Gamba, 2016). The duration and number of mountings, intromissions, and ejaculations were recorded for each mouse. The latency of each behavior was calculated by subtracting the female onset from the onset of each behavior. For optogenetic manipulation before intromission, appetitive behaviors, including sniffing and chasing a female mouse, were manually annotated (Figures 5E, 6H, and 7B).

#### Visualization of the intromission thrust movements

The body regions of two male and female mice during intromission were extracted using Python 3.7.2 and OpenCV 4.5.5. to visualize their thrusting movements. First, video frames with intromission behaviors were extracted from the original videos, and the intromission frames were then converted into grayscale images. Next, the contrast-limited adaptive histogram equalization (CLAHE) operator was applied to the grayscale images to improve their contrast (clipLimit = 2.0, tileGridSize = (3, 3)). The histogram of each CLAHE image was calculated using cv2.calcHist. The histogram had several pixel values with a height of 500 pixels. With V_500_ representing the minimum pixel value, the binarization operator was applied to the CLAHE image using threshold V_500_. Conversely, the createBackgroundSubstractorMOG2 (BSMOG2) was applied to the grayscale images, and the median smoothing filter with a kernel size of 25 was applied to the BSMOG2 images. The binarized CLAHE images were multiplied by the smoothed BSMOG2 images, and black (zero) pixels were extracted from the multiplied images as mouse body regions (Figure 2G). The Z-scored lines of mouse body region sizes and the bottom point as thrusting behavior were plotted using MATLAB (Figure 2H).

#### Fiber photometry experiment

##### i) Fiber photometry recording

For fiber photometry imaging of neurotransmitters, AAV-hSyn-GRAB_DA2m_, AAV-hSyn-GRAB_ACh3.0_, or AAV-hSyn-GRAB_rDA2m_ was injected unilaterally into the NAc of C57BL/6J mice. For fiber photometry imaging of Ca^2+^ dynamics, AAV-hSyn-DIO-GCaMP6s was injected unilaterally into the VTA of *DAT^Cre^* mice or the vsNAc of *ChAT^Cre^* mice, *Drd1^Cre^* mice, or *Drd2^Cre^* mice. More than 1 week after virus injection, a 200-μm core, 0.50 NA optical fiber (Thorlabs and Lymyth) was implanted unilaterally into the vsNAc or VTA. The mice were allowed to recover for at least 2 weeks before the experiments.

To monitor animal behavior during fiber photometry imaging, (method 1) a web camera (Logicool HD Webcam C525r, Logitech International S.A., Switzerland) was used. To synchronize the fiber photometry recording and the video recording that captures mouse behaviors, (method 2) a USB camera (FLIR USB3.0 Vision Camera # CM3-U3-13Y3M-CS; Teledyne FLIR, Wilsonville, OR) was used. The USB camera released the shutter at a rate of 10 fps based on the Fiber Photometry Console signals. On the day of the experiment, mice were recorded in their home cages with regular chow feeding for 2 h in the dark phase. More than 10 min after the onset of recording, an estrous female mouse was introduced into the home cages of male mice. The recording was stopped more than 2 h after introducing the female. After the test, we reviewed the video footage and examined the female mouse’s vagina to identify any signs of ejaculation by the male mouse. In the two-color imaging of GRAB_ACh3.0_, and GRAB_rDA2m_, signals were collected from three sexual behavior tests (at least 4 d interval) (Figures 2A–2K, and 4A–4G).

For DA, ACh, or GCaMP imaging, the fluorescence emitted by GRAB_DA2m_, GRAB_ACh3.0_, or GCaMP was collected under a blue LED (CLED465), and the fluorescence emitted by GRAB_rDA2m_ was collected under a green LED (CLED560). The GRAB_DA2m_, GRAB_ACh3.0_, and GCaMP signals were recorded at an acquisition rate of > 10 Hz, and those of GRAB_ACh3.0_ and GRAB_rDA2m_ were recorded at an acquisition rate of 20 Hz using Doric Neuroscience Studio (version 5. 4. 1. 5, Doric Lenses Inc.). The LED output intensity was adjusted so that the light excitation of the optical fiber cable was approximately 30 μW/mm^2^. The signals of GRAB_DA2m_, GRAB_ACh3.0_, and GCaMP between 500–540 nm and those of GRAB_rDA2m_ between 580–680 nm were recorded.

##### ii) Data analysis for fiber photometry signals

All fiber photometry signals, except for the two-color imaging of GRAB_ACh3.0_, and GRAB_rDA2m_, were transformed into 10 Hz by a MATLAB interp1 function to align with the behavioral events. As peri-event time plots (PETP), To analyze the signal dynamics around the onset and offset of each sexual behavior, we extracted the signal intensities for 20 s before and after the behavioral events (for a total of 40 s). The ΔF/F ratio was calculated by dividing the signal intensity in each frame by the mean value for 20 s before the behavioral events. The ΔF/F ratio was z-scored against the extracted 40-s signal intensities. To compare the intensity changes of each signal, mean ΔF/F every 10 s in each behavioral event, in which the baseline was the mean for the first 10 s were plotted (Figures 1E–1G, S1G, S1H, S1J, S2E, S2H, S6A, and S6B).

To analyze the DA rhythm (Figures 1H–1J, 2L, and S2M) or dual ACh-DA rhythms (Figures 2E, 2J, and 2K) during intromission, we calculated the power spectrum using MATLAB FFT with a 3 s window shifted every 0.3 s or 0.25 s, respectively. To show that ACh rhythm, but not DA rhythm, started before intromission, we analyzed the sum of the power spectrum of the 1.5–2.2 Hz band of ACh/DA rhythms every 2 s observed from 10 s before to 10 s after the intromission onset (Figure 2F). To analyze the DA rhythm during intromission, we collected the sum of the power spectrum every 0.2 Hz band during intromission behavior (Figures 2L and S2M).

To analyze the coherence between GRAB_ACh3.0_ and GRAB_rDA2m_, first we prepared the extracted signals around intromission. The GRAB_ACh3.0_ and GRAB_rDA2m_ signals were extracted 20 s before the intromission onset to 10 s before the intromission onset and 10 s before the intromission offset to 10 s after the intromission offset. These two signals were combined, and 50 s length signals were prepared. Using these 50 s length signals, MATLAB wavelet coherence was applied. Average values of the wavelet coherence over all intromission (Figure 2C), IO, and IPE (Figure S2K) were were visualized by heatmap plot. To compare each frequency of coherence between GRAB_ACh3.0_ and GRAB_rDA2m_ around the intromission, we calculated the mean of the wavelet coherence around <1 Hz, 1–4 Hz, and >4 Hz every 10 s (Figures 2D and S2J). To compare the coherence between GRAB_ACh3.0_ and GRAB_rDA2m_ around the IO and IPE, the mean of the wavelet coherence in IO and IPE around 1–4 Hz every 5 s were calculated (Figure S2L).

To visualize ACh and DA dynamics in the vsNAc during each thrust in the intromission phase, we extracted the time point that showed the bottom of the area of the mouse body, which was assumed to be the pelvic thrusting behavior. Based on the bottom time point, we extracted GRAB_ACh3.0_ and GRAB_rDA2m_ around the thrusting behavior ± 0.5 s (Figures 2G and 2H).

To analyze the GRAB_DA2m_ dynamics with optogenetic activation of DA^VTA→ vsNAc^ axons using ChrimsonR, the ΔF/F ratio was calculated by dividing the signal intensity in each frame by the mean value for 30 s before the stimulation onset (Figures 5C, S5A, and S5B). To compare the GRAB_DA2m_ intensity before and after the stimulation, mean ΔF/F -5∼0 s and 0∼5 s around the stimulation onset were plotted (Figure 5C).

To analyze the GRAB_DA2m_ dynamics with optogenetic activation of ChAT^vsNAc^ neurons using ChrimsonR, we calculated the power spectrum using MATLAB FFT with a 3 s window shifted every 0.3 s and collected the power spectrum of the 1.5–2.2 Hz band observed from 20 s before the ejaculation onset to the ejaculation onset during intromission (Figure 7L).

### Ex vivo imaging and optogenetics

We used 8–21-week-old *ChAT^Cre^* or *ChAT^Cre^*; *DAT^Cre^* male mice for imaging and optogenetic experiments during acute brain histological section preparation. These mice were injected with the following AAV vectors into the target brain areas at least 3 weeks prior to the experiments: (i) for imaging the DA response to photostimulation of cholinergic neurons in the vsNAc, AAV-Syn-FLEX-ChrimsonR-tdTomato and AAV-hSyn-GRAB_DA2m_ were injected into the vsNAc of *ChAT^Cre^* mice (Figures 3B and 3C); (ii) for imaging Ca^2+^ in cholinergic neurons in response to photostimulation of dopaminergic axons from the VTA, AAV-Syn-FLEX-ChrimsonR-tdTomato and AAV-hSyn-DIO-GCaMP6s were injected into the VTA and vsNAc of *ChAT^Cre^*;*DAT^Cre^*double transgenic mice, respectively (Figures 3D and 3E); (iii) for imaging the ACh response to photostimulation of cholinergic neurons in the vsNAc, AAV-Syn-FLEX-ChrimsonR-tdTomato and AAV-hSyn-GRAB_ACh3.0_ were injected into the vsNAc of *ChAT^Cre^* mice (Figures 3H–3K). Acute brain histological sections were prepared using a slightly modified N-methyl-d-glucamine (NMDG) protective recovery method(Takahashi et al., 2022; Ting et al., 2014). Briefly, mice were anesthetized with isoflurane and then cardiovascularly perfused with ice-cold NMDG-artificial cerebrospinal fluid (aCSF) consisting of 93 mM NMDG, 93 mM HCl, 2.5 mM KCl, 1.2 mM NaH_2_PO_4_, 30 mM NaHCO_3_, 20 mM HEPES, 25 mM D-glucose, 5 mM Na-ascorbate, 2 mM thiourea, 3 mM Na-pyruvate, 10 mM MgSO_4_, and 0.5 mM CaCl_2_ (pH 7.4, adjusted with HCl, oxygenated with 95% O_2_/5% CO_2_). The brains were sliced to a thickness of 200 μm in ice-cold NMDG-aCSF using a vibratome (Leica VT1200S). Slices containing the striatum were incubated in NMDG-aCSF at 32 °C for 12 min and then in HEPES-aCSF consisting of 124 mM NaCl, 2.5 mM KCl, 1.2 mM NaH_2_PO_4_, 24 mM NaHCO_3_, 5 mM HEPES, 12.5 mM D-glucose, 2 mM CaCl_2_, and 1 mM MgSO_4_ (∼310 mOsm, oxygenated with 95% O_2_/5% CO_2_) at room temperature for at least 1 h. Slices were transferred into a chamber under an upright fluorescence microscope (Zeiss AxioExaminer D1, HXP120, 20x or 40x water-immersion objective lens) and submerged in perfused HEPES-sCSF at 32–35 °C. Fluorescence signals were acquired using an EM-CCD camera (Andor iXon Ultra 888, 1024 × 1024 pixels, 16 bit) and iQ software (Andor), in which green and red fluorescence were observed using a Zeiss filter set 38HE (excitation 470/40 nm, emission 525/50 nm) and Zeiss filter set 43 (excitation 545/25 nm, emission 605/70 nm), respectively. Green fluorescence included GRAB_DA2m_, GRAB_ACh3.0_, and GCaMP6s. The frame rate was 5 fps, except for experiments with Ca^2+^ imaging or bath application of SKF38398 (frame rate of 0.2 fps). For optical stimulation, an optical fiber was placed just above the target area and outside the field of view, and optical stimuli (579 nm, 10-ms pulses for 30 s, Shanghai Laser) were controlled using a function generator (Master-9, AMPI). A data acquisition system (Digidata® 1440A Low-noise data acquisition system, Molecular Devices) was employed to synchronize the timing of optical stimuli and image acquisition. This system allowed for precise monitoring of the timing of both processes.

Image analysis was performed using Fiji and R. All images were subtracted from the background signal, which was the mean fluorescence intensity outside the field of view of the objective. Bleaching was corrected by linear interpolation of the medians of fluorescence intensity for 5–10 s before and after photostimulation. For optogenetics and imaging with GRAB_DA2m_ or GRAB_ACh3.0_, the edges of the 1024 × 1024-pixel images were removed to obtain 1008 × 1008-pixel images because the edges were outside the field of view of the objective lens. A total of 196 (16 × 16) square ROIs with 63 pixels per side were created in a 1008 × 1008 image, and the mean fluorescence intensity within each ROI was calculated. ROIs without GRAB_DA2m_ or GRAB_Ach3.0_ expression (mean intensity <100) were excluded from the analysis. The mean fluorescence intensity was z-scored against the value obtained for 20 s before photostimulation. For experiments with SKF38398, bleaching was corrected using the median values for 60 s before and 120–180 s after bath application, and the fluorescence intensity was z-scored against the values for 60 s before the application. For Ca^2+^ imaging with GCaMP6s, ROIs were set on the somata of the neurons expressing GCaMP6s.

ROIs were classified using the Mann–Whitney U test, with values for 5 s before and 10 s after photostimulation for the GCaMP experiments (Figure 3E), and 60 s before and 120 s after the application of SKF38398 for the GRAB_ACh3.0_ experiments (Figure 3K).

### Computational modeling of DA and ACh dynamics

As we observed similar dynamics in GRAB and GCaMP (GRAB_DA2m_ and GCaMP6s of dopaminergic axons in the vsNAc [Figures 1 and S1I–S1K], GRAB_ACh3.0_ and GCaMP6s of cholinergic neurons in the vsNAc [Figures S2D–S2I]) by fiber photometry imaging, we hypothesized that (1) the neuronal activity of the neuronal ensemble reflects the amount of NT (neurotransmitter) released; (2) the GCaMP dynamics reflect the release probability of the NT. Based on these hypotheses, we developed a computational model to estimate the cellular environment in which neurons and receptors are involved in generating dual ACh-DA rhythms during intromission. In this model, we set neuronal assembly 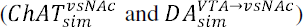, the internal activity of neurons (release probability measurement, *RPM*), NT release (*ACh_sim_* and *DA_sim_*) based on the *RPM*, receptors expressed on neurons (D1R, D2R, and nAChR), and the extracellular environment in which the NT can be released. To mimic the change of the amount of the extracellular NT over time, in this model, we assumed the three steps of cycle: (i) the activation of target neurons by the external stimuli; (ii) NT released from the target neurons; (iii) the effect of the receptor by binding NT to the receptor on the target neurons. These cycles were repeated a certain number of times and, during this period, we recorded the *RPM* and amount of *NT_sim_* released from the neurons which was determined based on the *RPM* value. Through this model, we attempted to estimate the appropriate cellular environment, including the type of receptors or neurons involved, by minimising *Diff* (*NT_sim_*, *NT_real_*), the difference between the actual ACh-DA dynamics in vivo (NT recorded by fiber photometry: *NT_real_*) and model-simulated ACh-DA dynamics (NT generated by the model: *NT_sim_*) (Figures 4A, 4B, and S4A). As *NT_real_*, we used typical rhythmic signals of in vivo GRAB_ACh3.0_ and rGRAB_DA2m_ during intromission from the data shown in Figures 2A–2K.

Hypothetical neurons and receptors used in this simulation were identified based on ex vivo imaging (Figure 3), the in situ hybridization results (Figures S3C–S3F), and previous reports (De Mei et al., 2009). As neurons and receptors, we placed D1R and D2R in 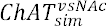 neurons, as well as D2R and nAChR in 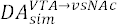 axon terminals. From our ex vivo imaging, we found that inhibitory mAChR may function as an autoreceptor (Figures 3B and 3C). However, since all subtypes of *mAChR* were expressed on ChAT^vsNAc^ and DA^VTA^ neurons (Figure S3C–S3F), to simplify our modeling, we focused on D1R, D2R, and nAChR receptors in this experiment (Figure 4A).

#### (i) Activation of target neurons by the external stimuli

To mimic the cellular environment that neurons are activated by the external stimuli, in this modeling, we assumed that the *RPM* which reflects the intensity of neuronal activity was changed by the external stimuli or the effect of the receptor expressed on the neurons. *Activation* includes the spontaneous activity of the target neurons and stimulation of neurons by substances other than ACh and DA. Here, we defined that the *Activation* continuously adds a certain value to the target neuron. Thus, the value added as an external stimulus to the *RPM* of the target neurons every cycle was constant.

#### (ii) Release of NT from the target neurons

To mimic the manner of NT release, we defined that the volume of *NT_sim_* released to the extracellular environment from the neurons was determined by the *RPM* value of the neurons. In this model, the volume of *NT_sim_* released is equivalent to the *RPM* value if *RPM* is positive and 0 if *RPM* is negative (Figure 4B). Additionally, we defined the diffusion function of DA to mimic the in vivo extracellular DA diffusion mechanism (Liu et al., 2021b). The time course and intensity of the DA diffusion were investigated using this model (Figures S4B–S4D).

#### (iii) Effect of the receptor by binding NT to the receptor on the target neurons

Next, we considered the strategy to mimic the behavior of nAChR, D1R, and D2R. As reported in previous studies, DA receptors are GPCRs that are thought to respond more slowly than nAChR, the ionotropic receptor (Cruz et al., 2004; Fucile, 2004; Galzi et al., 1992; Greif et al., 1995; Neve et al., 2004). Incorporating these insights, we defined the characteristics of each receptor using two variables: *RE* (receptor efficacy), the effect intensity which changes RPM; *RS* (receptor speed), the latency where the *Receptor* function affects *RPM* from the time that *NT_sim_* binds to the receptor. In this experiment, to simplify the setting, we set the nAChR related variables as constant values (nAChR’s *RE*=1, and the *RS*=1), and estimated the characteristics of D1R and D2R by exploring their *RE* and RS. *RE* of D1R and D2R was searched over a range of positive values, and their *RS* was searched over a range of natural numbers.

For example, when *ACh_sim_* binds to the nAChR expressed on the 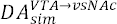 axon terminals, there is an increase in the *RPM* of the 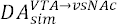 axon terminal *ACh_sim_* × 1 (nAChR’s *RE*=1) one step later (nAChR’s *RS*=1). Since the D1R receptor is an excitatory GPCR, we defined its operation as follows: when D1R receives DA_sim_, “D1R’s *RS*” step later, the *RPM* is increased to DA_sim_ × “D1R’s RE.” By contrast, we defined operation of the D2R receptor, an inhibitory GPCR, receiving DA_sim_ as “D2R’s *RS*” step later, *RPM* is decreased by a factor of DA_sim_ × “D2R’s *RE*”. With these receptors’ settings, we conducted computational modeling to estimate the *RE* and *RS* values.

To estimate receptor efficacy (*RE*) and receptor speed (*RS*) of DA receptors, we explored *RE* and *RS* values which minimise *Diff* (*NT_sim_*, *NT_real_*), the difference between *NT_sim_* and *NT_real_* using Bayesian optimisation (Figure 4B). We used Optuna (Akiba et al., 2019), an automatic hyperparameter optimization software framework, to determine the best values for *RE* and *RS* using *Diff* (*NT_sim_*, *NT_real_*) as an indicator. We used the following values as a fixed value throughout exploration. The value added by *Activation* is 1. The number of trials conducted to estimate *RE* and *RS* of D1R and D2R was approximately 1,000 to 10,000, depending on the experimental design. In each experimental design, out of these trials, the one representing the smallest *Diff* (*NT_sim_*, *NT_real_*), the condition where *NT_sim_* and *NT_real_* are most similar, was used to define *RE* and *RS* of D1R and D2R. To express the effect speed of each receptor in seconds, the *RS* of each receptor and DA diffusion step were divided by 200, the sampling rate (Figures 4E and S4D).

## Experiment 1: Performance comparison by the different activation patterns

To prove which neurons should be activated in the generation of dual ACh-DA rhythms, we compared the *Diff(NT_sim_,NT_real_*) under the following three conditions: 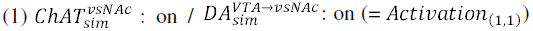; 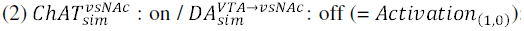; 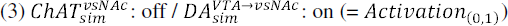 (Figures 4D and S4A).

## Experiment 2: Estimation of *RE* and *RS* of dopamine receptors

To reveal the *RE* and *RS* values of D1R and D2R generating dual ACh-DA rhythms, we estimated the *RE* and *RS* values under the *Activation*_(1,1)_ with full receptors condition (D1R and D2R on 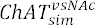, and D2R and nAChR on 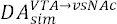). We focused on trials showing the converged *Diff* (*NT_sim_*, *NT_real_*) because otherwise the trial indicated the failure of the variable search (Figures 4E and 4F).

## Experiment 3: Performance comparison by deletion of each candidate receptor

To find which candidate receptor is important for the generation of dual ACh-DA rhythms, we estimated the *RE* and *RS* values under five conditions which delete a target *receptor*, deletion of 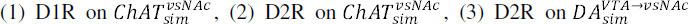, (4) D2R on both 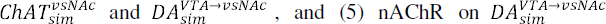. Then, we compared the *Diff* (*NT_sim_*, *NT_real_*) among these five conditions and the condition with full receptor using the estimated each candidate *RE* and *RS* of D1R and D2R (Figures 4G, S4E, and S4F).

### Gene knockdown experiments

#### Sexual behavior test with shRNA treatment

To silence the gene expression of ChAT, D1R, or D2R, AAV-CMV-mCherry-U6-shmChAT, AAV-CMV-mCherry-U6-shD1R, or AAV-H1-shD2R-CMV-mCherry was injected into the vsNAc of C57BL/6J male mice. In the control group, AAV-H1-shScramble-CMV-mCherry or AAV-CMV-mCherry-U6-shScramble was injected into the vsNAc of C57BL/6J male mice. Three weeks (± 1 day) following AAV injection, male mice underwent the sexual behavior test once a week (± 1 day) for 3 weeks. Male mice were relocated to the test room 30 min before starting the test. After the acclimation period, a female stimulus mouse was introduced into the home cage of a male mouse. Video recordings were conducted over the next 2 h. After the test, we reviewed the video footage and inspected the female mice to determine whether male mice exhibited sexual behaviors. The intromission and ejaculation rates were then calculated based on the frequency of these behaviors observed across the three tests (Figures 4H, 4L, and 4M). The average latency, the number of bouts, and total duration of each behavior were calculated across the tests (Figures S4G– S4M).

#### Rotarod test with shRNA treatment

A rotarod test was conducted to confirm the motor coordination of mice expressing shRNA in the vsNAc. To silence the gene expression of ChAT, D1R, or D2R, AAV-CMV-mCherry-U6-shmChAT, AAV-CMV-mCherry-U6-shD1R, or AAV-H1-shD2R-CMV-mCherry was injected into the vsNAc of C57BL/6J male mice. For the control group, AAV-H1-shScramble-CMV-mCherry or AAV-CMV-mCherry-U6-shScramble was injected into the vsNAc of C57BL/6J male mice. More than 3 weeks after the AAV injection, male mice were moved to the test room. After more than 15 min, we initiated a habituation process for the rotarod machine. During this phase, mice were placed in the stationary state of the rotarod machine for 2 min. After acclimatization, the male mice were allowed to practice on a rotarod for 2 min (12 rpm). Subsequently, we conducted three rotarod tests (each gradually increasing from 5 to 40 *RPM* for 300 s). We recorded the latency period before the mice were dropped from the rotarod machine. Between each test, the mice were allowed a 5-min break. The median value was plotted as an effect of motor coordination (Figure 4N).

#### Optogenetics experiments

To activate DA^VTA→^ ^vsNAc^ axons, ChAT^vsNAc^ neurons, D1R^vsNAc^ or D2R^vsNAc^, AAV-Ef1a-DIO-ChR2-EYFP, or AAV-hSyn-DIO-EGFP was injected bilaterally into the VTA of *DAT^Cre^* mice or the vsNAc of *ChAT^Cre^* mice, *Drd1^Cre^* mice, or *Drd2^Cre^* mice. To inhibit DA^VTA→vsNAc^ axon terminals or ChAT^vsNAc^ neurons, AAV-Ef1a-DIO-eNpHR3.0-EYFP or AAV-Ef1a-DIO-EGFP was injected bilaterally into the VTA of *DAT^Cre^* mice or the vsNAc of *ChAT^Cre^*mice. After more than 2 weeks, 200-μm core and 0.50 NA optical fibers (Thorlabs and Lymyth) were bilaterally implanted into the vsNAc. The mice were allowed to recover for at least 2 weeks before conducting the experiments. Before the tests, it was confirmed that the male mice ejaculated twice as a training. The interval between each training was at least 4 days. Ejaculation was determined by verifying the presence of a plug in the vagina of the female mouse. If male mice did not show any sexual behavior in 3 training trials, the mice were excluded.

Two tests were performed: manipulation before (20 Hz, 10 ms) and during intromission (20 Hz, 10 ms) (Figures 5D–5K, 6G–6Q, 7A–7G, S5C–S5H, S6C–S6F, and S7). The interval between each test was at least 4 days. On the test day, the mice were moved to the experimental room 1 h before the experiments in the dark phase and optical fibers were connected to their heads. The axonal terminals of dopaminergic neurons and cholinergic neurons in the vsNAc were activated (462 nm, Shanghai Laser) or inhibited (579 nm, Shanghai Laser). We created a system using an electrical board to record the stimulation timing and manage the stimulation frequency using the Arduino UNO and Doric Neuroscience Studio. Video recordings and recordings of the optogenetic stimulation schedule were initiated simultaneously. In each test, the experimenter observed the behavior of the mice and turned on the switch to shine the laser. The switch was immediately turned off if the target behavior was prevented. If the male mice did not exhibit any sexual behavior within 10 min of starting the test, the test was terminated and repeated on a different day. If the subject male mice did not ejaculate within 10 min after the onset of stimulation, we stopped the stimulation and confirmed the timing of ejaculation until 20 min after the onset of stimulation (Figure 7B). In our pilot experiment, we observed that most experienced mice (>2 times) show ejaculation within 20 min after female introduction, thus we waited for 20 min at least after the optogenetic stimulation onset. If the male mice did not show any sexual behavior within 10 min after the onset of the test, we stopped the test and retested it more than 4 d after the test. In the case that male mice did not show any sexual behavior in 5 tests, they were excluded.

For manipulation before the intromission (20 Hz, 10 ms) test, the experimenter turned on the laser while observing sniffing, chasing, and mounting. When mounting was not prevented, the mice showed intromission, the stimulation was stopped, and the timing of ejaculation was confirmed. In the other case, the target behavior was prevented within 10 min of starting the stimulation; accordingly, we stopped the stimulation and confirmed the timing of ejaculation. If the male mice did not ejaculate within 20 min of starting the stimulation, the test was terminated, and the ejaculation latency was recorded as 1,200 s.

For manipulation during the intromission tests (20 Hz, 10 ms), the experimenter turned on the laser when seeing the intromission, specifically when thrusting occurred more than twice. In cases where ejaculation was not observed within 10 min of starting the stimulation, the stimulation ceased, and we confirmed the timing of ejaculation within 20 min of starting the stimulation. The recorded ejaculation latency in such cases was 1,200 s.

The latency of each behavior was calculated by subtracting the 1^st^ optogenetic stimulation onset from the onset of each 1^st^ behavior (Figures 5I, S5C, S5D, 6M, 6O, S6C, S6E, 7D, 7E, 7F, and S7A). Epoch duration of sniffing and intromission were calculated by subtracting the optogenetic stimulation onset from the onset of each target behavior (Figures 6J, 6K, and 6N). To reveal the rebounded mounting bouts by the optogenetic activation of D1R^vsNAc^ neurons during intromission, the mounting behaviors observed during 10 min after the optogenetic stimulation were used, except for the mounting preceding intromission (Figure 6Q). Ejaculation likelihood was calculated by dividing ejaculation occurred or not in the test (1 or 0) by the number of optogenetic stimulation (Figures S5D, S5G, and 7G).

### Fiber photometry with optogenetics

To monitor DA dynamics when stimulating cholinergic neurons, AAV-Syn-FLEX-ChrimsonR-tdTomato and AAV-hSyn-GRAB_DA2m_ were injected into the VTA of *DAT^Cre^* mice (Figures 5A–5C, S5A, and S5B) and the vsNAc of *ChAT^Cre^* mice (Figures 7H–7L). Subsequently, optical fibers with a 200-μm core and 0.50 NA (Thorlabs and Lymyth) were bilaterally implanted in the vsNAc. The mice were allowed to recover for at least 2 weeks before performing the experiments. GRAB_DA2m_ recordings were conducted without optogenetics to confirm the GRAB_DA2m_ signals. After confirmation, bilateral optogenetic manipulations without fiber photometry recordings were conducted to confirm optogenetic effects on behavior. Target neurons were activated using a laser (579 nm, Shanghai Laser).

After confirming the behavioral expression, DA dynamics were recorded during optogenetic activation of cholinergic neurons in the vsNAc. In this experiment, unilaterally DA dynamics were monitored, and target neurons were activated during intromission. We created a system using an electrical board to record the stimulation timing and manage the stimulation frequency with the help of the Arduino UNO and Doric Neuroscience Studio. Fiber photometry recordings, video recordings using a USB camera (FLIR USB3.0 Vision Camera # CM3-U3-13Y3M-CS), and recordings of the optogenetic stimulation schedule were synchronized using Doric Neuroscience Studio. In each test, the experimenter observed the behavior of the mice. When observing intromission, particularly more than two intromission-thrusting movements, the experimenter turned on the switch button to shine the laser. The fluorescence emitted by GRAB_DA2m_ was recorded under a blue LED (CLED465) using a Doric fiber photometry system for DA imaging. GRAB_DA2m_ was recorded at 10 Hz using Doric Neuroscience Studio (version 5.4.1.5). The LED output intensity was adjusted such that the amount of light exiting the optical fiber cable was approximately 30 μW/mm^2^. The GRAB_DA2m_ signals within the 500–540 nm range were recorded (Figures 5A and 7J).

### Quantification and statistical analysis

Recorded videos and histological samples were analyzed blind to relevant variables, including administrated solution, genotype, prior housing condition, surgical procedure, and virus injection. Videos were manually annotated using BORIS. In particular, anogenital investigation (sniffing and chasing), mounting, repeated pelvic thrust (intromission), and ejaculation were scored for male sexual behaviors. After annotations, various parameters including latency, number, and duration were calculated for further analysis.

Data were processed and analyzed using MATLAB, R, Python 3.7.2, and GraphPad PRISM 9.4.1 (GraphPad Software, San Diego, CA). All results are expressed as mean ± SEM. Data of non-paired samples were analyzed with two-tailed non-parametric Mann–Whitney U test, Kruskal– Wallis test with Dunn’s multiple comparisons test or two-way ANOVA followed by Šídák’s post-hoc multiple comparison test. Data of paired samples were analyzed with Friedman’s ANOVA followed by Dunn’s multiple comparisons test. The significance threshold was held at α=0.05, two-tailed (not significant [ns], *p* > 0.05; *\*p* < 0.05; *\*\*p* < 0.01; *\*\*\*p* < 0.001; *\*\*\*\*p* < 0.0001). Details are available in Supplemental Table S1–S25.

## Data and code availability

Further information and requests for resources should be directed to and will be fulfilled by the Lead Contact, Qinghua Liu. This study did not generate new unique reagents. The code and software used in this study is available through Zenodo (DOI: 10.5281/zenodo.12748186, URL: https://zenodo.org/records/12748186).

## Acknowledgements

We thank M. Jing for sharing reagents; A. Yoshida, M. Endo, N. Fujii-Sekine, T. Lou, and A. Watanabe for the behavioral analysis; Editage (www.editage.com) for English language editing; K. Tanaka, N. Takata, T. Miyazaki, and H. Yamada for discussing the experiment; A. [gmo, S. Inoue, and A. Hirano for valuable discussions and comments on the manuscript. This study was supported by JSPS KAKENHI (21J22555 to A.M.), JSPS KAKENHI Grant-in-Aid for Scientific Research on Innovative Areas, “Willdynamics” (19H05006 to K.S.); AMED under Grant JP21zf0127005 (to Q.L.); the WPI program from Japan’s MEXT.

## Author Contributions

A.M., K.S., T.S., and Q.L. conceived the project and designed the experiments. A.M. performed all stereotaxic surgeries and conducted experiments except as noted. A.M. wrote code for photometry and optogenetics data acquisition and analyses. Y.T. performed tissue samplings. A.M. and Y.T. performed histology analyses and behavioral analyses. K.S. performed retrograde tracing. T.K., A.M., and Y.I. performed ex vivo brain slice imaging. N.N., A.M., and J.S. performed computational modeling. Y.C., A.M., and K.S. prepared AAV vectors. A.M. and T.K. performed the statistical analyses. Y.L. provided the GRAB sensors rDA2.0m and ACh3.0. A.M. and Q.L. wrote original manuscript with input from all co-authors and A.M., T.K., N.N., Y.C., M.Y., H.T., K.S., T.S., and Q.L. edited and reviewed the manuscript.

