## Supplementary figures and tables for "Sequential Transitions of Male Sexual Behaviors Driven by Dual Acetylcholine-Dopamine Dynamics"

### Supplemental Figures

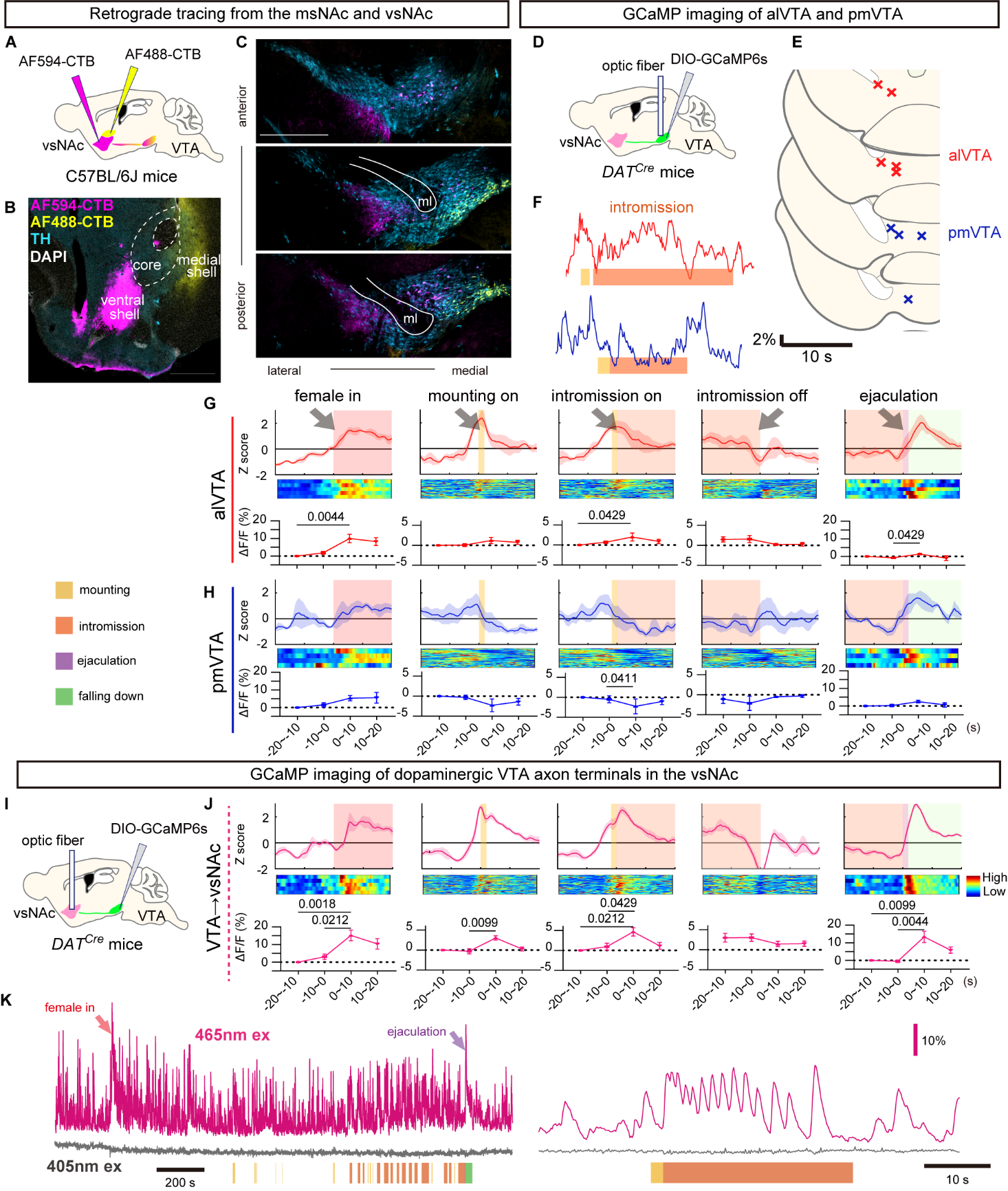

##### Figure S1. Rhythmic Ca^2+^ activity is specifically observed in the DA^VTA-vsNAc^ axon terminals (related to Figure 1).

(A–C) The anterior-lateral VTA projects to the vsNAc, while the posterior-medial VTA projects to the msNAc (scale bars = 500 μm).

(D–H) The neuronal activity of dopaminergic neurons in VTA subregions correspond to the projection sites in NAc but did not fluctuate during intromission. See Table S3.

(D) Strategy for imaging of GCaMP6s around the VTA subregions in *DAT^Cre^* mice.

(E) Recording sites of GCaMP6s imaging in the VTA of *DAT^Cre^* mice: alVTA and pmVTA.

(F) Representative traces for ΔF/F of GCaMP6s around the VTA subregions during intromission with laser illustration of sexual behavior.

(G and H) Same as Figures 1E–1G, but for GCaMP6s fluorescence at the alVTA (G, *n* = 5 mice / group) and pmVTA (H, *n* = 4 mice / group) around each sexual behavior. See Table S3.

(I) Strategy for imaging of GCaMP6s of the axon terminals of the VTA in the vsNAc of *DAT^Cre^* mice (DA^VTA→vsNAc^ axons).

(J) Same as (G and H), but for GCaMP6s signals at DA^VTA→vsNAc^ axons (*n* = 5 mice / group). See Table S4.

(K) Representative traces for ΔF/F of GCaMP6s in DA^VTA→vsNAc^ axons excited by 465 nm (Ca^2+^ dependent) and 405 nm (Ca^2+^ independent) LED around the VTA subregions during intromission with laser illustration of sexual behavior.

Mean ± SEM.

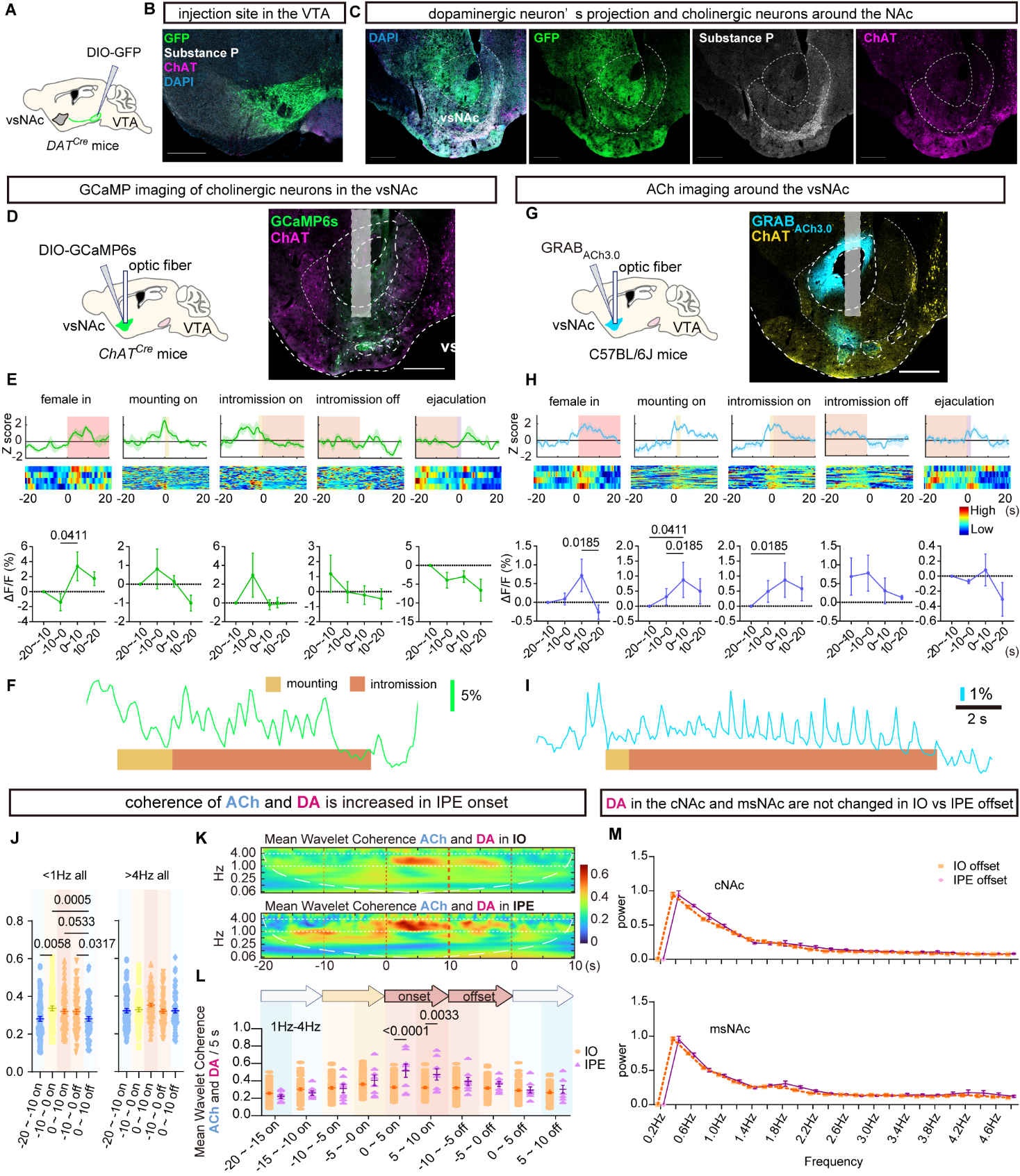

##### Figure S2. Rhythmic ACh release and Ca^2+^ activity of cholinergic interneurons are observed in the vsNAc during intromission (related to Figure 2).

(A–C) Dopaminergic axons project to the vsNAc and cholinergic interneurons in the vsNAc (scale bar = 500 μm). Strategy for labeling of dopaminergic neurons using AAV (A). Immunostained coronal brain sections showing immunostaining of GFP, ChAT, substance P, and DAPI around the VTA (B) and NAc (C).

(D–F) The neuronal activity of cholinergic neurons in vsNAc is rhythmic during intromission.

(G–I) ACh release into vsNAc is rhythmic during intromission.

(D and G) Strategy for imaging of GCaMP6s in ChAT^vsNAc^ neurons in *ChAT^Cre^* mice (D) and GRAB_ACh3.0_ in the vsNAc in C57BL/6J mice (G) (left). Coronal brain sections for GCaMP6s (D) and GRAB_ACh3.0_ (G) immunostained for ChAT (scale bars = 500 μm).

(E and H) Same as Figures 1E–1G, but for GCaMP6s at ChAT^vsNAc^ neurons (E, *n* = 5 mice / group) and GRAB_ACh3.0_ in the vsNAc (H, *n* = 4 mice / group) fluorescence around each sexual behavior. See also Table S5.

(F and I) Representative traces for ΔF/F of GCaMP6s (F) and GRAB_ACh3.0_ (I) during intromission with laser illustration of sexual behavior.

(J) Mean wavelet coherence between GRAB_ACh3.0_ and GRAB_rDA2m_ every 10 s around intromission of <1Hz (left) and >4Hz (right). *n* = 67 events from 3 mice. See Table S7.

(K) Heatmap plot of mean wavelet coherence between GRAB_ACh3.0_ and GRAB_rDA2m_ around IO (top) and IPE (bottom).

(L) Comparison of 1–4 Hz wavelet coherence between GRAB_ACh3.0_ and GRAB_rDA2m_ every 10 s around intromission among IO and IPE. *n* = 59 (IO) and 8 (IPE) events from 3 mice (K and L). See Table S7.

(M) Power spectrum of GRAB_DA2m_ in the cNAc (top, *n* = 10 mice / group) and msNAc (bottom, *n* = 9 mice / group) during 10 s offset of IO and IPE. Two-way ANOVA Šídák's multiple comparisons test over IO offset and IPE offset every 0.2 Hz. *ns* in all frequency in the cNAc and msNAc. See Table S8.

Mean ± SEM.

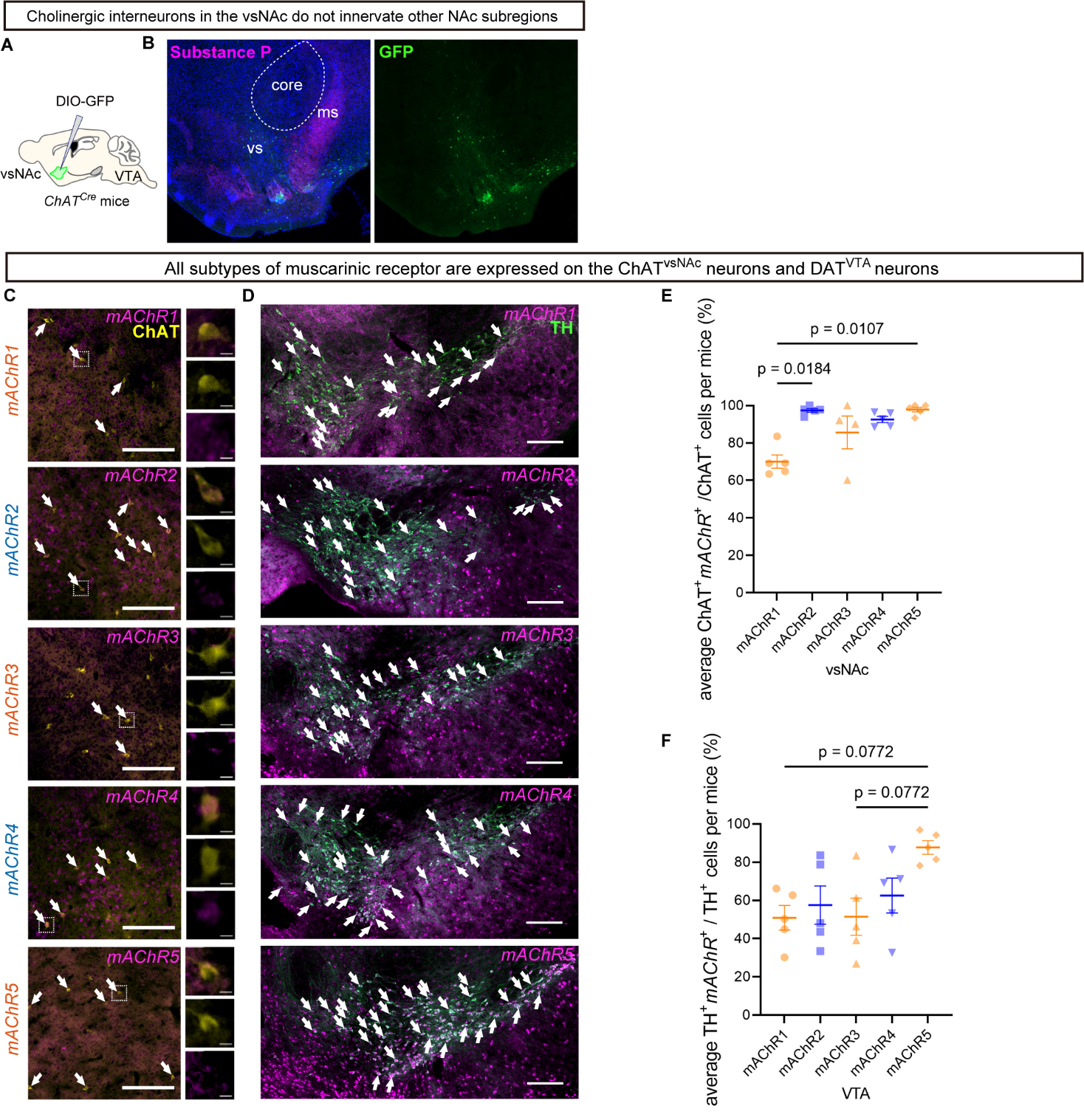

##### Figure S3. In situ hybridization of *mAChR1*–*5* mRNA in the ChAT^vsNAc^ and on DA^VAT^ neurons (related to Figure 3).

(A and B) ChAT^vsNAc^ neurons project to only the vsNAc but not the cNAc and msNAc. Surgery design (A). Coronal brain sections around the NAc stained with Substance P (B).

(C) Coronal brain sections around the NAc stained with anti-ChAT and in situ hybridization of *mAChR* subtype1–5 (*mAChR1, mAChR2, mAChR3, mAChR4,* and *mAChR5* represent *Chrm1*, *Chrm2*, *Chrm3*, *Chrm4*, and *Chrm5*, respectively). Scale bars = 200 µm (left). Enlarged images of white squares surrounding areas in left images (scale bars = 10 µm) (right).

(D) Coronal brain sections around the VTA stained with anti-TH and in situ hybridization of *mAChR* subtypes 1–5 (scale bars = 200 µm). White arrows indicate the double positive cells such as *mAChR1*^+^ChAT^+^ cells (C and D).

(E) The ratio of ChAT^+^ cells expressing each *mAChR* subtype to the total number of ChAT^+^ cells in the vsNAc (%). *n* = 5 mice in each *mAChR1,2,4,5* and 4 mice in *mAChR3*

(F) The ratio of TH^+^ cells expressing each *mAChR* subtype to the total number of TH^+^ cells in the VTA (%). *n* = 5 mice in each *mAChR*1–5.

Mean ± SEM. See also Table S9 (E and F).

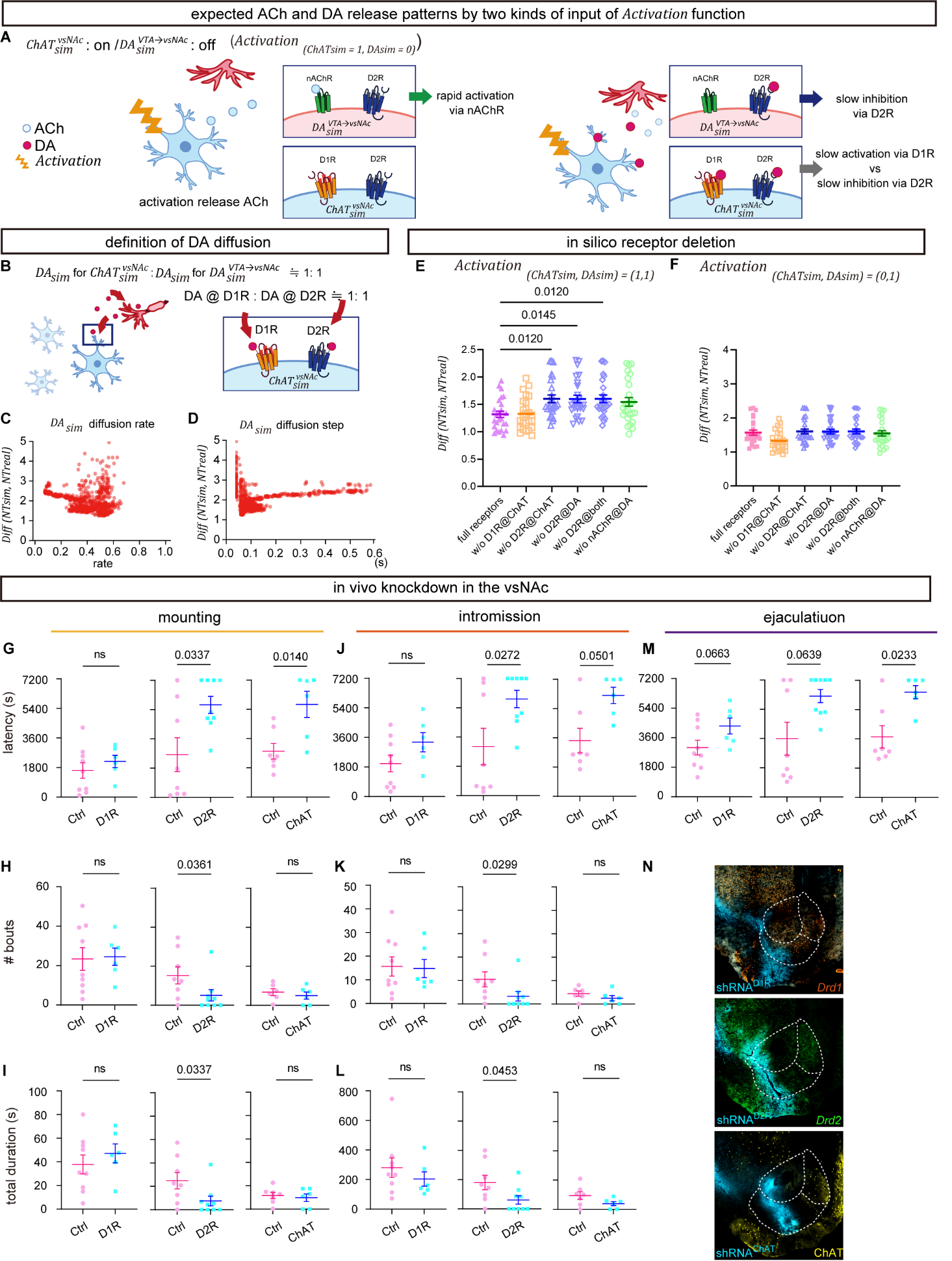

##### Figure S4. Modeling of dual ACh-DA rhythms and the effects of knockdown of D1R, D2R, or ChAT expression in the vsNAc on male sexual behaviors (related to Figure 4).

(A) The expected release pattern of ACh and DA depend on the different activation inputs on $ChAT_{sim}^{vsNAc}$ ($Activation_{\left( 1,0 \right)})$.

(B) Distribution setting of released $DA_{sim}$ toward D1R and D2R on $ChAT_{sim}^{vsNAc}$ neurons and D2R on $DA_{sim}^{VTA\to vsNAc}$ axon terminals (D1R on $ChAT_{sim}^{vsNAc}$: D2R on $ChAT_{sim}^{vsNAc}$: D2R on $DA_{sim}^{VTA\to vsNAc}$=1:1:2).

(C) Relationship between $Diff\left( NT_{sim}, NT_{real} \right)$ and explored DA diffusion rate.

(D) Relationship between $Diff\left( NT_{sim}, NT_{real} \right)$ and the explored DA diffusion step.

(E and F) Comparison of $Diff\left( NT_{sim}, NT_{real} \right)$ with each receptor deleted condition under activation both $ChAT_{sim}^{vsNAc}\mathrm{and} DA_{sim}^{VTA\to vsNAc}$(E) and $DA_{sim}^{VTA\to vsNAc}$ only (F)$.$ See Table S12.

(G–M) The results of gene knockdown of D1R (left), D2R (center) and ChAT (right) using AAV-shRNA^D1R^, AAV-shRNA^D2R^, and AAV-shRNA^ChAT^. See also Table S15.

(G–I) The knockdown effects on mounting’s latency (G), the number of its bout (H), and its total duration (I).

(J–L) The knockdown effects on intromission’s latency (J), the number of its bout (H), and its total duration (I).

(M) The knockdown effects on the ejaculation’s latency.

(N) The representative brain slices expressing AAV-shRNA^D1R^ (top), AAV-shRNA^D2R^ (middle), and AAV-shRNA^ChAT^ (bottom) in the vsNAc.

Mean ± SEM.

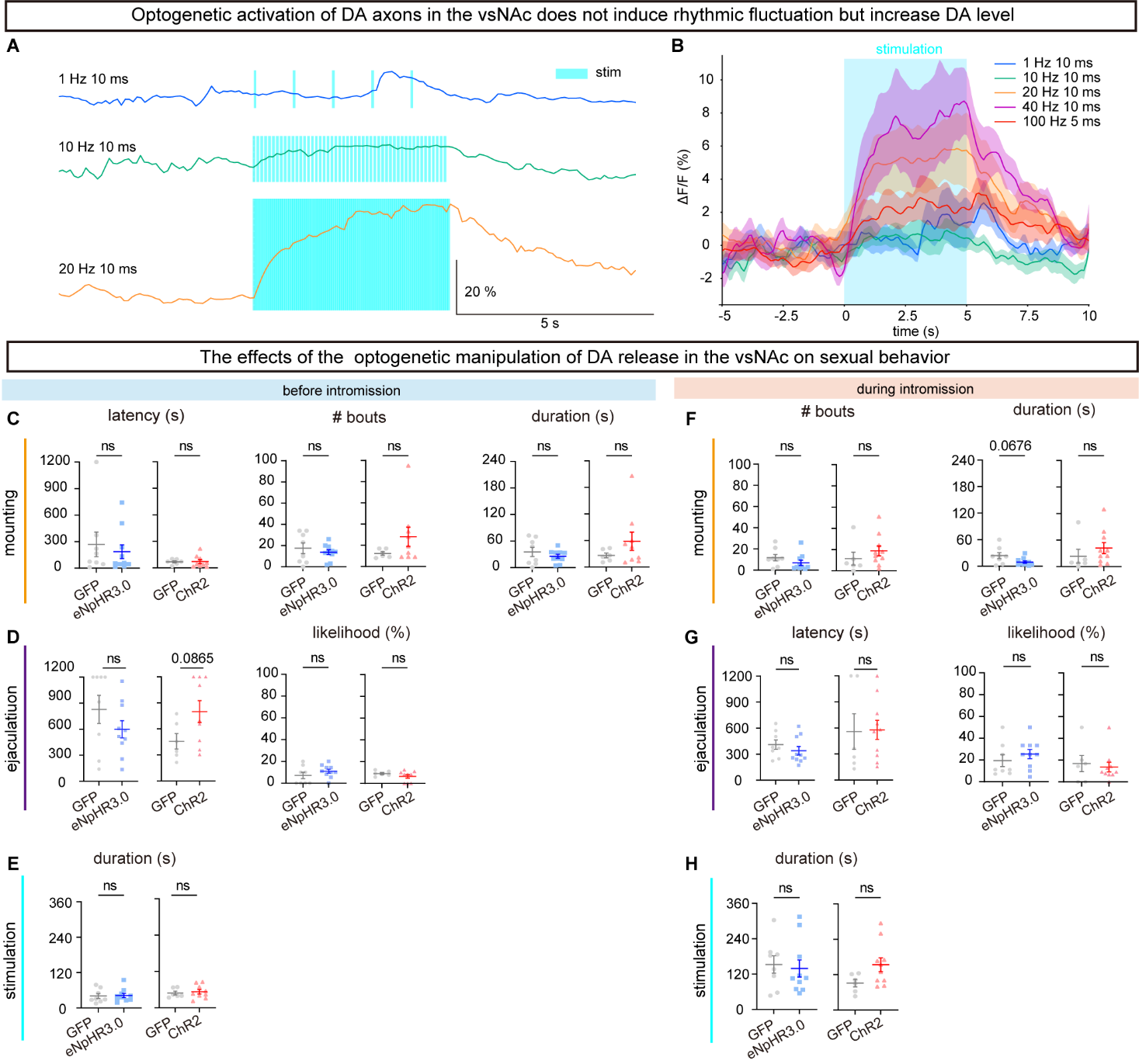

##### Figure S5. The effects of optogenetic manipulation of DA release in the vsNAc around intromission on male sexual behaviors (related to Figure 5).

(A) Representative traces of GRAB_DA2m_ around optogenetic stimulation of DA^VTA→vsNAc^ axons with 1 Hz 10 ms (top), 10 Hz 10 ms (middle), and 20 Hz 10 ms (bottom).

(B) PETP of ΔF/F of GRAB_DA2m_ fluorescence around the optogenetic stimulation of DA^VTA→vsNAc^ axons.

(C–J) The effects of optogenetic inhibitory (left, GFP^DA^ vs. eNpHR3.0^DA^) and excitatory (right, GFP^DA^ vs. ChR2^DA^) manipulation of DA^VTA→vsNAc^ axons before intromission (C–E) and during intromission (F–H).

(C–E) The effects of optogenetic manipulation of DA^VTA→vsNAc^ axons before intromission on the mounting (C), ejaculation (D), and stimulation duration (E).

(F–H) The effects of optogenetic manipulation of DA^VTA→vsNAc^ axons during intromission on the mounting (F), ejaculation (G), and stimulation duration (H).

Latency (s) = the 1^st^ target behavior onset – the 1^st^ optogenetic stimulation onset.

Ejaculation likelihood (%) = ejaculation occurred or not in the test (1 or 0) / # optogenetic stimulation.

Mean ± SEM. See also Table S18 (C–H).

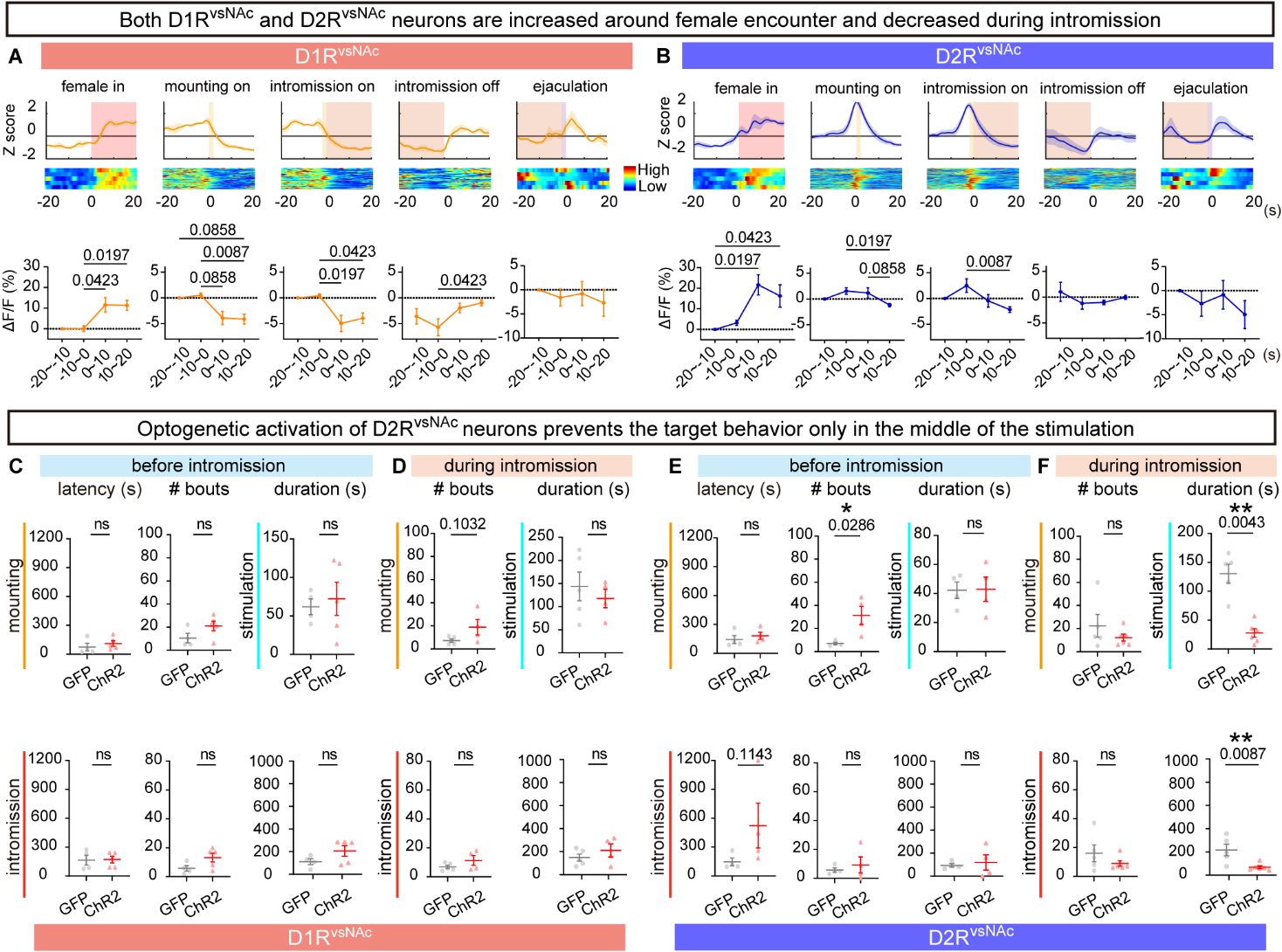

##### Figure S6. Differential effects of optogenetic activation of D1R^vsNAc^ and D2R^vsNAc^ neurons around intromission on male sexual behaviors (related to Figure 6).

(A and B) Same as Figures 1E–1G, but for GCaMP fluorescence at D1R^vsNAc^ neurons (A, *n* = 5 mice / group) and D2R^vsNAc^ neurons (B, *n* = 5 mice / group) around each sexual behavior. See Table S19 (A) and S20 (B).

(C–F) The effects of optogenetic excitatory (ChR2) manipulation of D1R^vsNAc^ (C and D, GFP^D1R^ vs. ChR2^D1R^) and D2R^vsNAc^ (E and F, GFP^D2R^ vs. ChR2^D2R^) before intromission (C and E) and during intromission (D and F). See Table S21 (C and D) and S22 (E and F).

Latency (s) = the 1^st^ target behavior onset – the 1^st^ optogenetic stimulation onset.

Mean ± SEM. ^*^*p* < 0.05, ^**^*p* < 0.01.

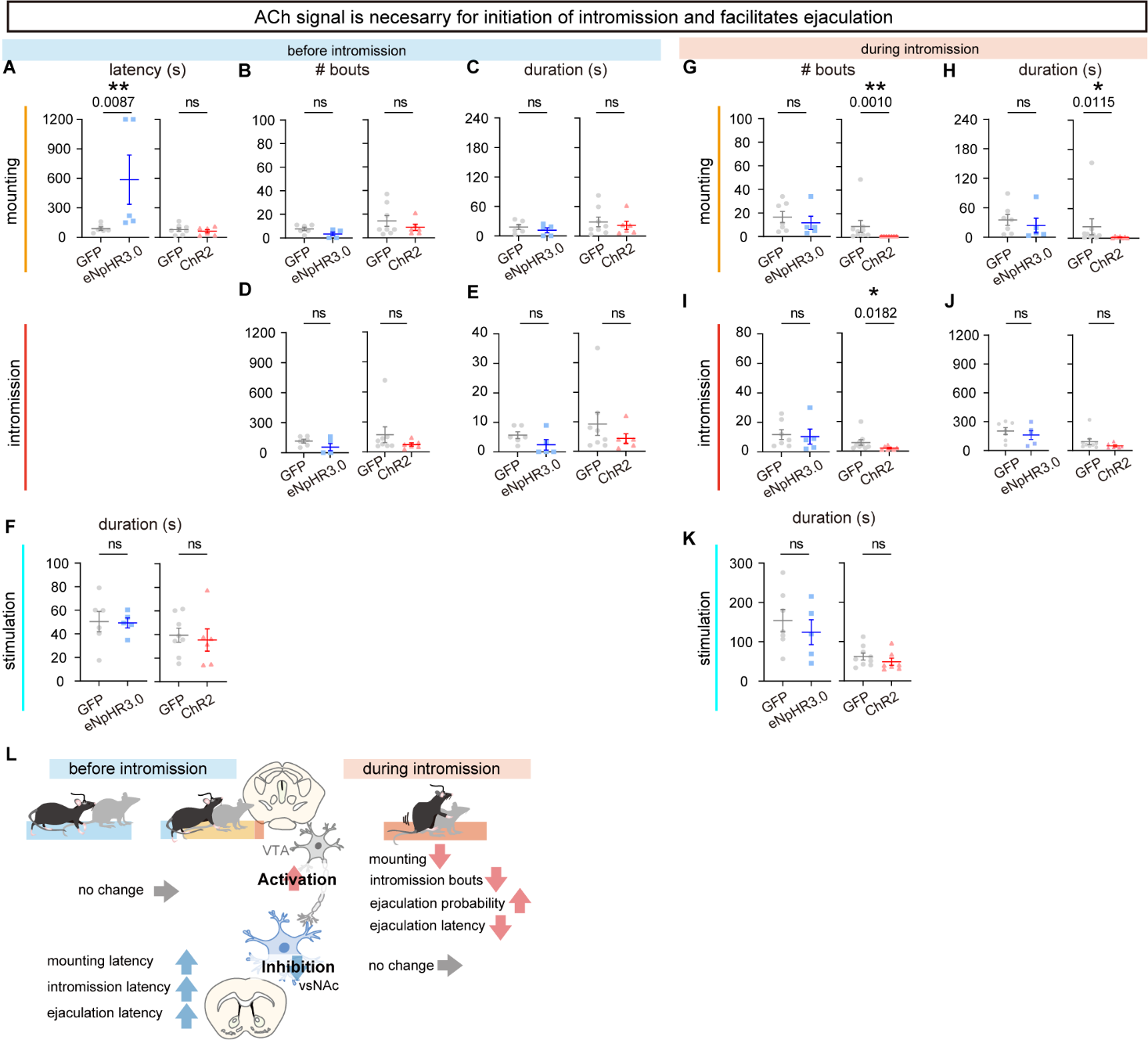

##### Figure S7. The effects of optogenetic activation of ACh release in the vsNAc around intromission on male sexual behaviors (related to Figure 7).

(A–K) The effects of optogenetic inhibitory (left, GFP^ACh^ vs. eNpHR3.0^ACh^) and excitatory (right, GFP^ACh^ vs. ChR2^ACh^) manipulation of ChAT^vsNAc^ neurons before intromission (A**–**F) and during intromission (G**–**K).

(A**–**F) Inhibition of ChAT^vsNAc^ neurons before intromission prolongs the latency of mounting behavior (A), while inhibition does not affect the sexual behavior (A**–**G). The duration of optogenetic stimulation is not different depending on each group (F). See Table S23.

Latency (s) = the 1^st^ target behavior onset – the 1^st^ optogenetic stimulation onset.

(G**–**K) Activation of ChAT^vsNAc^ neurons during intromission decreases the number of mounting bouts (G), intromission bouts (I) and the total duration of mounting (H), while inhibition does not affect sexual behavior (G**–**J). The duration of optogenetic stimulation is not different depending on each group (K). See Table S24.

(L) Optogenetic activation of ChAT^vsNAc^ neurons before intromission does not affect sexual behavior, while inhibition increases the latency of mounting, intromission, and ejaculation. Optogenetic activation of ChAT^vsNAc^ neurons during intromission decreases mounting, the number of intromission bouts, and ejaculation latency, and increases ejaculation probability which the optogenetic manipulation induces ejaculation.

Mean ± SEM. ^*^*p* < 0.05, ^**^*p* < 0.01.

### Supplemental Tables

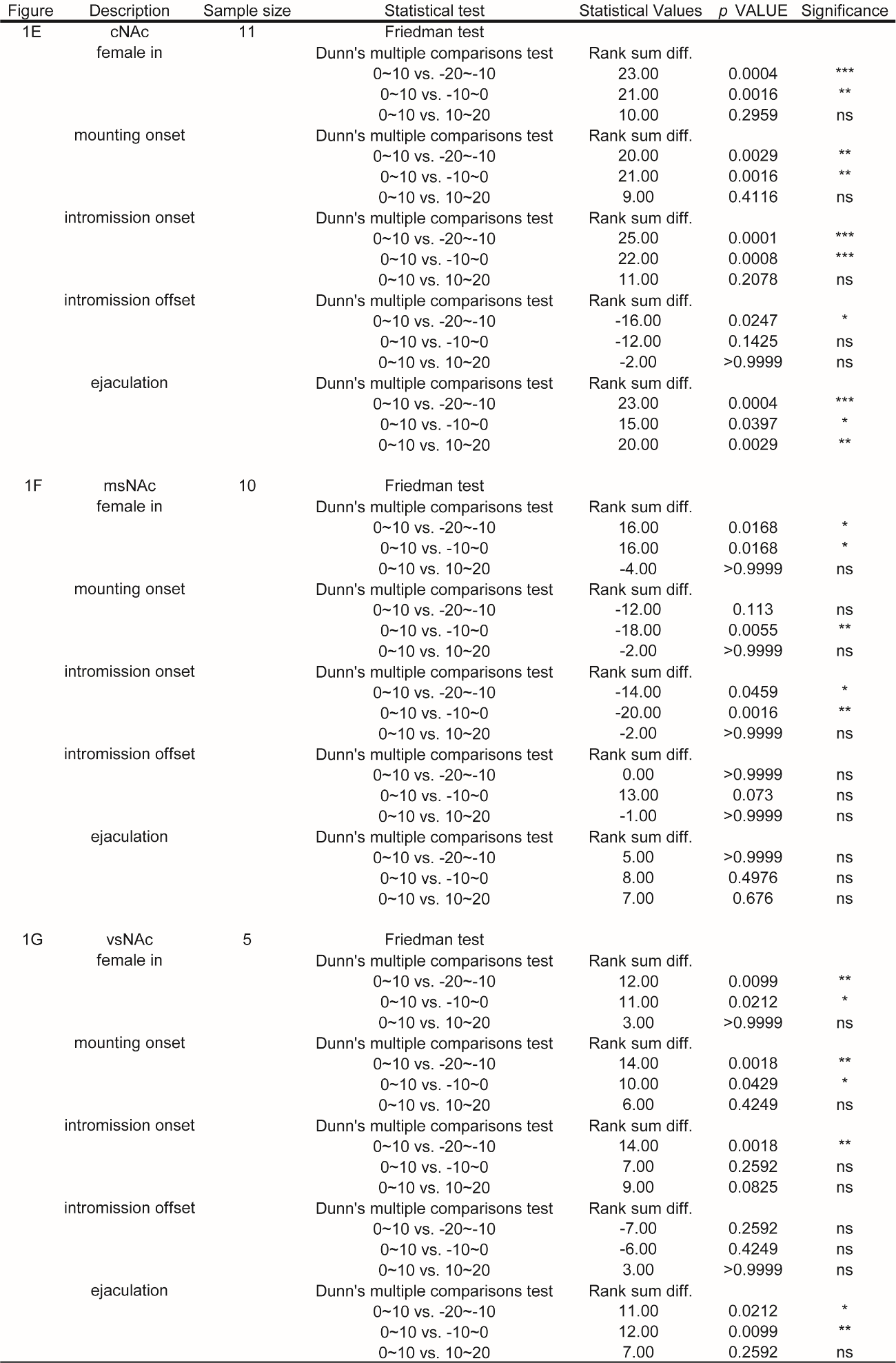

##### Table S1. Comparison of GRAB_DA2m_ intensity around each sexual behavior (related to Figures 1E–1G).

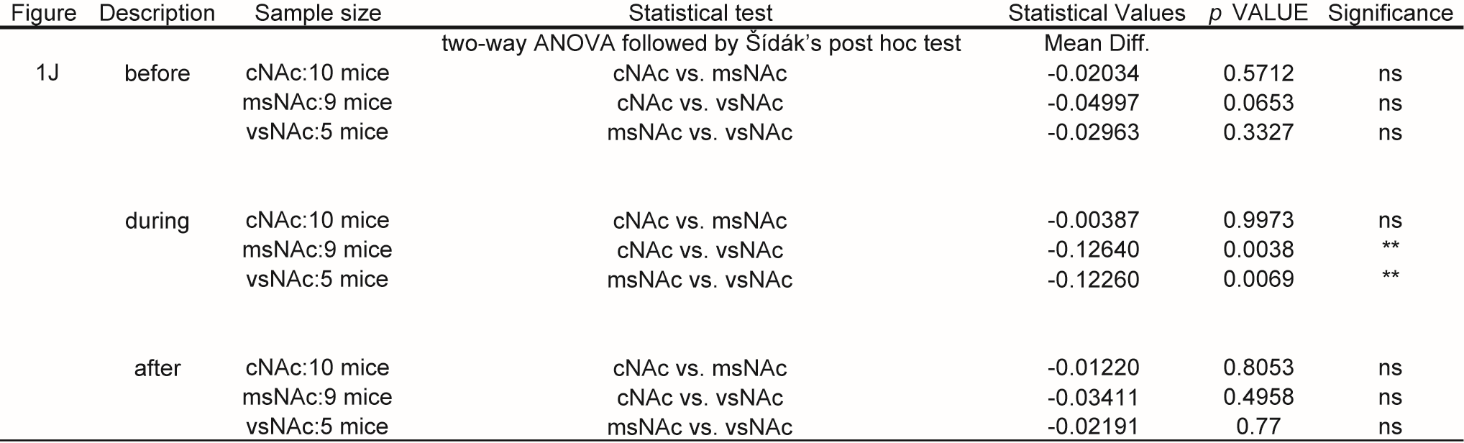

##### Table S2. Comparison of the mean 1–2Hz power spectrum in GRAB_DA2m_ signals around intromission (related to Figure 1J).

##### Table S3. Comparison of GCaMP dynamics of the somata of alVTA and pmVTA neurons during male sexual behaviors (related to Figures S1G and S1H).

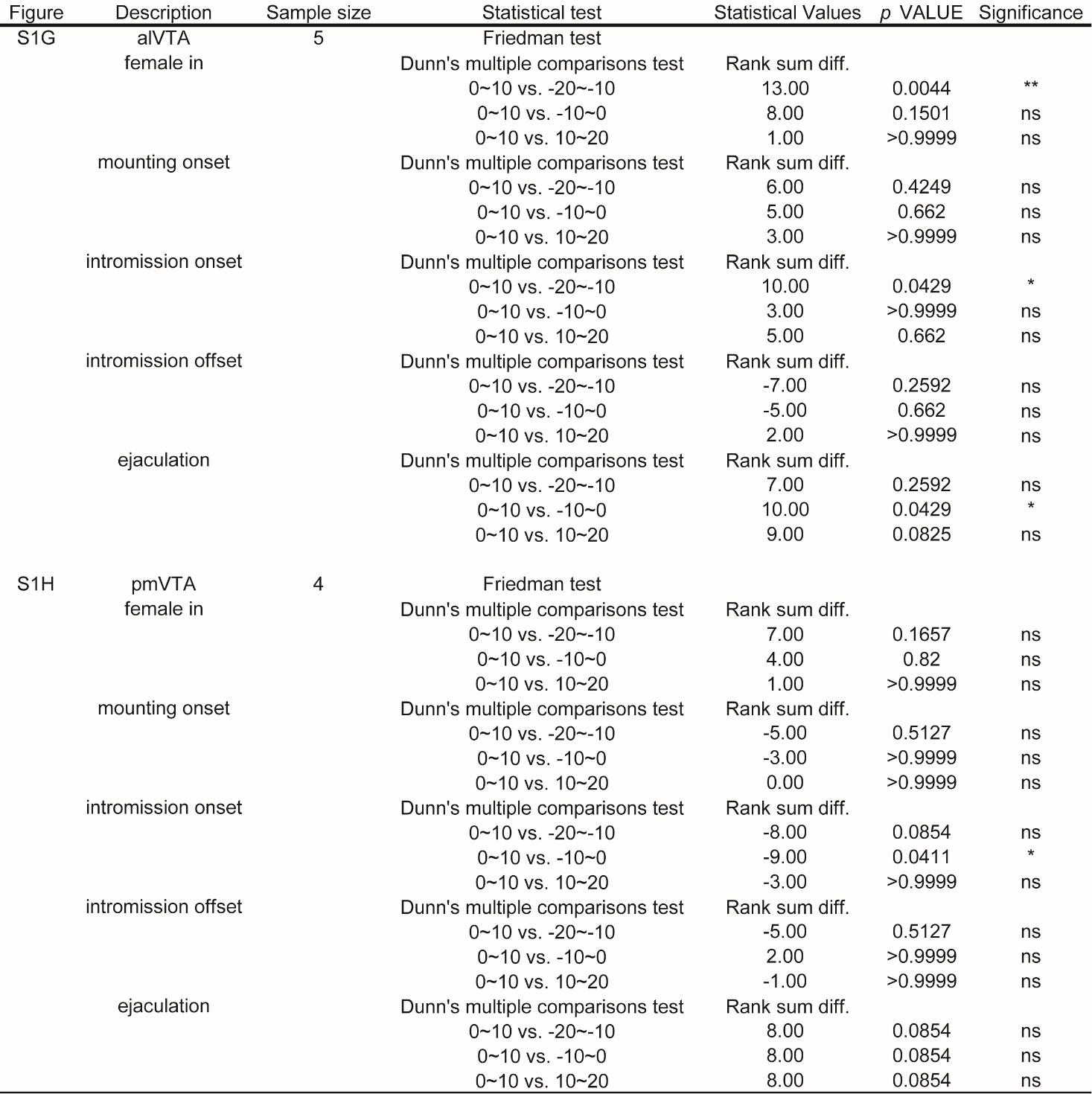

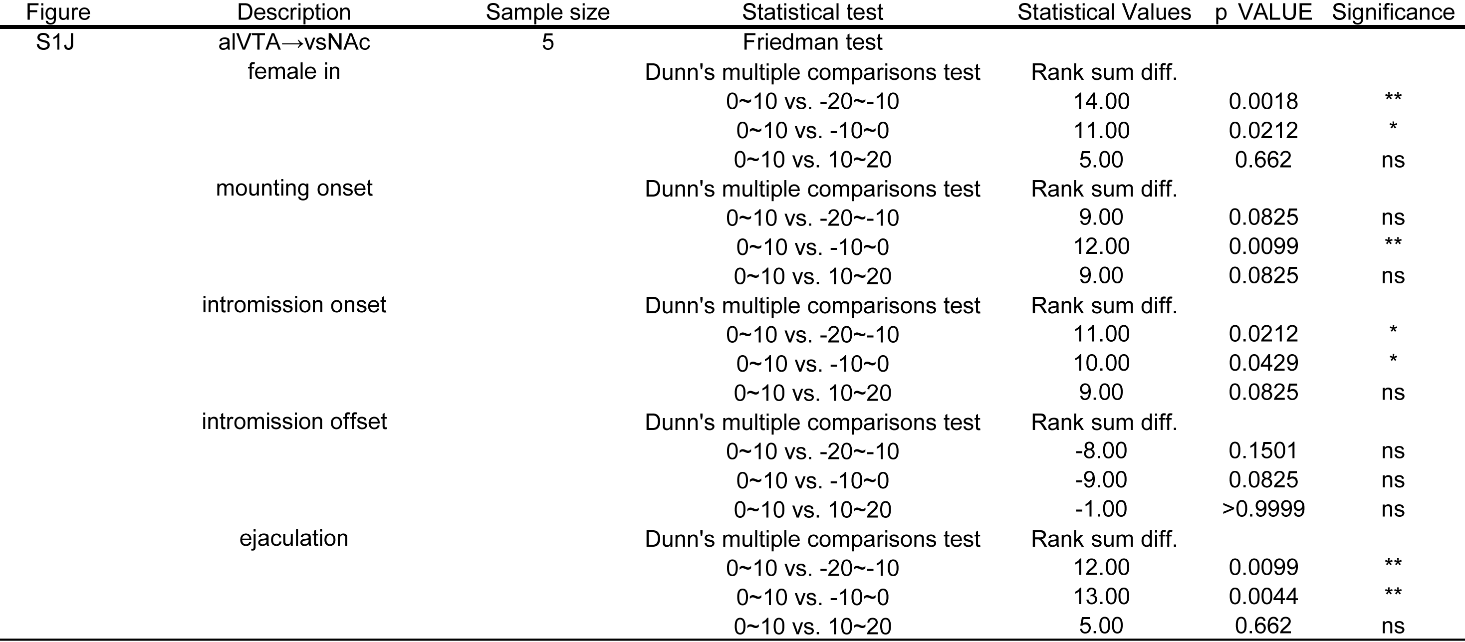

##### Table S4. Comparison of GCaMP dynamics of the DA^VTA^ axons in the vsNAc during male sexual behaviors (related to Figure S1J).

##### Table S5. Comparison of GCaMP dynamics in ChAT^vsNAc^ neurons and GRAB_ACh_ dynamics during male sexual behaviors (related to Figures S2E and S2H).

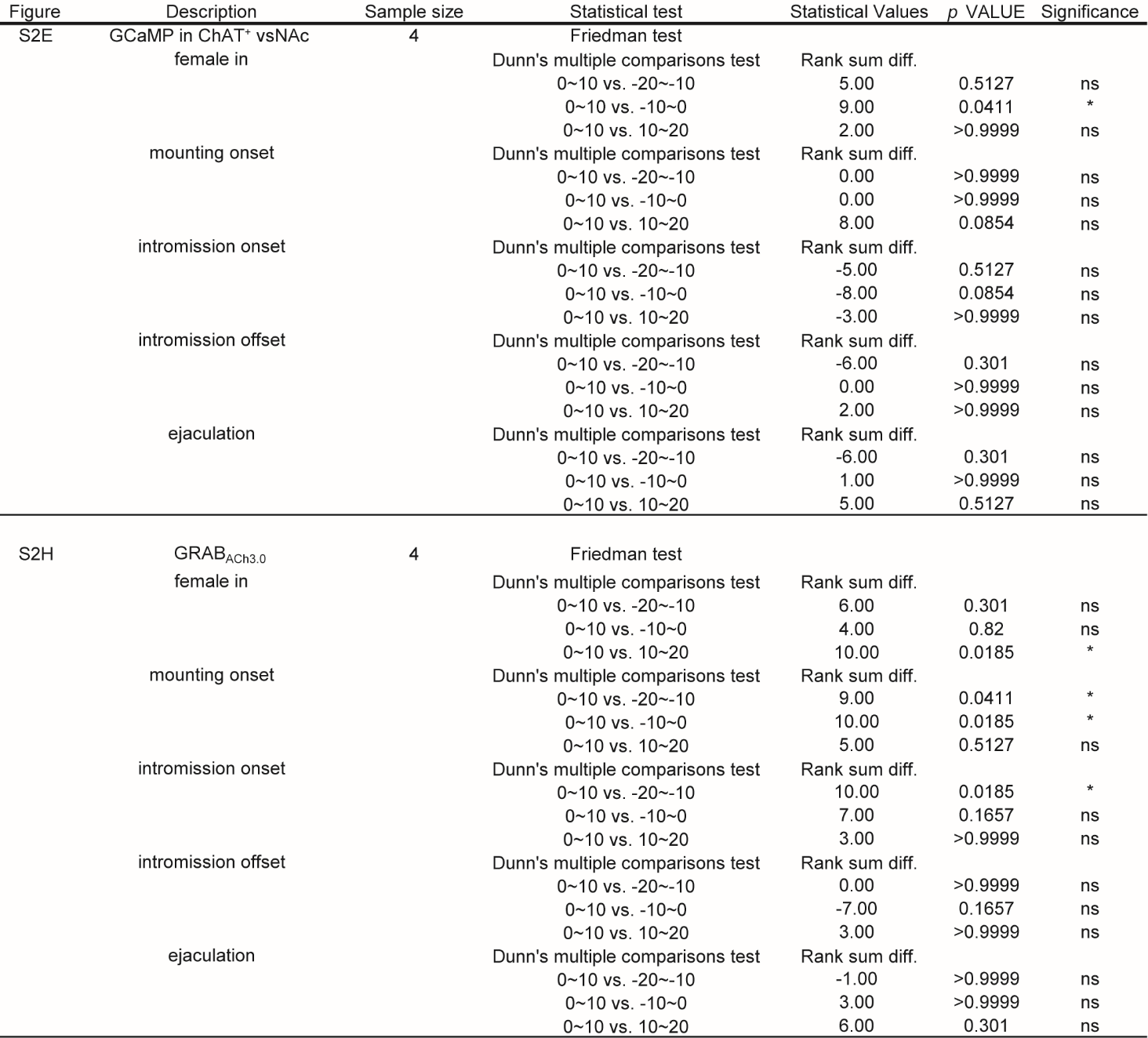

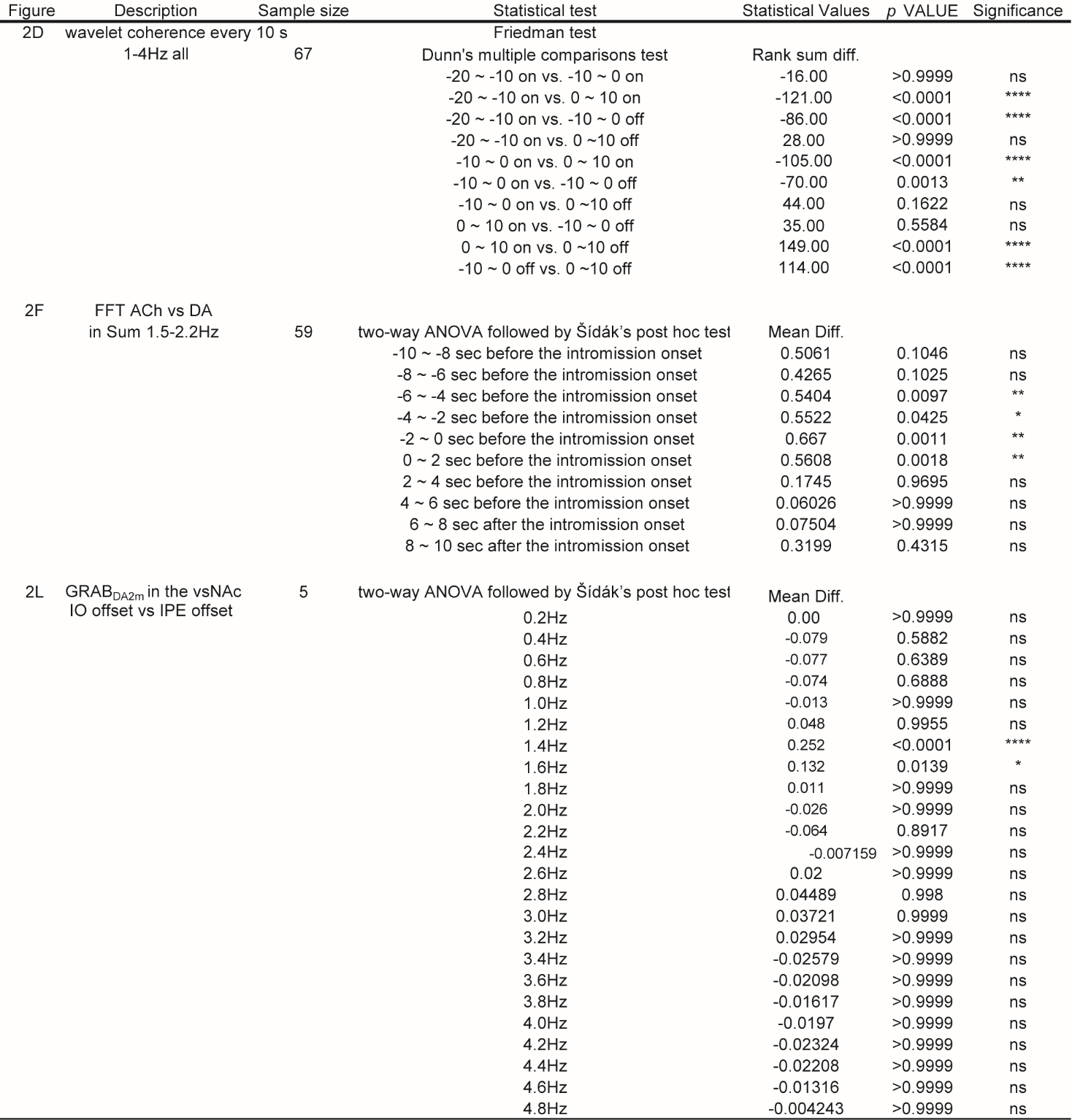

##### Table S6. Analysis of power spectrum and coherence between ACh and DA rhythms in the vsNAc around intromission (related to Figures 2D, 2F, and 2L).

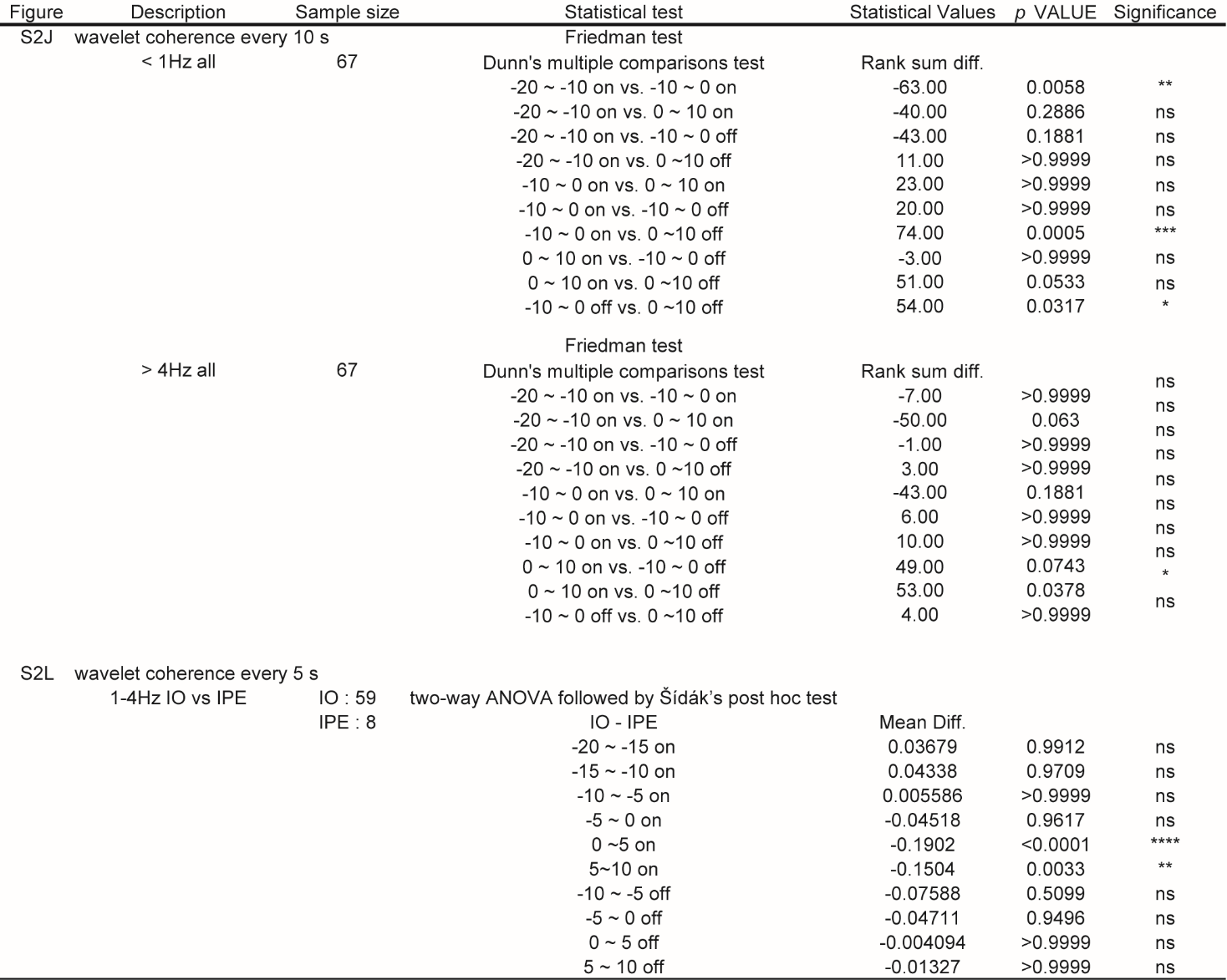

##### Table S7. Comparison of coherence between ACh and DA rhythms in the vsNAc around intromission (related to Figures S2J and S2L).

##### Table S8. Power spectrum analysis of GRAB_DA2m_ dynamics in the cNAc and msNAc during intromission (related to Figure S2M).

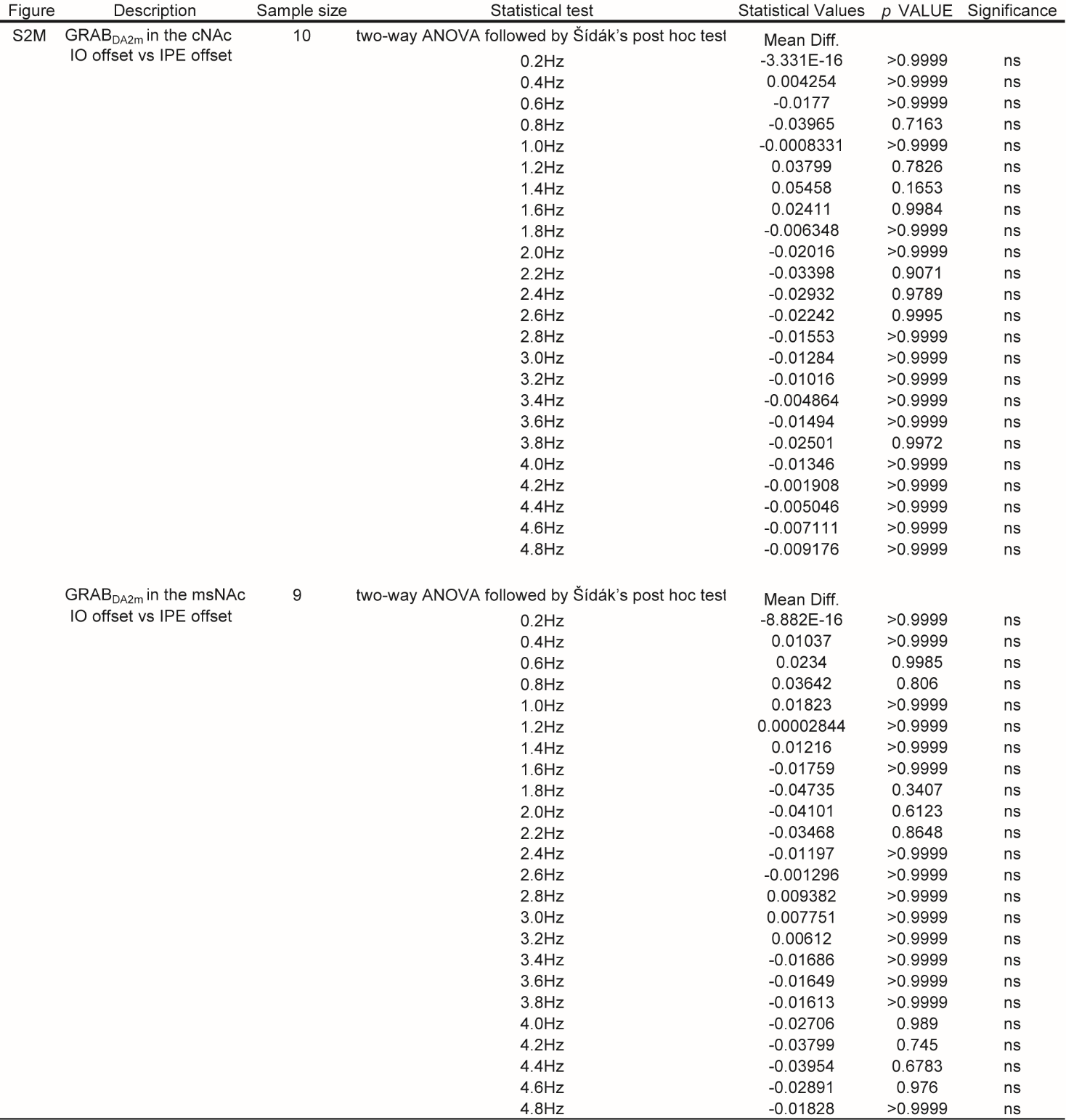

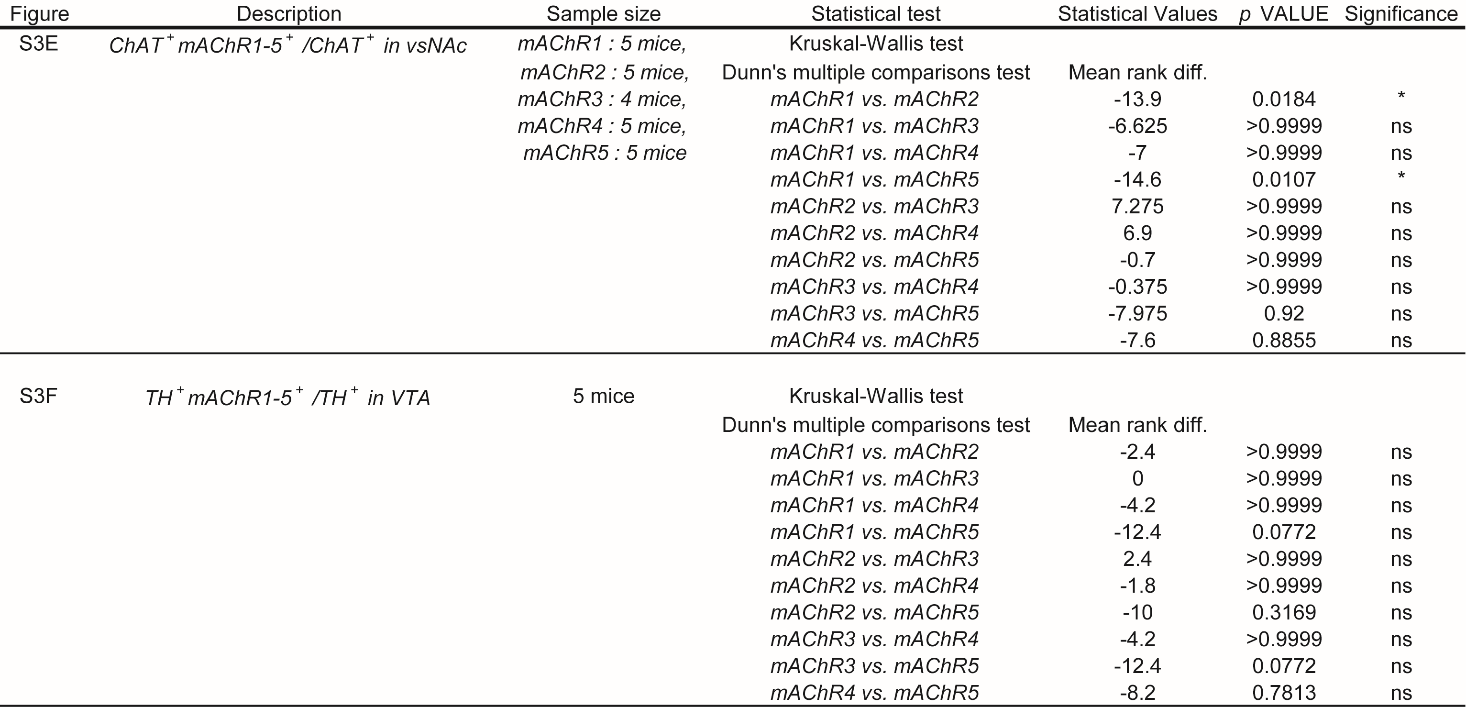

##### Table S9. Quantification of *mAChR1-5*^+^ cholinergic neurons in the vsNAc and *mAChR1-5^+^* dopaminergic neurons in the VTA (related to Figures S3E and S3F)

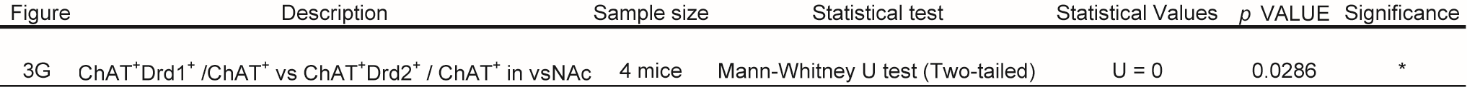

##### Table S10. Quantification of the *Drd1*^+^ or *Drd2*^+^cholinergic neurons in the vsNAc (related to Figure 3G)

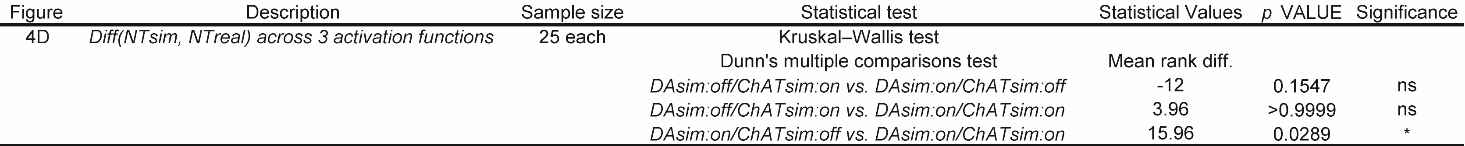

##### Table S11. Comparison of $\boldsymbol{Diff}\left( \boldsymbol{N}\boldsymbol{T}_{\boldsymbol{sim}}\boldsymbol{, N}\boldsymbol{T}_{\boldsymbol{real}} \right)$ in the three different activation input patterns with four full receptors (related to Figure 4D)

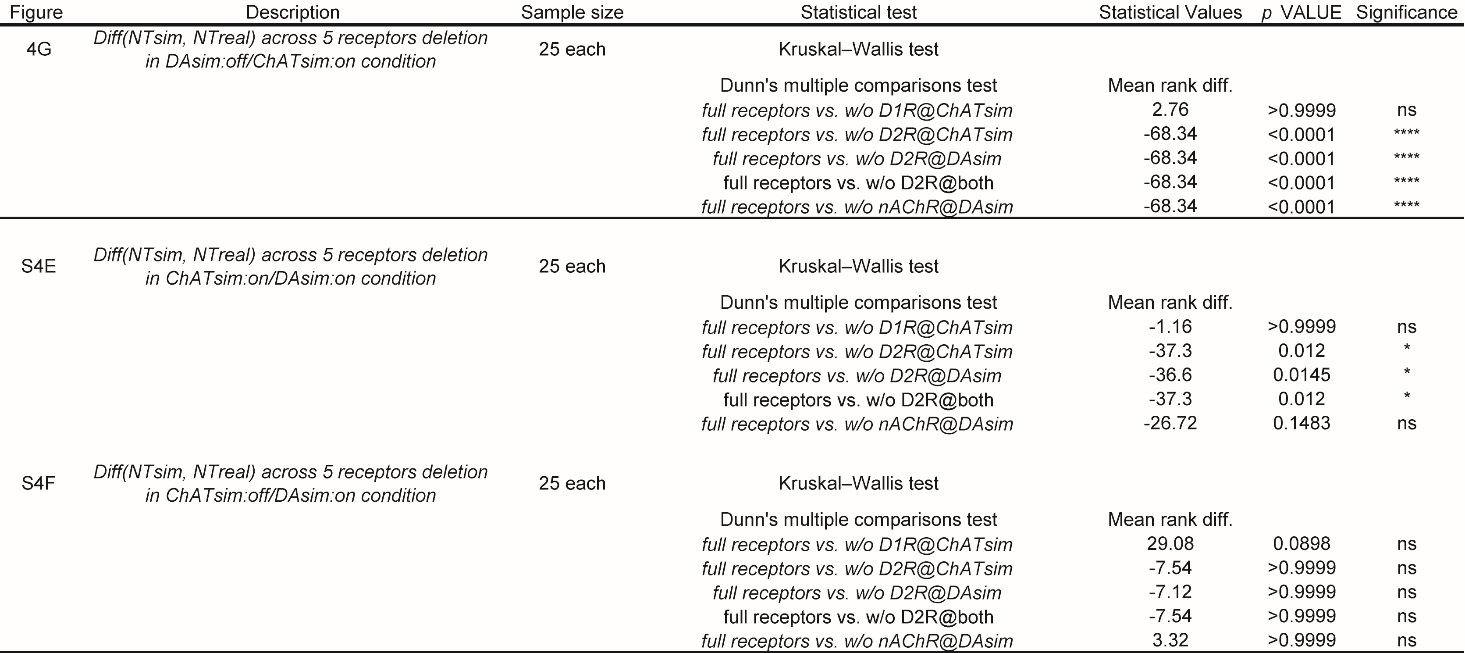

##### Table S12. Comparison of $\boldsymbol{Diff}\left( \boldsymbol{N}\boldsymbol{T}_{\boldsymbol{sim}}\boldsymbol{, N}\boldsymbol{T}_{\boldsymbol{real}} \right)$ with deletion of each receptor in the three different activation input patterns (related to Figures 4G, S4E, and S4F)

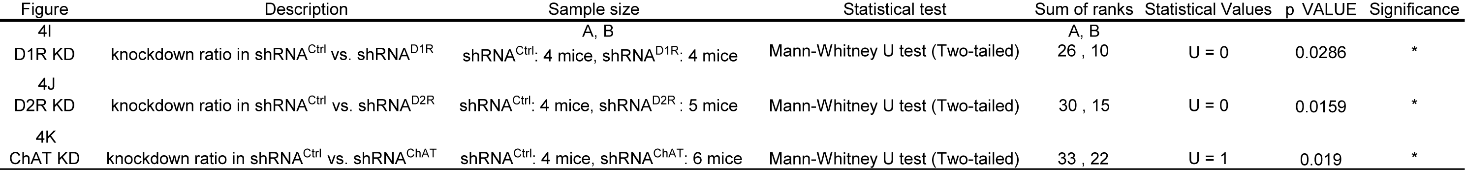

##### Table S13. Comparison of knockdown score between shRNA-Scramble and shRNA-target (related to Figures 4I–4K).

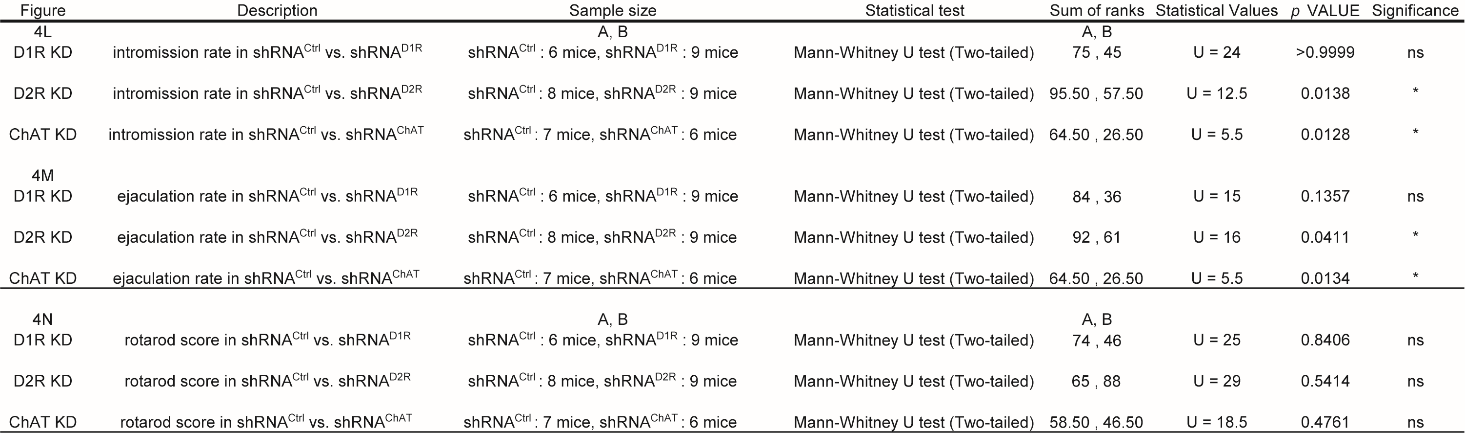

##### Table S14. Comparison of intromission rate, ejaculation rate, and rotarod median score between shRNA-Scramble and shRNA-target expressing mice (related to Figures 4L–4N).

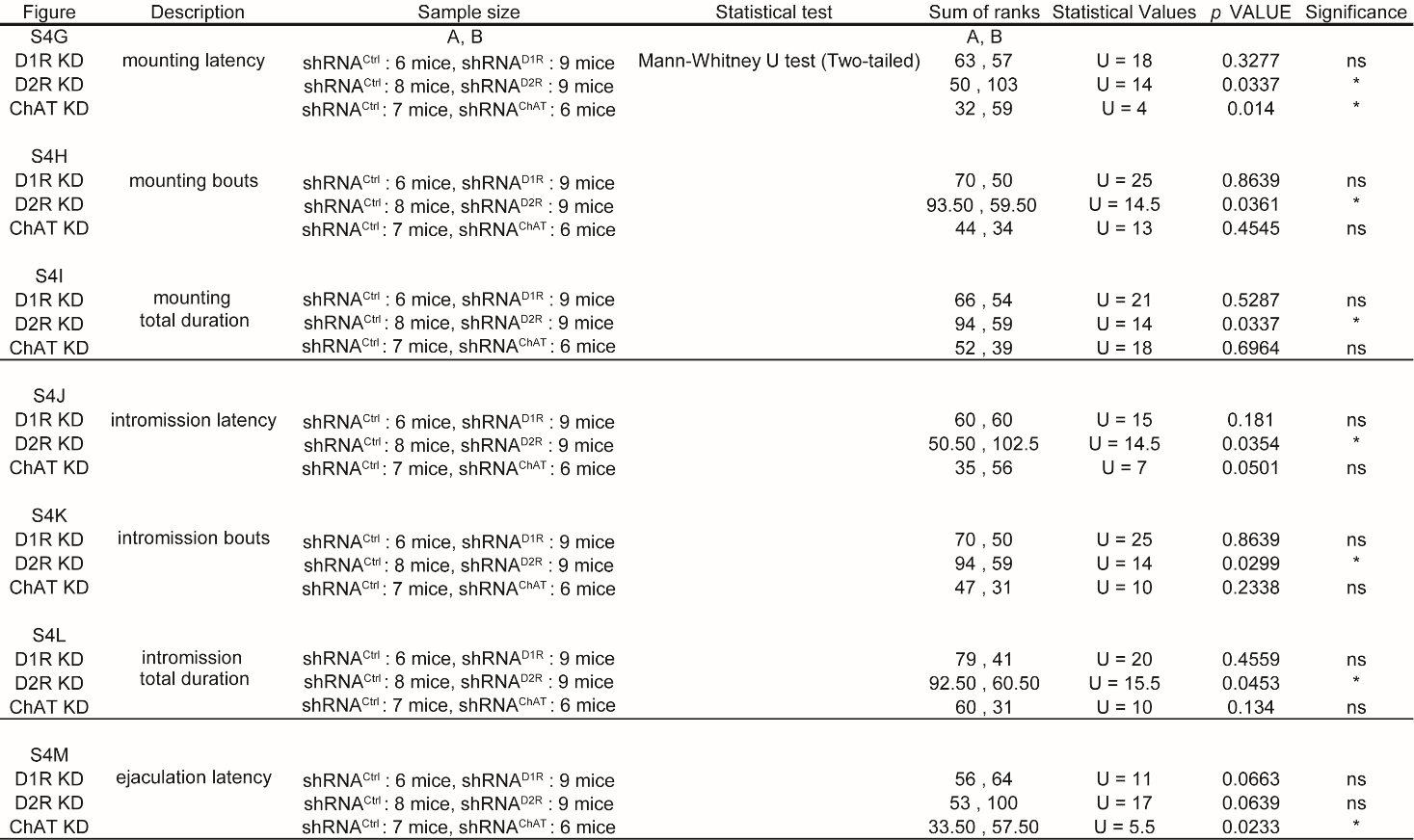

##### Table S15. Quantification of the effects of knockdown of ChAT, D1R, or D2R expression in the vsNAc on male sexual behaviors (related to Figures S4G–S4M)

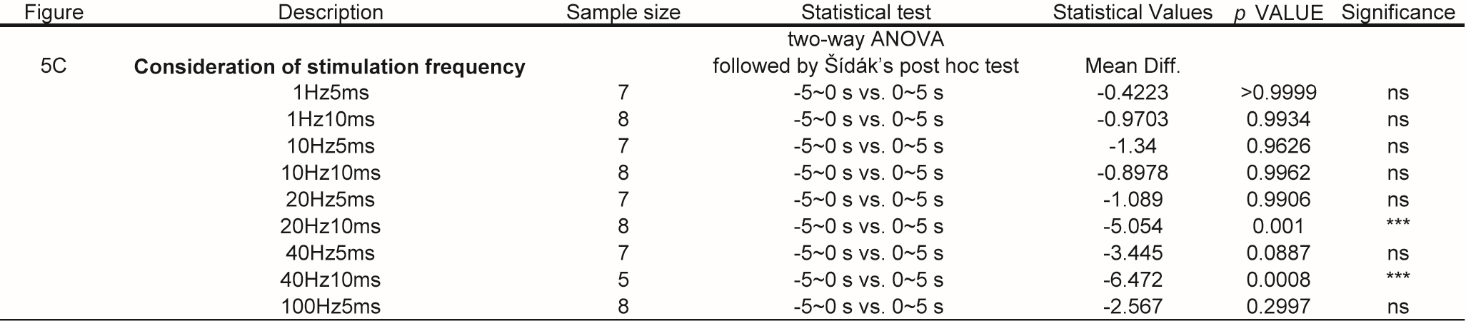

##### Table S16. Analysis of the effects on DA release level following optogenetic manipulation of DA axons in the vsNAc (related to Figure 5C)

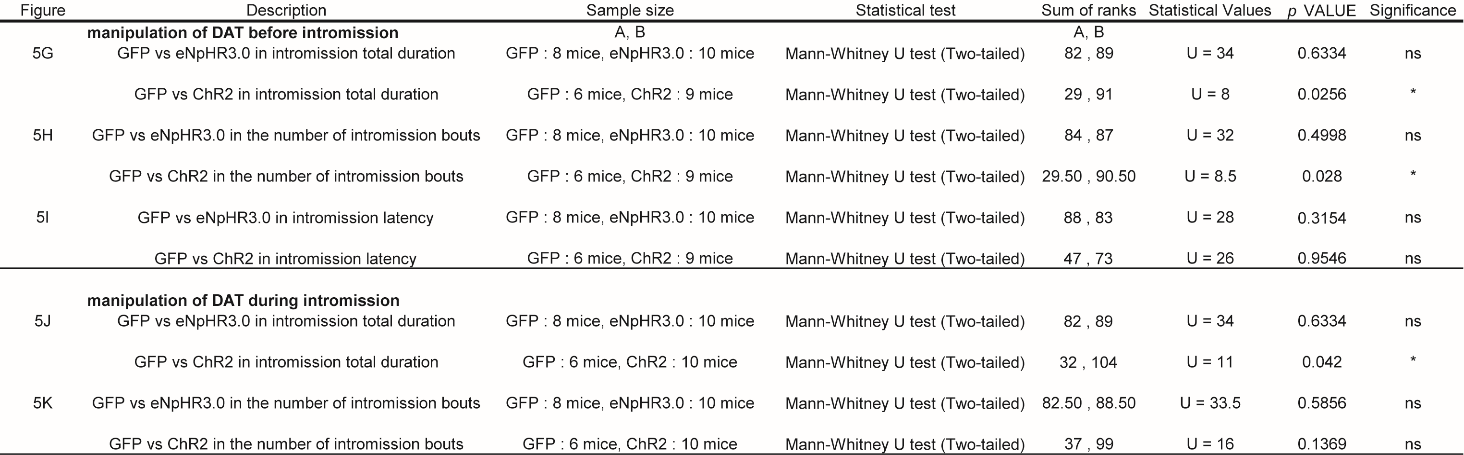

##### Table S17. Analysis of the effects on the intromission total duration and ejaculation latency following optogenetic manipulation of DA release in the vsNAc (related to Figures 5G–5K)

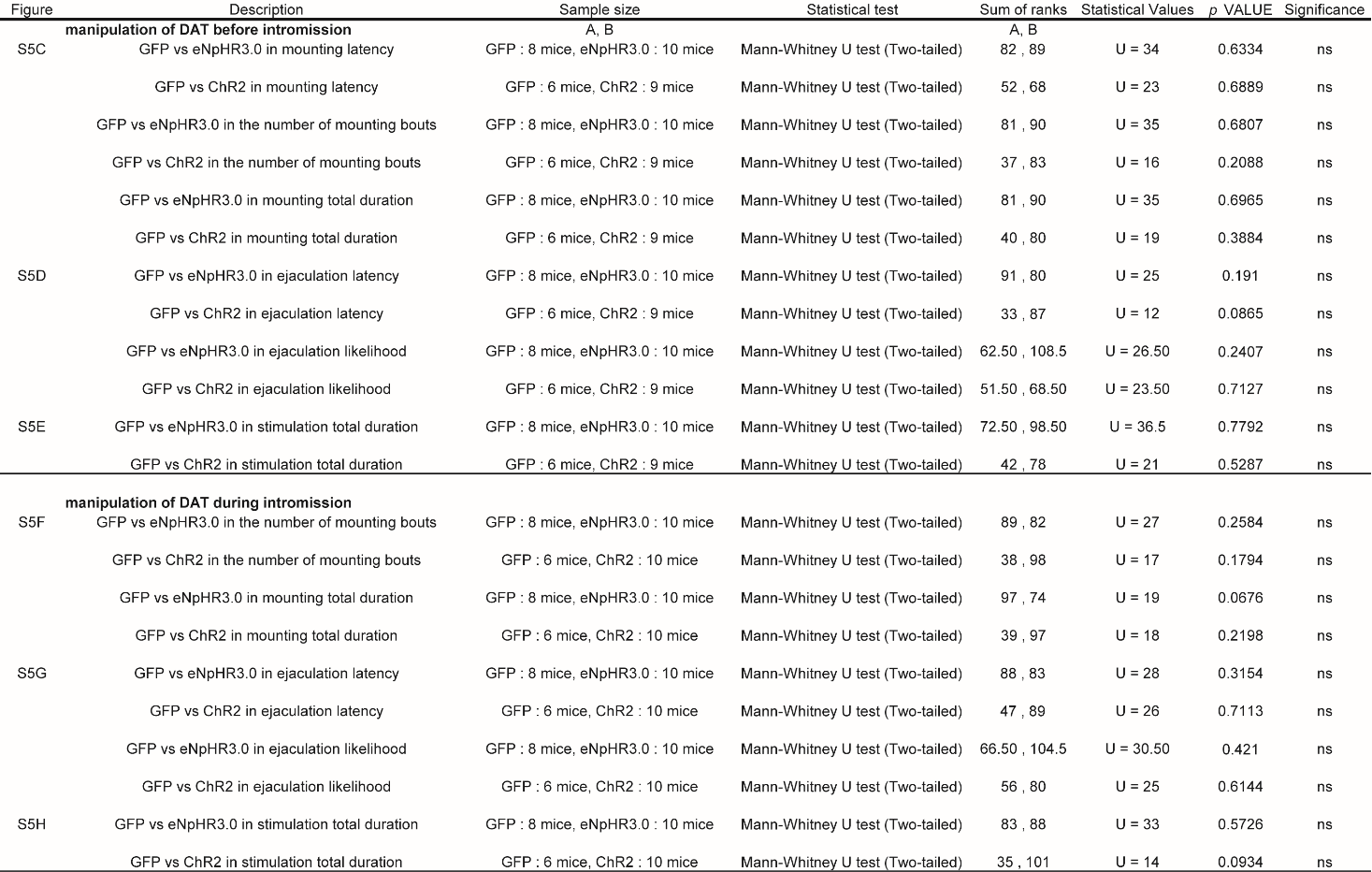

##### Table S18. Analysis of the effects on sexual behaviors following optogenetic manipulation of DA release in the vsNAc (related to Figure S5)

##### Table S19. Analysis of GCaMP dynamics of D1R^vsNAc^ neurons during male sexual behaviors (related to Figure S6A).

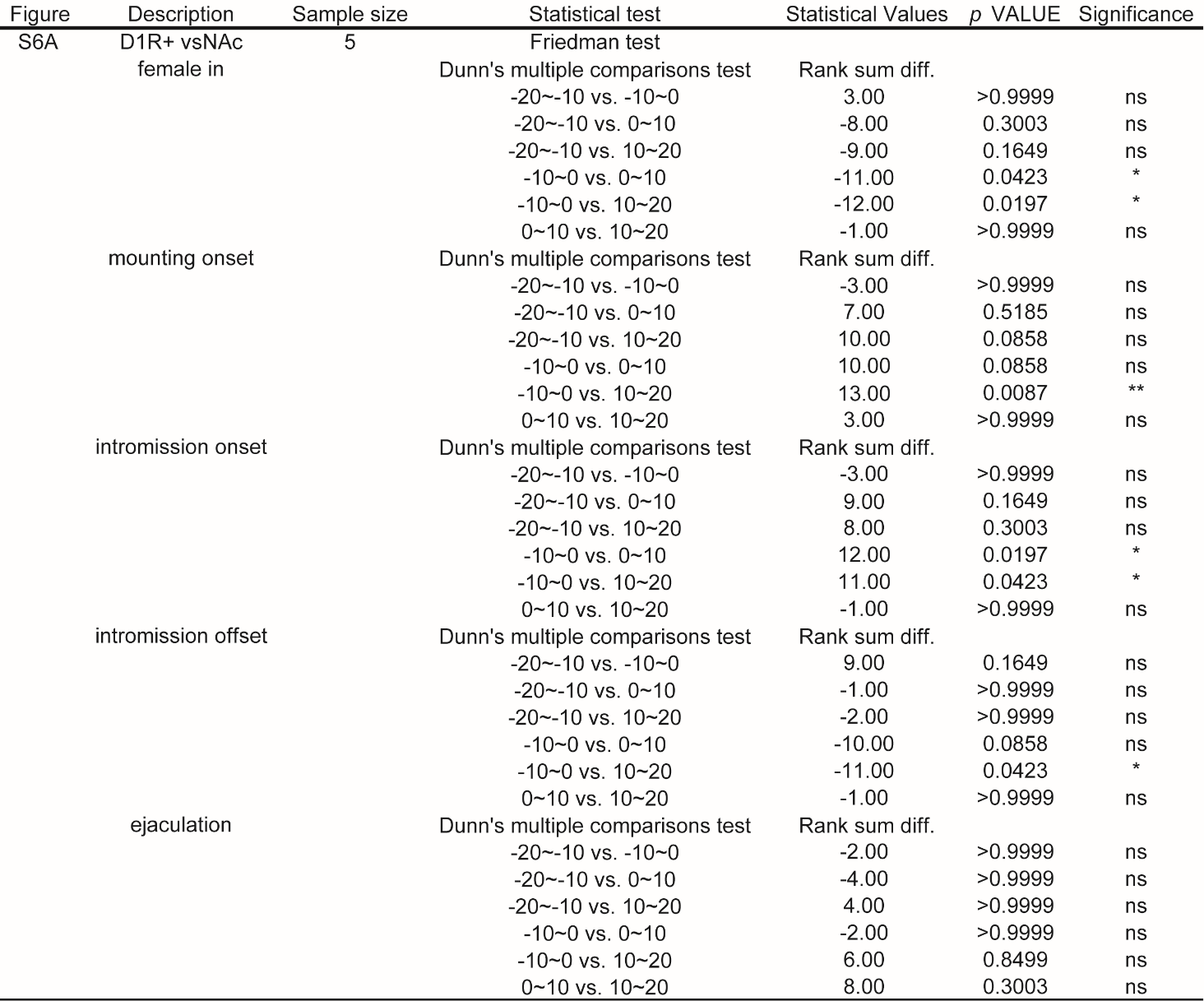

##### Table S20. Analysis of GCaMP dynamics D2R^vsNAc^ neurons during male sexual behaviors (related to Figure S6B). Table S21. Analysis of the effects on sexual behaviors following optogenetic activation of D1R^vsNAc^ neurons before and during intromission (related to Figures 6J–6Q, S6C, and S6D)

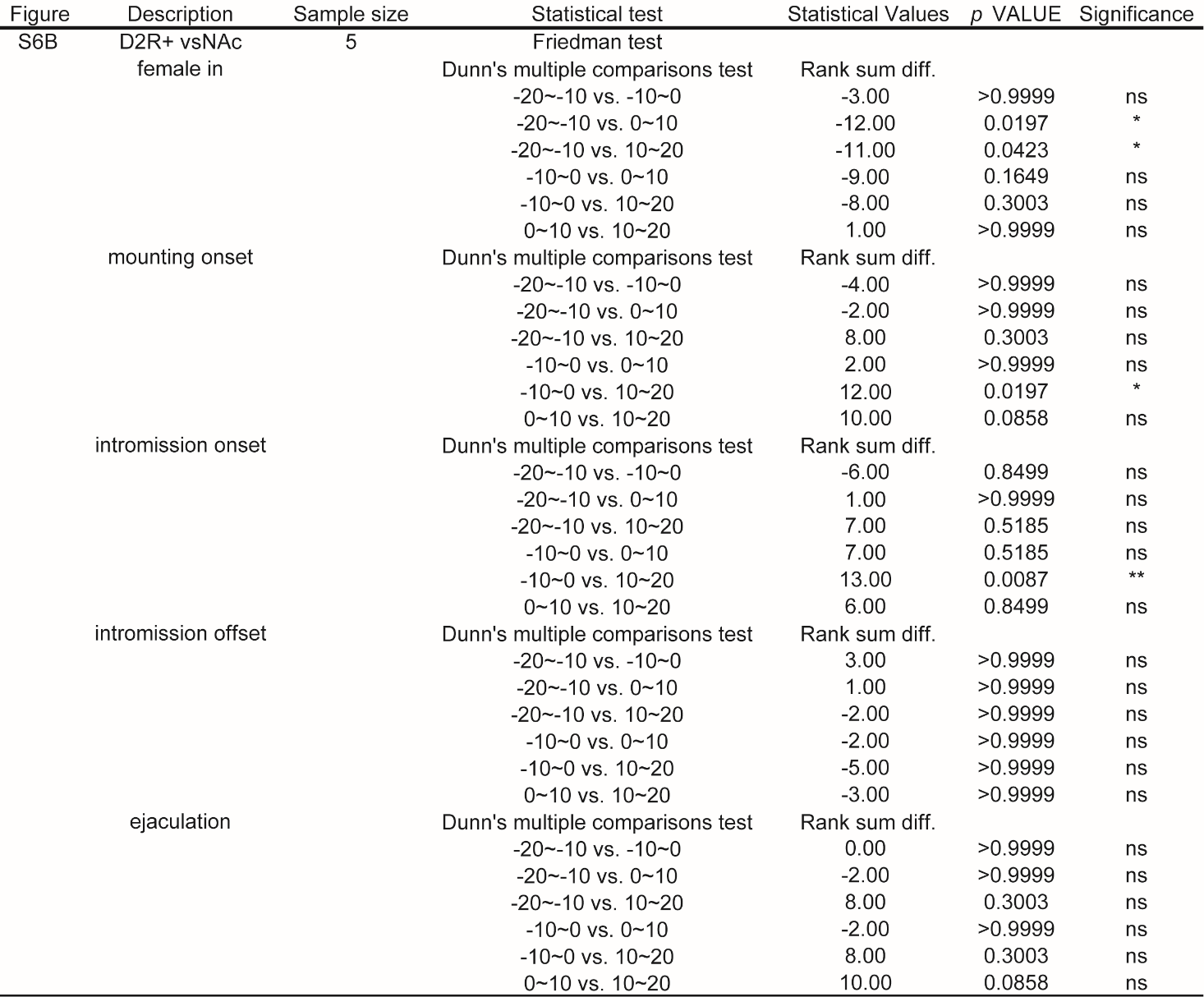

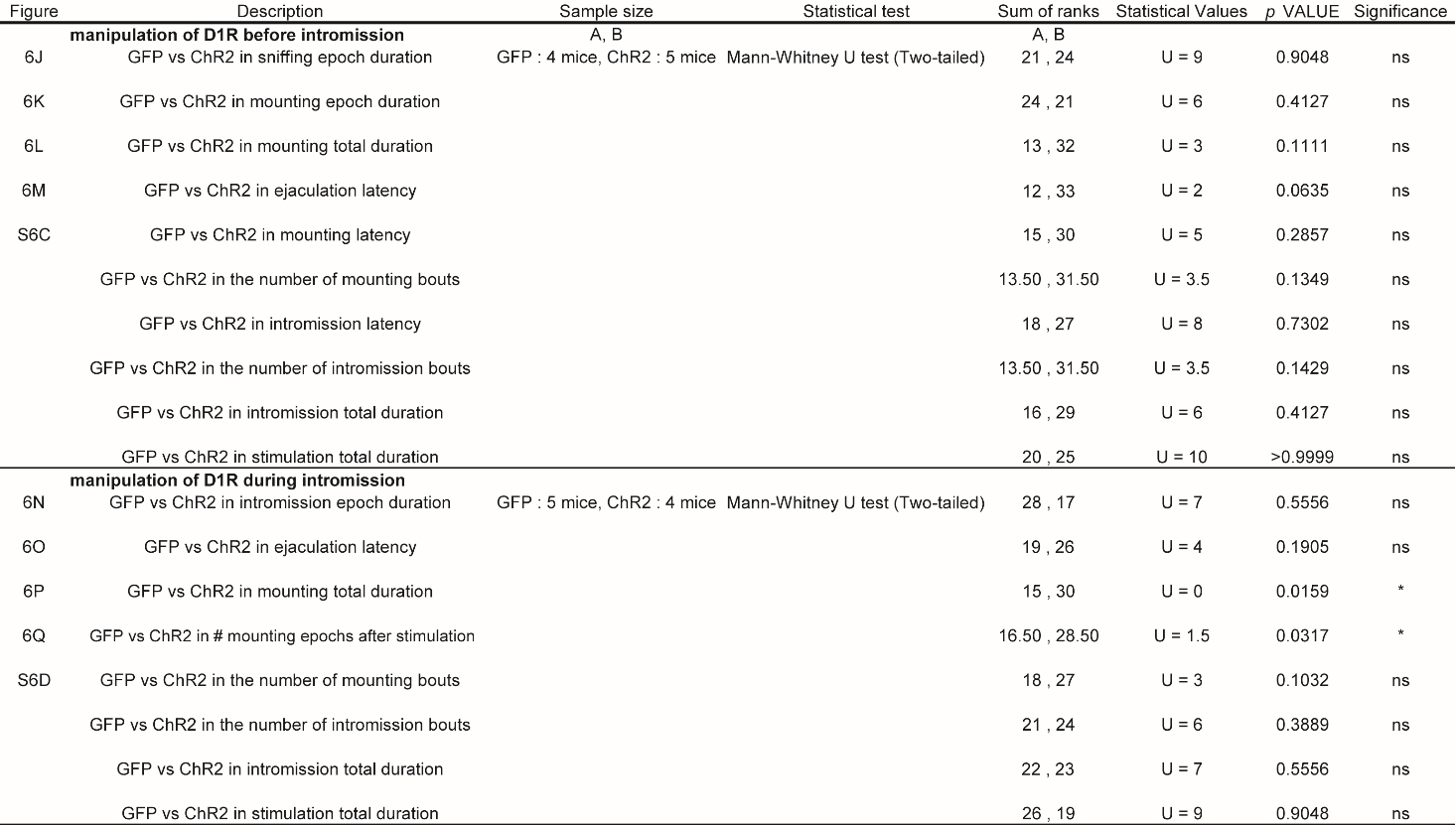

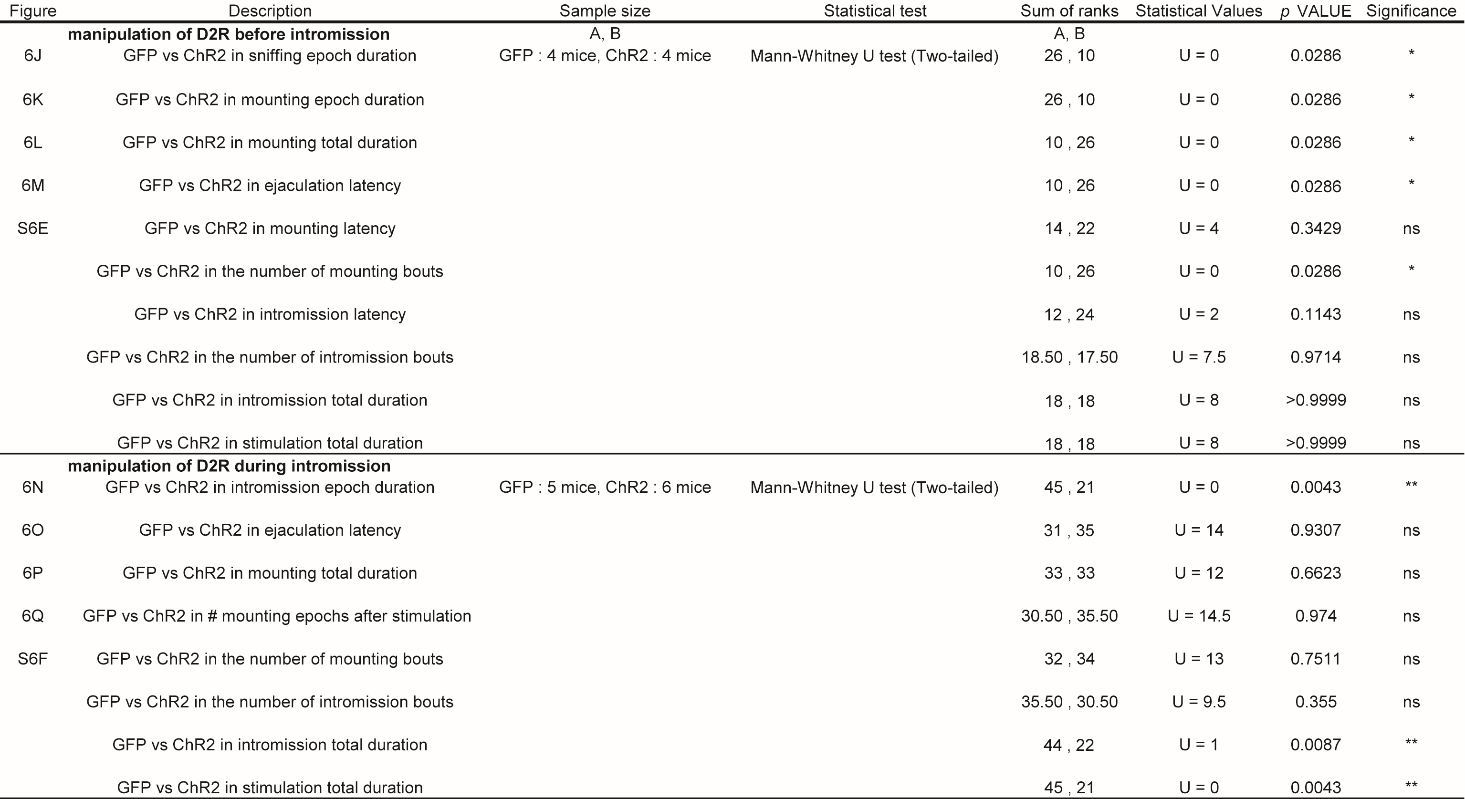

##### Table S22. Analysis of the effects on sexual behaviors following optogenetic activation of D2R^vsNAc^ neurons before and during intromission (related to Figures 6J–6Q, S6E, and S6F)

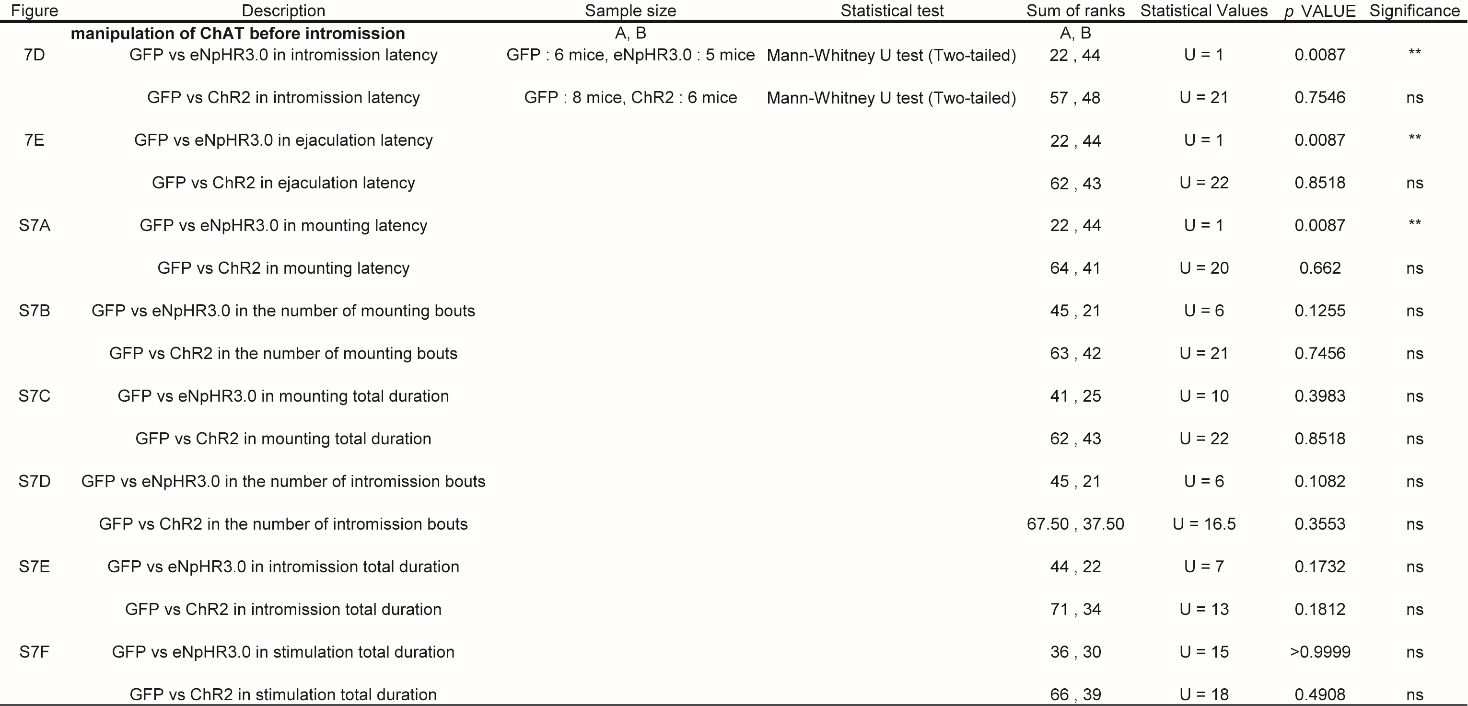

##### Table S23. Analysis of the effects on sexual behaviors following optogenetic activation of ChAT^vsNAc^ neurons before intromission (related to Figures 7D, 7E, S7A–S7F)

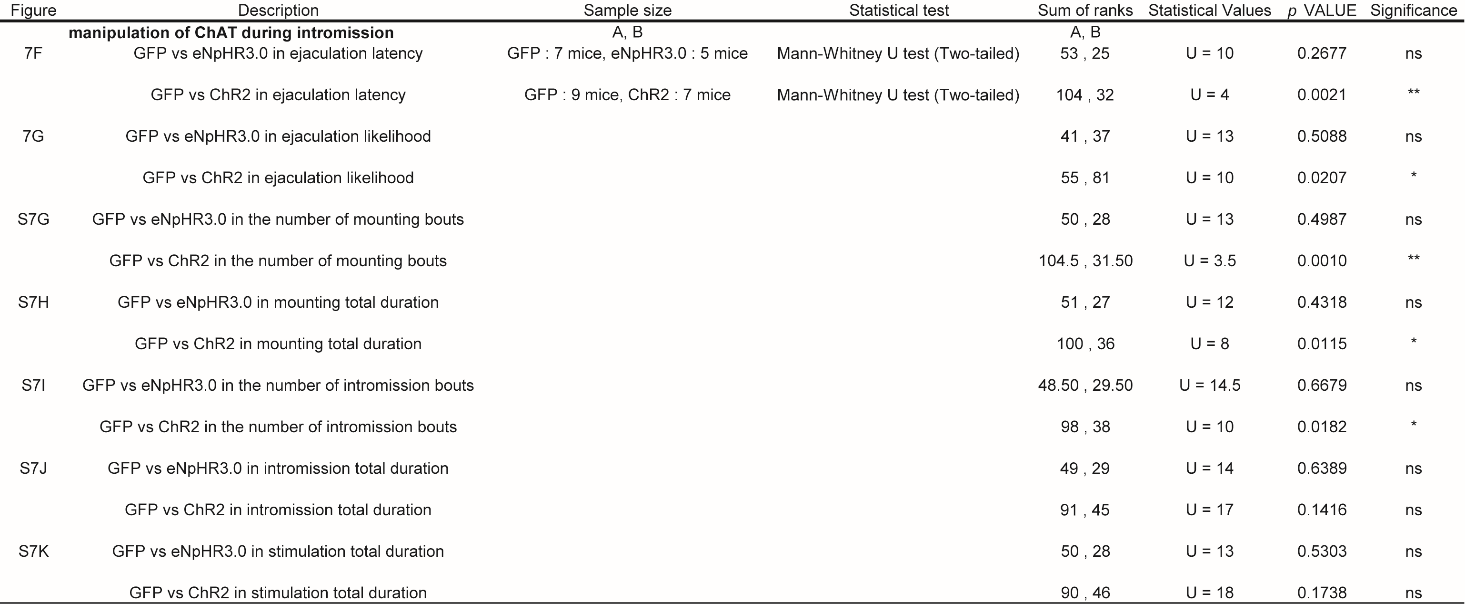

##### Table S24. Analysis of the effects on sexual behaviors following optogenetic activation of ChAT^vsNAc^ neurons during intromission (related to Figures 7F, 7G, S7G–S7K)

##### Table S25. Comparison of the sum of 1.5–2.2Hz of DA rhythms around intromission onset (related to Figure 7L)

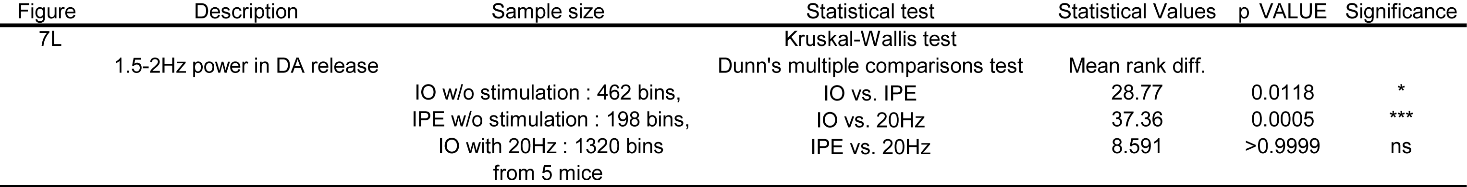

##### Table S26. A list of AAV viruses used in this study.

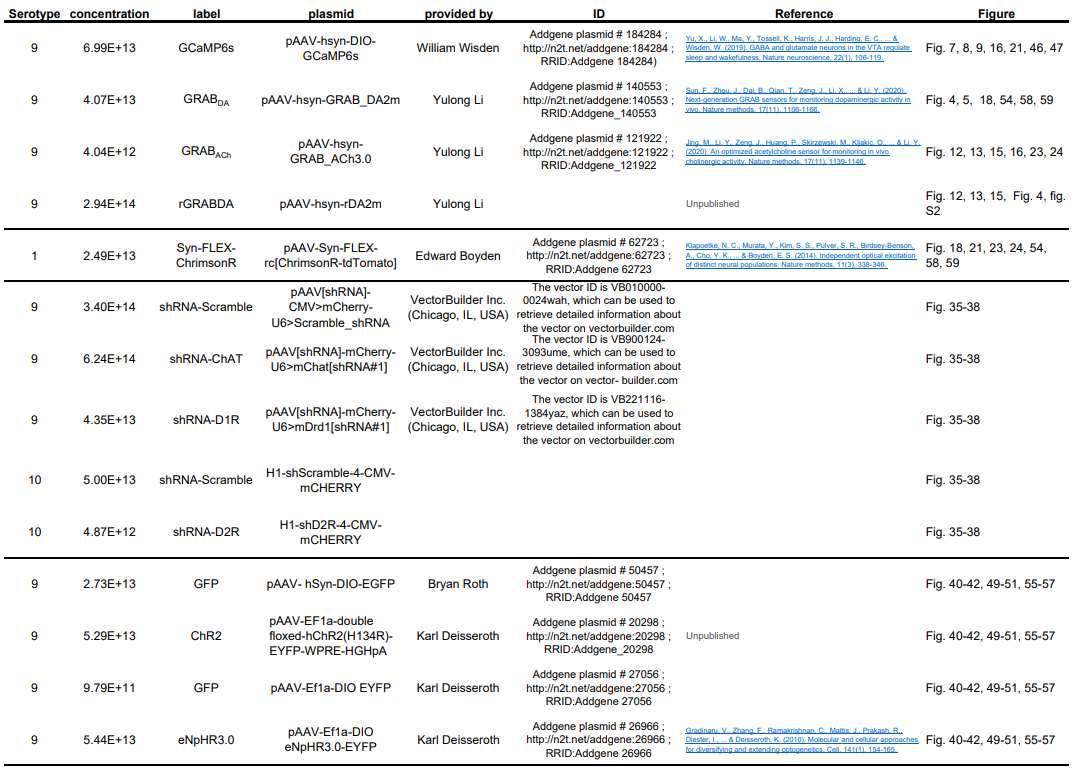

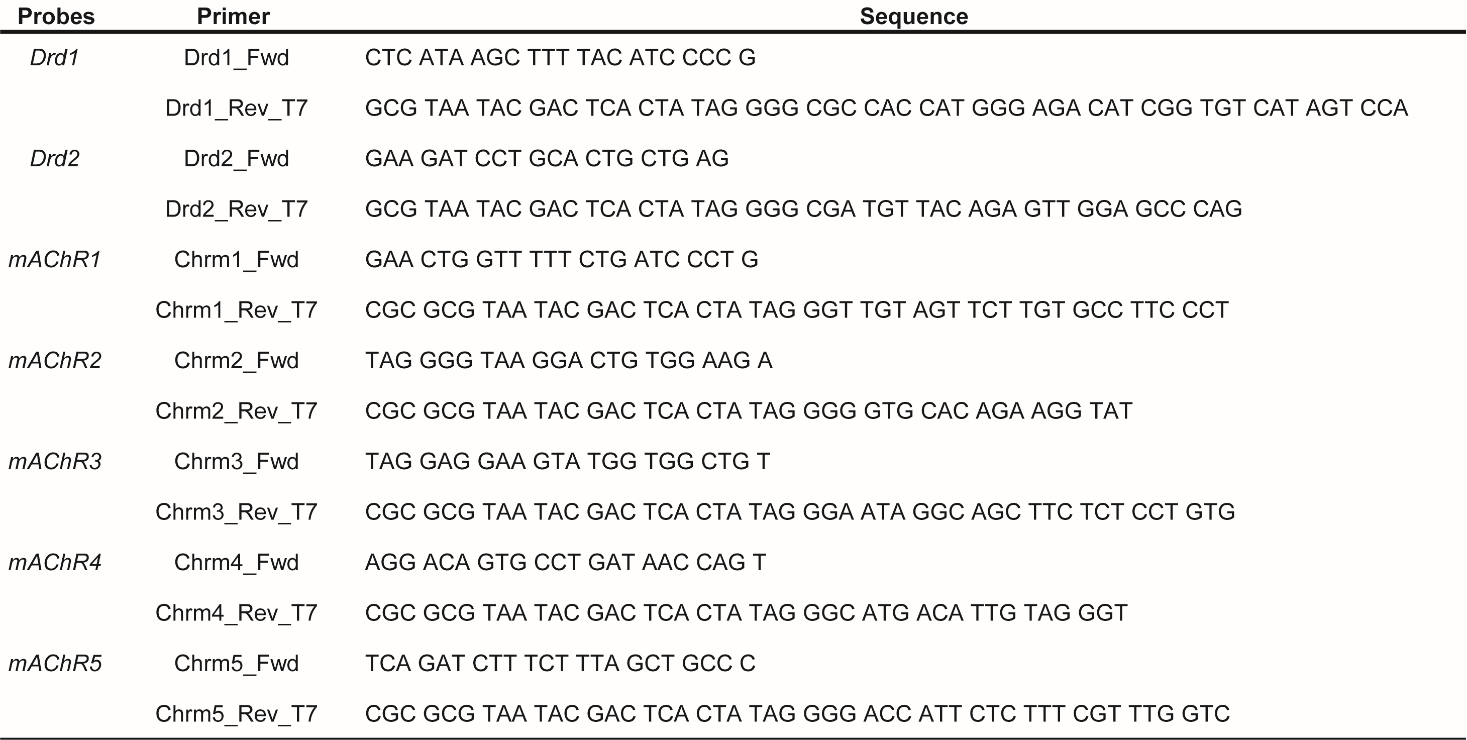

##### Table S27. A list of probes used for in situ hybridization (5`-3`)
